# A General Framework to Interpret Hydrogen-Deuterium Exchange Native Mass Spectrometry of G-Quadruplex DNA

**DOI:** 10.1101/2023.08.24.554600

**Authors:** Eric Largy, Matthieu Ranz, Valérie Gabelica

## Abstract

G-quadruplexes (G4s) are secondary structures formed by guanine-rich oligonucleotides involved in various biological processes. However, characterizing G4s is challenging because of their structural polymorphism. Here, we establish how hydrogen-deuterium exchange native mass spectrometry (HDX/MS) can help to characterize G4 structures and dynamics in solution. We correlated the time range of G4 exchange to the number of guanines involved in inner and outer tetrads. We also established relationships between exchange rates, number of tetrads, bound cations, and stability. The use of HDX/native MS allows for the determination of tetrads formed and assessment of G4 stability at a constant temperature. A key finding is that stable G4s exchange through local fluctuations (EX2 exchange), whereas less stable G4s also undergo exchange through partial or complete unfolding (EX1 exchange). Deconvolution of the bimodal isotope distributions resulting from EX1 exchange provides valuable insight into the kinetics of folding and unfolding processes, and allows one to detect and characterize transiently unfolded intermediates, even if scarcely populated. HDX/native MS thus represents a powerful tool for a more comprehensive exploration of the folding landscapes of G4s.

**TOC graphic:** 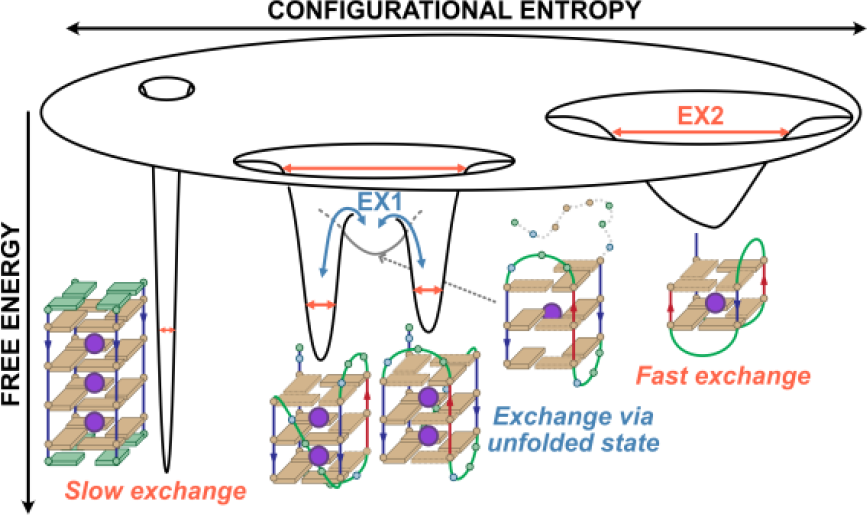

## 1 Introduction

Nucleic acids containing multiple guanine repeats can fold into quadruple-stranded secondary structures called G-quadruplexes (G4s).^1,2^ In G4s, four guanines form G-tetrads (also called G-quartets) through eight hydrogen bonds on both their Watson−Crick and Hoogsteen faces (Figures **1** and S**1**), and at least two G-tetrads stack by coordinating a cation, usually potassium, in the central channel of the structure.^3^ G4s are associated with important biological functions at both the DNA and RNA levels. They may constitute selective therapeutic targets because their structures differ markedly from the canonical double-helix.^4–7^ G4s are also putative drugs themselves, with many aptamers adopting this particular secondary structure.^8,9^ Finally, G4s are increasingly used in nanotechnology applications.^10^

**Figure 1:**
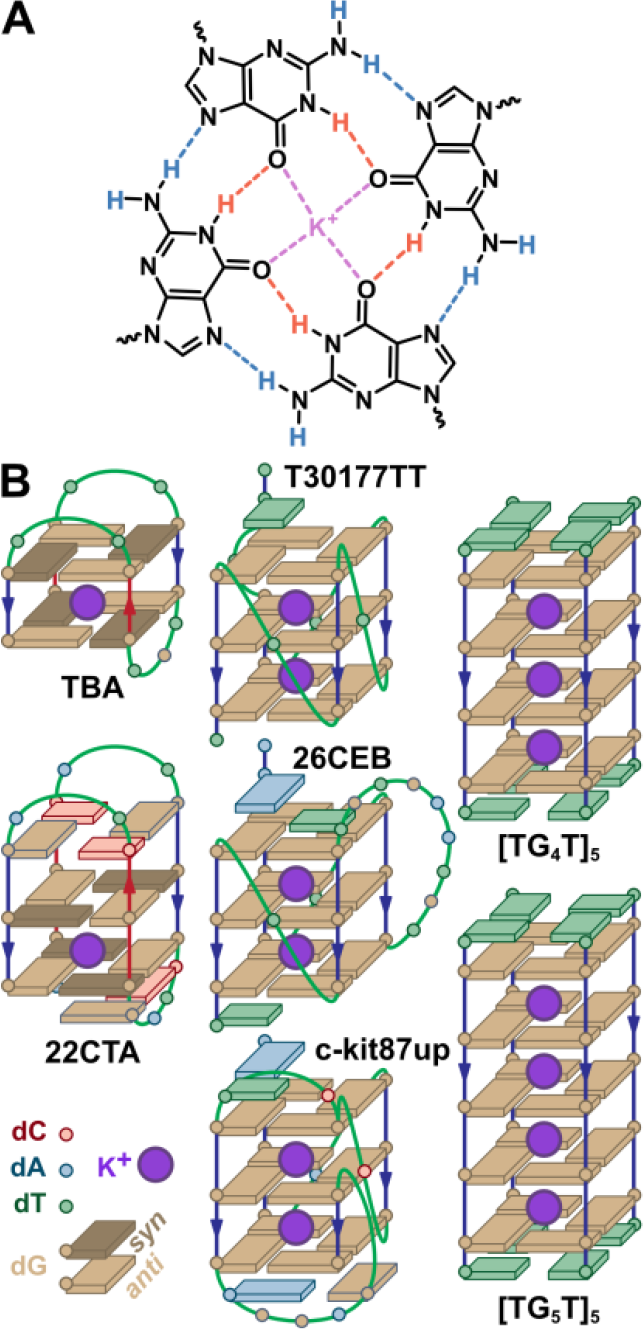
A. Tetrad of guanines, wherein imino (orange) and amino (blue) protons are engaged in H-bonds. B. Simplified schemes of selected G4 structures containing two (left), three (middle), four and five (right) tetrads.

Whatever the application, a bottleneck in the structural characterization comes from the polymorphism of G4s: the number of tetrads, the relative orientation of the strands (parallel, antiparallel, or hybrid), and loop geometries (e.g. lateral, diagonal, propeller) are all heavily dependent on the sequence, but also the nature and concentration of cations, temperature, co-solutes and binders (small molecules, proteins).^3,11–15^ Under given experimental conditions, a single sequence can fold into several conformers in dynamic equilibrium.^15,16^

The characterization of these conformers is challenging. Traditional methods often favor a single conformer (x-ray crystallography) or only provide information about the conformational ensemble (circular dichroism, UV-melting). NMR is most often performed using sequences and experimental conditions tailored to favor a single conformer.^17^ Although these techniques provide invaluable information on nucleic acid structures, complementary techniques are required to study the dynamic conformational ensembles and decipher co-existing topologies.^16^

A method of choice for studying dynamic biomacromolecules, especially proteins, is hydrogen-deuterium exchange coupled to mass spectrometry (HDX/MS).^18–24^ Indeed, the rate of hydrogen exchange is highly dependent on the H-bonding status and solvent accessibility, so measuring these rates by mass spectrometry provides information about the structure and dynamics of the molecule. Following the pioneering work of Linderstrøm-Lang,^25,26^ the exchange has been formally described by a two-step model,^27^ where the sites must first become exchange-competent by opening at the *k*_*op*_ rate before the actual chemical exchange takes place at the *k*_*ch*_ rate (shown in Equation (1) in the D-to-H direction).^18,20,23,24,28–30^ The sites close again, whether or not the exchange has occurred, at the *k*_*cl*_ rate. If the experiment is performed with a large excess of one isotope, the reaction is irreversible.^28^ This model has been proposed to apply to nucleic acids.^31^

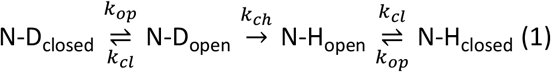

The apparent exchange rate *k*_*HDX*_ is defined in Equation (2). If the favored state in given experimental conditions is native (folded and exchange incompetent), then *k*_*op*_ ≪ *k*_*cl*_ + *k*_*ch*_, leading to the simplified Equation (3).

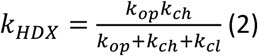

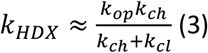

Recent HDX/MS developments have focused heavily on the study of proteins and their complexes.^18–24^ Yet, hydrogen exchange studies to characterize the structural dynamics of nucleic acids were pioneered over five decades ago by Printz & Von Hippel for DNA,^32^ and S.W. & J.J. Englander for RNA,^33^ using the tritium-Sephadex method. Recently, we have shown that HDX coupled to native MS is well suited for the study of structured oligonucleotides.^34^ The proton/deuteron exchange of amino and imino groups is slowed down by H-bond formation between nucleobases and/or screening from the bulk medium. The rate of exchange may therefore reflect the structure of nucleic acids. Furthermore, coupling to native mass spectrometry, a method that ensures that non-covalent complexes are maintained in the gas phase, gives information on individual analytes separated by their mass (binding stoichiometry, binding constant),^35,36^ and is amenable to the measurement of fast-exchange events.^34^ Thus, HDX/native MS might characterize the structural dynamics of individual oligonucleotide conformers and complexes in mixtures. It would be a valuable complement to ^1^H-NMR monitoring of HDX, which allows site-specific measurement of imino (but not amino) protons over a wider range of experimental conditions (buffer, temperature), provided they are resolved and assigned.^37^ However, it suffers from longer dead times (> 2 min), which precludes the measurement of fast-exchanging sites and usually limits the scope of the analysis to internal tetrads.^38–45^

Here, we present the application of HDX/native MS to a set of seventeen G4-forming oligonucleotides of different topologies (Figures **1**B, S**1**) and with a wide range of stabilities (*T*_*m*_ = 28—73°C; Table **1**). In the first section, we systematically describe the exchange behavior of G4s, which will serve as a reference for future investigations, and will allow us to frame the time range of G4 exchange. We then explore the quantitative relationship between this exchange, the number of tetrads and specifically bound cations, and the stability of G4s. Finally, we explore the different exchange scenarios (local fluctuations, large-scale unfolding) and their implications for interpreting G4 dynamics.

**Table 1:**
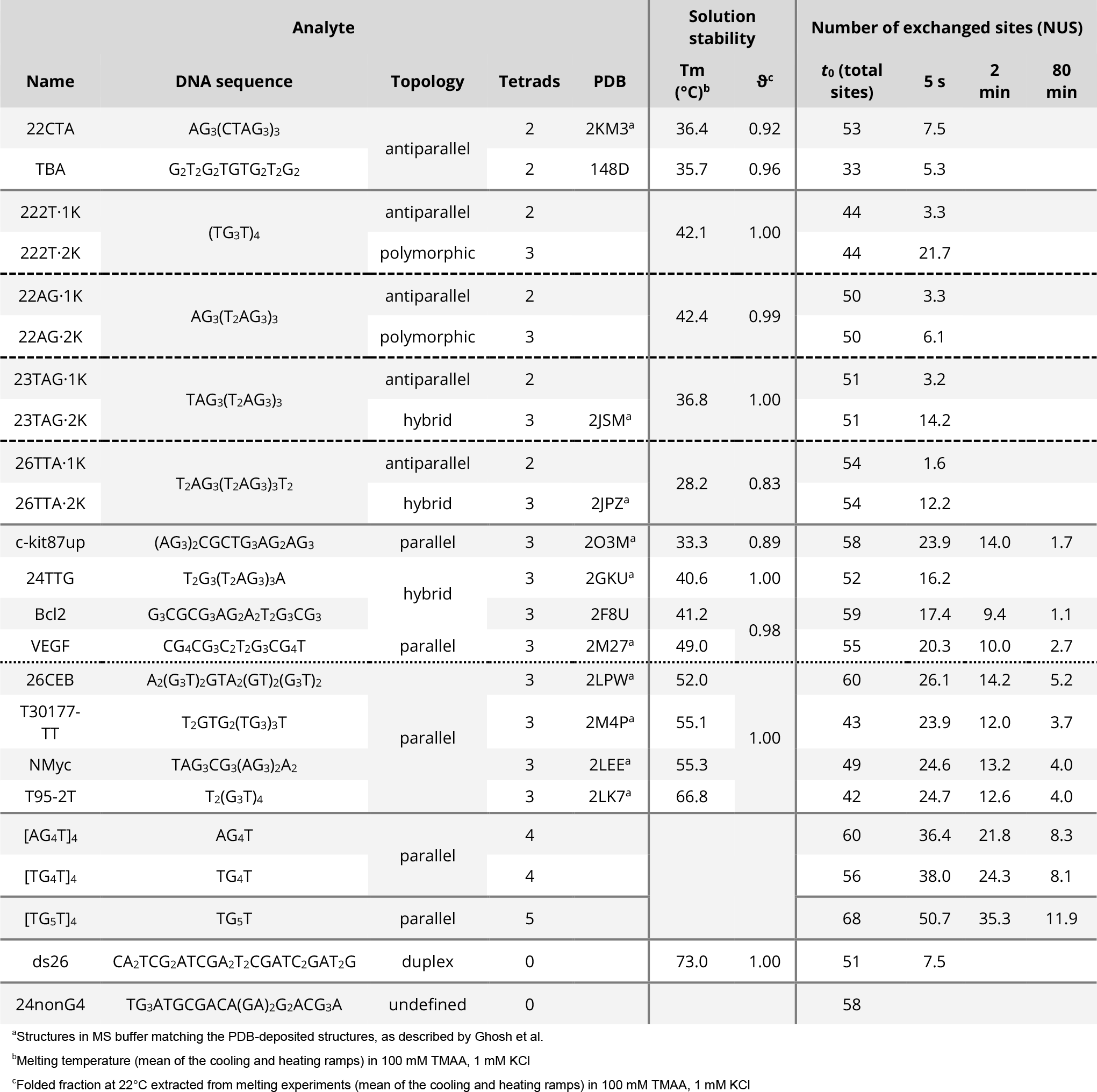
Oligonucleotides analyzed in this study: structure, solution stability, and protection at time points of interest.

## 2. Experimental section

### 2.1. Materials

Oligodeoxynucleotides were purchased from Eurogentec (Seraing, Belgium) or IDT (Leuven, Belgium) in desalted and lyophilized form, and dissolved in either nuclease-free water (Ambion, Life technologies SAS, Saint-Aubin, France) or D_2_O at a target concentration of 1 mM. Their sequences, structures and melting temperatures are summarized in Table **1**. Trimethylammonium acetate (TMAA) (1 M aqueous solution) was purchased from SantaCruz Biotechnology (Santa Cruz, CA, USA), and KCl (99.999% trace metal basis) and D_2_O (99.9% D atom) were obtained from Sigma-Aldrich (Saint-Quentin Fallavier, France).

### 2.2. Methods

Data analysis was performed with R 4.2.1^46^ in RStudio 2023.06.0^47^. UV/visible, UV-melting, circular dichroism (CD), and NMR methods are described in the supporting information. Native MS binding kinetics were performed as previously described.^48^ UV-melting, CD, and HDX/NMR were performed as described in supporting information (Figures S**2**-S**4**).

#### 2.2.1. HDX/MS data acquisition

HDX/MS experiments were performed on a Thermo Orbitrap Exactive mass spectrometer equipped with a daily calibrated electrospray source (ESI) operating in negative mode. Typical acquisition parameters were: 300—3000 m/z scan range, 50 000-resolution setting, 2.95 kV spray voltage, sheath gas: 55, aux gas: 5, heater temperature: 25°C, 200°C capillary temperature, capillary voltage: -45 V, tube lens voltage: -165 V, skimmer voltage: -34 V. The softness of these tuning parameters was assessed by infusing a 10 µM solution of the [G_4_T_4_G_4_]_2_ G4 in 100 mM ammonium acetate and adjusted to be soft enough to avoid in-source back-exchange and maintain non-covalent interactions (Figure S**5**).^34,49^

The deuterated (pre-exchange) samples were prepared by diluting the oligonucleotides to 50 μM DNA (from stock solutions in D_2_O) in 100 mM TMAA, 1 mM KCl (unless otherwise stated), 90% D_2_O (pH = 7.0). The samples were then annealed at 85°C for 5 min (to ensure statistical exchange of all sites to 90% D), and stored at 4°C for at least a night before use. The diluent solutions used to induce D-to-H exchange consisted of 100 mM TMAA, 1 mM KCl (or a concentration matching that of the sample) in H_2_O (pH = 7.0). Experiments were performed at 22°C by mixing one volume (70 µL) of deuterated sample (90% D) with nine volumes (630 µL) of diluent (0% D). Mixing was performed either by continuous-flow mixing, to access exchange events ranging from approximately one second to five minutes, or manually to monitor longer exchange times (i.e minutes to days).^34^ In manual mixing experiments, the sample and diluent were thoroughly mixed by vortexing, and the resulting solution (9% D) was transferred to a 500-μL glass syringe, which was installed on a syringe pump operated at 5 μL/min and coupled to MS. In continuous-flow experiments, the deuterated sample and the diluent solutions were pumped at a 1:9 ratio towards a PEEK high pressure static mixing tee, equipped with a UHMWPE frit (IDEX Health & Science, Oak Harbor, WA, USA). The resulting mixed solution was brought to the MS source through PEEK tubing of varying volumes to produce different mixing times. In all cases, the 9%-D reference samples were prepared from the post-exchange samples, which were further incubated at 85°C for 5 minutes to ensure statistical exchange.

#### 2.2.2. HDX data processing

Raw data files in .raw format were converted to mzML using the MSConvert utility (Version: 3.0.20175)^50^ and then semi-automatically processed using OligoR, a software developed in-house and described elsewhere.^51^ Briefly, OligoR allows one to convert centroid *m*/*z* to number of unexchanged sites *NUS*. For continuous-flow experiments, raw HDX/MS data were analyzed by first selecting the isotope distributions of species of interest for each discrete exchange time point *t* and combining scans over one minute of acquisition (Figures S**6**—S**8** and S**9**A). The exchange time *t* = *V*/*F* itself is defined by the time spent by the analytes in the mixing fluidics (from the mixing tee to the MS source) of volume *V*, at a flow rate *F*. In real-time experiments, the scans were processed individually.

At the intact level, the apparent exchange kinetics of an oligonucleotide is the sum of the exchange kinetics of each site it contains, which are expected to exchange at their own rate. Depending on the exchange time range, some sites may already be fully exchanged while others are barely exchanged. Nevertheless, some sites may have rates that are similar enough to be grouped together in the same exponential decay process, thus reducing the number of terms needed to fit Equation (4), where *NUS*_∞_ is the offset, *N*_*i*_ the number of exchanging sites, *k*_*i*_ is the exchange rate and *j* the number of groups of sites with similar rates (Figure S**9**B,C). Half lives were derived from the decay rate using Equation (5).

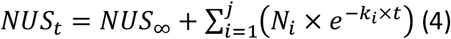

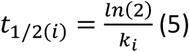

Here, we systematically evaluated which of a single (*j* = 1) or double exponential (*j* = 2) was better to obtain a satisfactory fit, using the Akaike’s Information Criterion with small-sample correction (AICc)^52,53^ and the Bayesian Information Criterion (BIC). The fitting procedure itself was performed with multiple initialization values to improve its robustness, using the *nls*.*multstart* R package.^54^ The HDX/MS exchange plots and their non-linear fitting are shown in Figure S**10**.

#### 2.2.3. Deconvolution of multimodal isotopic distributions

Above, we track the exchange of oligonucleotides by monitoring their *m*/*z* centroid. This is sufficient for monomodal isotopic distributions (‘EX2 exchange’). In some cases, however, bimodal isotopic populations have been observed. Detection and deconvolution of these isotopic populations may indicate exchange *via* unfolded states (‘EX1 exchange’) and/or several conformers of the same mass but with different protection factors. Therefore, systematic least-squares optimization of the experimental isotopic distribution was performed with one (monomodal) or two (bimodal) theoretically generated isotopic distributions using OligoR (Figure S**11**).^51^

From the deconvoluted bimodal distributions, the *NUS*_*j*_(*t*) values at a time *t* for a population *j* (1 or 2) can be determined using Equation (6), where *DC*_*j*_(*t*) is the optimized deuterium content for the corresponding time and population, and *nX* is the number of exchangeable sites.

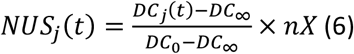

The ‘pure’ EX2 contributions of each population are its *NUS* as a function of the exchange time, and can be fitted using Equation (4), similar to the non-deconvoluted data. The EX1 contribution is visualized by plotting the abundance of each population as a function of the exchange time. The apparent rate of this contribution is obtained using Equation (7), where *ab*_∞_ is the final abundance and *ab*_*Δ*_ is the amplitude of the change in abundance.

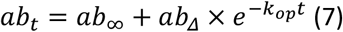

## 3. Results

### 3.1. Relationship between apparent exchange rates, G4 structures, and stability

#### 3.1.1. Overview of exchange behaviors

H/D exchange rates are very sensitive to protein folding. However, little is known about the influence of DNA secondary structures on these rates. Therefore, we first investigated the apparent exchange behavior of a set of model oligonucleotides forming G4s of different stabilities and topologies (parallel, antiparallel and hybrid, with two to five tetrads, see Table **1**). Among these sequences, ten adopt the same conformation as those deposited in the PDB when prepared in our native MS buffer.^13^ Bcl-2 and TBA are expected to fold into their PDB-deposited conformations based on CD, and both have a small unfolded population. TG4T, AG4T, and TG5T associate to form parallel tetramolecular G4s, with small amounts of unfolded single strands being detected.

Some also fold into 2-tetrad G4 conformers (23TAG, 26TTA, 22AG, 222T),^15,48^ and others contain small unfolded populations (c-kit87up, VEGF).^13^ Hereafter, conformers formed by a single sequence but containing different numbers of tetrads are distinguished by the number of tetrad-bound potassium cations, since a *n*-tetrad G4 typically coordinates *n-1* cations.^1,3,13,48,55^ For example, 22AG·1K^+^ is a 2-tetrad conformer, while 22AG·2K^+^ adopts a 3-tetrad conformation. The exchange kinetics of such species have been measured independently from the data of a single experiment because they are mass-resolved under native conditions.

HDX/NMR was used to establish a global time frame of G4 exchange for individual imino protons, considering that amino protons were not monitored. The experiment was performed on T30177TT in the buffer used for HDX/MS (Figure S**4**). The exchange of imino protons from external tetrads is completed within the first five minutes. Imino protons from internal tetrads, however, are exchanged over the course of five hours, with no significant differences between guanines. MS monitoring of H/D exchange was therefore performed by continuous-flow mixing to access exchange events ranging from approximately a second to five minutes, and real-time mixing to probe longer exchange times (*i*.*e*., minutes to days with periodic sampling), yielding exchange plots as shown in Figure S**9**B (all data shown in Figure S**10**).

##### 3.1.1.1. Unfolded strands

All unfolded single strands, i.e. 24nonG4 and non-hybridized AG4T, TG4T, and TG5T, are completely exchanged within the dead time of approximately two seconds (*NUS* = 0 at all time points; gray plots in Figure S**10**A) due to the lack of stable H-bonds.^34^ This is in contrast to the structured oligonucleotides (G4s and duplex), which are all protected to some extent (Table **1**). For the sake of standardization, the *NUS* values after 5 seconds of exchange, *NUS*_5*s*_, will be used as a reference for short-exchange times.

##### 3.1.1.2. Duplex DNA

ds26 is a hairpin that exchanges in about two minutes, with most of the exchange occurring in the first minute. It has 51 exchangeable sites, many of which are involved in base-pairing (25 assuming the formation of a 4-nucleotide loop). However, the equivalent of less than 8 sites are unexchanged after five seconds of mixing (*NUS*_5*s*_ = 7.5), and exchange at approximately the same rate thereafter (Figures S**12**-S**14** and tables S**1**-S**3**). Exchange by unfolding of a significant portion of ds26 is unlikely because of its high stability (*Tm* = 73°*C, ΔG*° = −28 kJ/mol), and the isotopic distributions are monomodal, so the exchange must occur through structural breathing.^56^

##### 3.1.1.3. Two-tetrad G4s

The two-tetrad G4s (TBA, 22CTA, 22AG·1K^+^, 23TAG·1K^+^, 26TTA·1K^+^, 222T·1K^+^) have *NUS*_5*s*_ values between 1.6 and 7.5 (Figure S**10**A, Table **1**). At this short exchange time, it is likely that only H-bonded imino sites from tetrads remain (partially) unexchanged. Other sites that are permanently exposed and thus exchange-competent (e.g., non H-bonded loop, flanking residues) can exchange in the folded state of the oligonucleotide at an apparent rate close to that of its chemical rate *k*_*ch*_, following Equation (8) where the probability *β* of being locally exposed is close to or equal to 1.^57^ The exchange of these sites in not visible in the time range we studied (red zones in Figure **2**A).

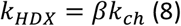

**Figure 2:**
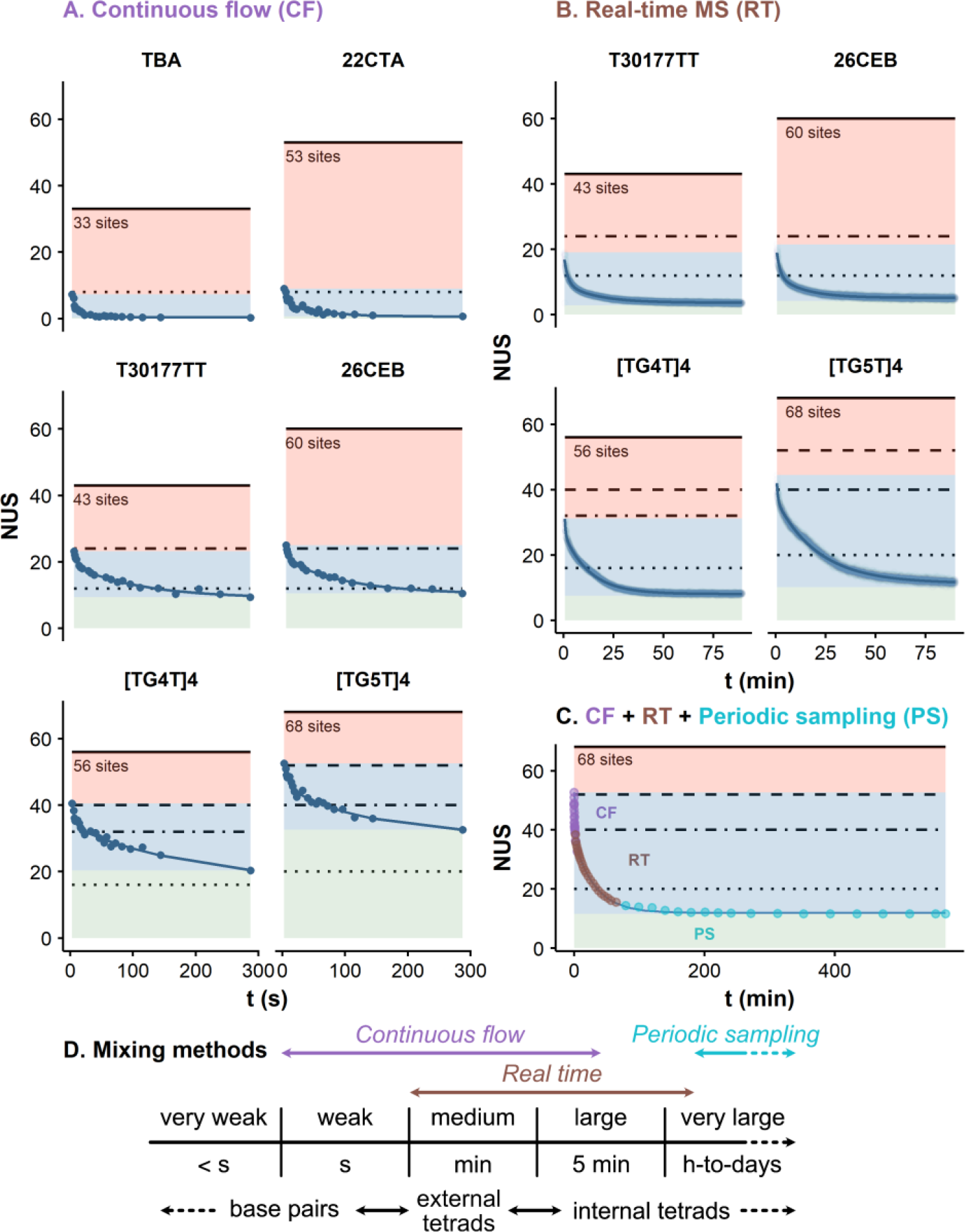
Exchange plots of selected species acquired by A. Continuous-flow/MS (dead time ∼ 1 s), B. Manual mixing with real-time MS monitoring (dead time ∼ 1 min) and C. A combination of A, B and periodic sampling for [TG_5_T]_4_. Plots are divided in three areas along the y-axis: Red: dead time exchange, invisible with the method, blue: visible exchange events, green: sites that did not exchange yet. Four horizontal lines indicate noteworthy site numbers: total number of imino protons in tetrads (4 per tetrads; dotted), total number of H-bonds in tetrads (8 per tetrads; dot-dashed), total number of H-bonds in tetrads plus non-H-bonded amino sites in internal tetrads (12n-8 per n tetrads; dashed), and total number of exchangeable sites (solid). The blue lines are the results of non-linear fitting with Equation (4) with j = 2, except for panel C where j = 3. D. Different protection levels found in G4s can be probed by different HDX/MS methods.

The protected sites of the 2-tetrad G4s are completely exchanged within two minutes. 22CTA, which contains two G·C base-pairs forming a pseudo third tetrad,^58^ appears slightly more protected at short exchange times (*NUS*_5*s*_ = 7.5 vs 5.3—1.6). Nonlinear fitting analysis using Equation (4) shows that 2-tetrad G4s contain two sets of 2 to 4 sites, whose exchange rates differ by about an order of magnitude (*k*_1_ = 0.2—1.3 s^-1^, *k*_2_ = 0.01—0.09 s^-1^; Figures S**12**-S**14** and Tables S**1**-S**3**).

##### 3.1.1.4. Three-tetrad G4s

All three-tetrad structures, except 22AG·2K^+^, have *NUS*_5*s*_ above 12 (Table **1**), which corresponds to their number of imino sites in the tetrads (dotted lines in Figures **2**A and S**15**). Four species (T30177TT, NMyc, T95-2T, 26CEB; *T*_*m*_ > 50°C) have at least 24 *NUS*_5*s*_ (dot-dashed lines in Figures **2**A and S**15**). This suggests that either all imino and amino H-bonds in tetrads are protected, or that a few non-tetrad base-paired nucleotides contribute. Other less stable analytes (*T*_*m*_ < 50°C) have lower *NUS*_5*s*_, ranging from 16 to 22 for 24TTG, Bcl2, VEGF, and 222T. Oligonucleotides that also form significant amounts of 2-tetrad conformers or even unfolded strands, such as 22AG·2K^+^, 26TTA·2K^+^, and 23TAG·2K^+^, are outliers that behave midway between the 2- and 3-tetrad classes (*NUS*_5*s*_ = 6—14), even though the 2·K^+^ stoichiometry complexes were specifically analyzed. This can be explained by the contributions from 2-tetrad species binding two cations (including one non-specifically) and/or exchange *via* unfolding (see section 3.2). However, in contrast to the 2-tetrads G4s, the exchange is not completed within the first 300 s for any of the 3-tetrad G4s: at the largest time point measured, these structures have between 5.7 and 12.6 *NUS*, indicating that one or more tetrads remain partially or completely deuterated. This is consistent with the decrease in imino proton exchange rate expected in internal tetrads:^59^ HDX/NMR shows that the imino sites from the internal tetrad of T30177TT remain largely unexchanged during the first five minutes, while the external tetrads exchange more extensively (Figure S**4**). The outliers 22AG·2K^+^, 26TTA·2K^+^, and 23TAG·2K^+^ are however almost completely exchanged within the first five minutes, with *NUS* of 2.3 or less.

Nonlinear fitting analysis shows that at least two sets of sites can be distinguished in 3-tetrad G4s, with exchange rates differing by an order of magnitude (Figures S**12** and S**13**, and Tables S**1**—S**3**). The first set exchanges at the second time scale (*k*_1_ = 0.1—1.4 s^-1^; *t*_1/2_ = 1—7 s; Figure S**14**), similar to sites from 2-tetrad G4s, and is therefore consistent with the exchange of weakly-protected sites. The second set contains slower-exchanging sites (*k*_2_ = 0.007—0.022 s^-1^; *t*_1/2_ = 32—106 s), keeping in mind that some even slower exchanging sites may contribute very little to the total exchange in this time frame (green area in Figure **2**A). The number of sites in this set ranges from 7 to 11 (except for the 22AG·2K^+^ outlier: 4), suggesting similarly-behaving, slow-exchanging sites in 3-tetrad G4s.

Seven 3-tetrad G4s from the panel were subjected to longer exchange times, using the real-time MS setup (Figures **2**B, S**10**B and S**16**). The most stable ones (T95-2T, T30177TT, NMyc, and 26CEB; *T*_*m*_ = 52.0— 66.8°C) still have many protected sites after two minutes (*NUS*_2*min*_ = 12.0—14.2; Table **1**) and plateau after 50 minutes to *NUS* values of around 4 (*NUS*_50*min*_ = 3.9—5.3). These remaining sites are likely the well-protected imino protons from internal tetrads that exchange slowly over the next 30 minutes (*NUS*_80*min*_ = 3.7—5.2). Their lifetime is significantly longer than the time range studied, as evidenced by HDX/NMR (Figure S**4**). Conversely, the less stable VEGF, Bcl2 and c-kit87up G4s (*T*_*m*_ = 49, 41 and 33°C, respectively) exchange more extensively (*NUS*_80*min*_ < 2.7). Two sets of sites can be distinguished by this manual mixing approach (Tables S**4**—S**6**). The fastest decay events are remarkably similar to the slowest measured by continuous-flow (average *k*_1_ = 0.01 s^-1^, *t*_1/2_ = 49 s), consistent with the overlap in incubation time between these approaches. The slower regime has rate constants that are at least an order of magnitude smaller (average *k*_2_ ≈ 0.001 s^-1^, *t*_1/2_ = 11 min).

##### 3.1.1.5. Four and five-tetrad G4s

The four-tetrad [TG_4_T]_4_ and [AG_4_T]_4_ have similar exchange plots, with *NUS*_5*s*_ of 38 and 36, respectively, which is more than the 32 imino and amino H-bonds in their four tetrads (Table **1**). Similarly, [TG_5_T]_4_ contains 40 H-bonds in its five tetrads, but has a *NUS*_5*s*_ of 51. In these three cases, there are few opportunities for H-bonds to form outside the tetrads. One hypothesis is that both amino deuterons of each guanine buried in internal tetrads are protected from exchange, although only one is H-bonded at any time. According to this hypothesis, the number of significantly protected sites of an *n*-tetrad G4 (containing *n* − 2 internal tetrads) would be: *n* × 8 + (*n* − 2) × 4 = 12*n* − 8. This fits well for five- and four-tetrad species (dashed lines in Figures **2**A and S**15**), but not for G4s containing fewer tetrads. Many sites are still protected after 250 s (*NUS*_250*s*_ = 21, 20, and 33), well above their number of imino protons. The non-linear fit yields two distinct decay regimes, both of which are globally slower than those of the 3-tetrad G4s (*k*_1_ = 0.077—0.167 s^-1^ and *k*_2_ = 0.005—0.007 s^-1^ vs. averages of 0.316 and 0.012 s^-1^ for 3-tetrads species). After 80 minutes, the *NUS* values plateau remarkably close to the number of internal imino protons, e.g. 11.9 for [TG_5_T]_4_ (12 imino sites), 8.1 and 8.3 for [TG_4_T]_4_ and [AG_4_T]_4_ (8 imino sites), respectively. These buried sites are particularly well protected from exchange: they barely exchange at all over ten hours (see the periodic data sampling in Figure **2**C).

#### 3.1.2. Relation between number of tetrads, specific cation binding, and exchange

Sequences with many sites in the loops and flanking sequences contain more exchangeable sites, but these additional sites are rapidly exchanged when not involved in H-bonds (red areas in Figures **2**A, S**15** and S**16**). This is the so-called folded state exchange (*β* ≈ 1 in Equation (8)). For example, TBA has fewer sites than the other 2-tetrad G4s (33 vs. 44—54), which have enough guanines to form 3-tetrad conformers, but all have similar exchange plots (compare TBA and 22CTA in Figure **2**A). Another striking example is the contrast between the 26-mer 26CEB and the 19-mer T30177TT, which have very similar exchange behaviors despite having very different numbers of sites (60 vs. 43). Thus, the number of non-exchanged sites does not correlate with the total number of exchangeable sites (i.e. the primary structure), but with the number of protected sites, and thus with the secondary structure.

Besides the very weakly protected sites (*t*_1/2_ in the subsecond range; invisible with our setup), four sets of sites of increasing protection can be distinguished: weak (second range), medium (minute range), large (ten-minute range), and very large (hours to days) (Figure **2**C)). Two-tetrad G4s have at most medium-protected sites, whereas 3-tetrad G4s have sites with large protections. Very large protections are observed for internal imino sites ((*n* − 2) × 4 for *n* tetrads) within the more stable 3-tetrad species, and very kinetically stable 4- and 5-tetrad G4s. Overall, this results in a visible correlation between the number of tetrads and the number of non-exchanged sites at each time point (Figure **3**A,B). Indeed, the relationship between *NUS*_*t*_ and the number of tetrads for *t* seems linear in the range 5—250 s and 2—80 min (limited to *t* < 100 s for 2-tetrad species because they are completely exchanged thereafter), as shown in Figure **3**C,D.

**Figure 3:**
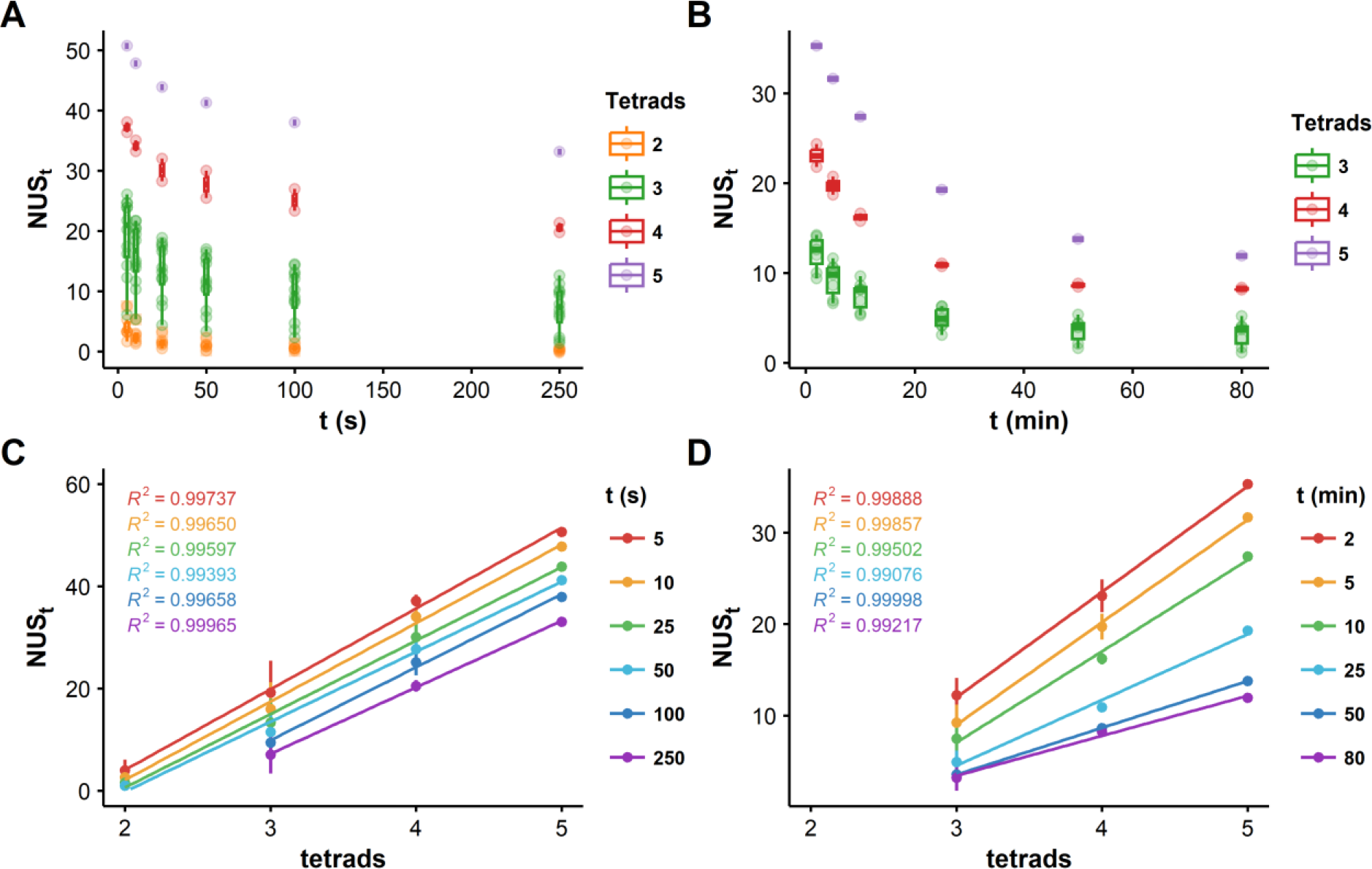
Relationship between the exchange kinetics and the number of tetrads of G4s measured by continuous-flow (left) and real-time (right) HDX/MS. A,B: Distributions of exchange plots grouped by number of tetrads; C,D: mean NUS as a function of the number of tetrads, at different exchange times. The values for 2-tetrad G4 exchange are excluded at 100 s and above as the exchange is already complete for most or all species. The straight lines are the result of linear fitting.

These relatively strong trends are useful to confirm G4 formation, and to assign a number of tetrads to the formed conformers. We have deposited our panel data online to help with this assignment.^60^ The number of tetrads can be confirmed by determining the number of specifically bound cations, since *n* tetrads specifically bind *n* − 1 cations.^55^ However the total signal contains specifically and non-specifically bound cations. Non-specific cations do not alter HDX rates,^34^ and therefore HDX/MS is extremely useful to provide the number of specific cation(s) simply by comparing the exchange kinetics of different adduct stoichiometries (see example in Supporting Information, Figure S**17**).

Despite the correlation between the number of tetrads and the *NUS* values, there are significant differences within the tetrad groups, suggesting that the HDX kinetics depend on specific topologies and not just on a number of tetrads. This is particularly evident within the three-tetrad group, for which we tested a wider range of structures often containing loop- and flanking-base pairing.

#### 3.1.2. Relation between G4 stability and exchange rates and mechanism

Above, we have hinted at the influence of the thermal stability of G4s and their exchange rates. A more systematic analysis shows a correlation between the *T*_*m*_ and the *NUS* values at each time point (*NUS* at 5 s and 50 min are shown in Figure **4**A,B, but similar trends can be observed at other exchange times in Figures S**18** and S**19**). Protection from exchange is also correlated to apparent 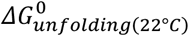, which better reflects the isothermal stability of G4s than *T*_*m*_ does (Figures **4**C,D, S**20** and S**21**). However, both *T*_*m*_ and 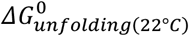 are derived from melting experiments assuming only two states (folded and unfolded), whereas global HDX rates reflect a combination of many local unfolding events.^61^ The rates of these events differ depending on their location, since we observed order-of-magnitude differences between the *k*_*HDX*_ of external and internal tetrads. Nucleotide-level HDX/MS data will be required to access local 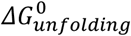 values.

**Figure 4:**
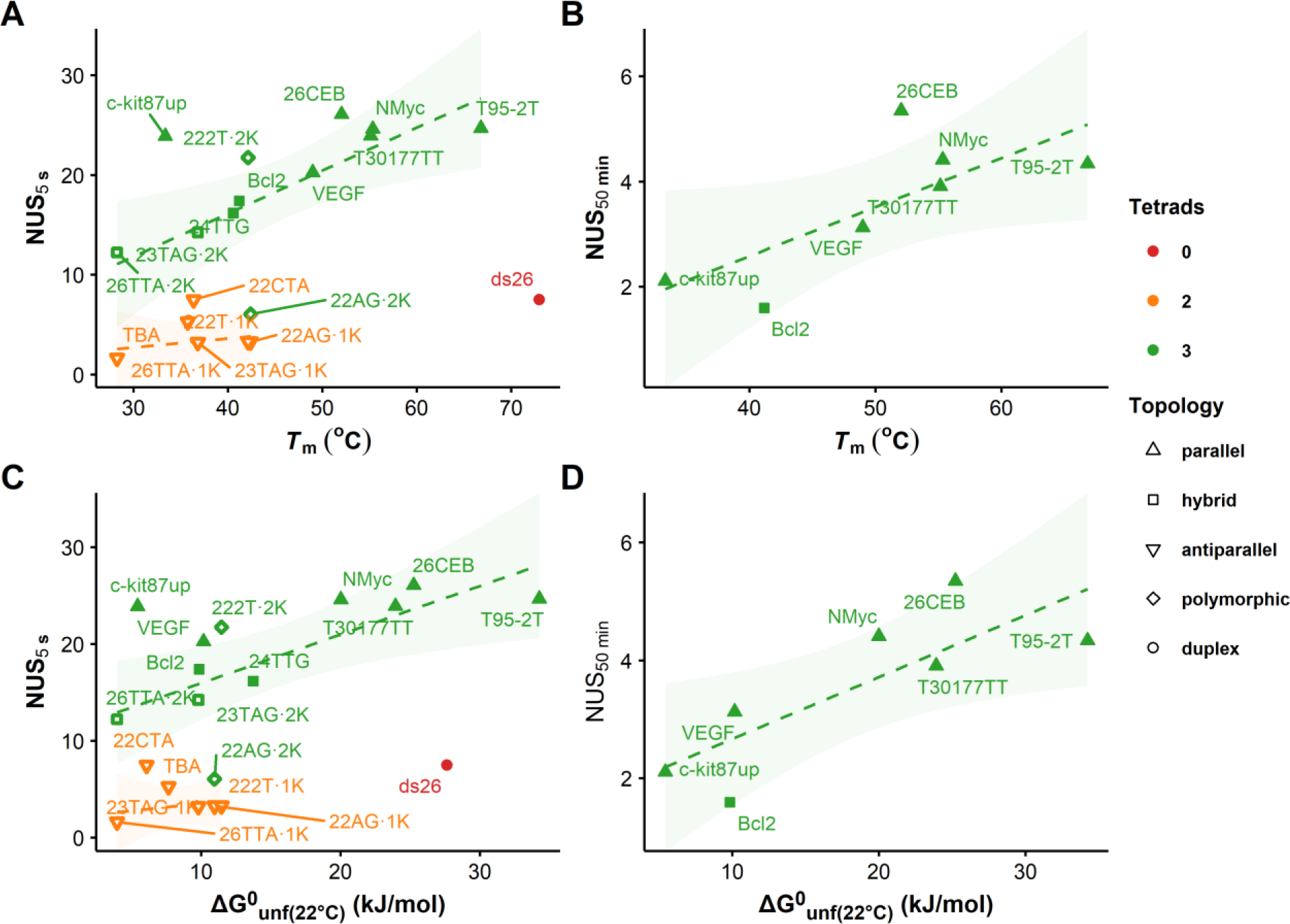
Relationship between the exchange kinetics and the stability of G4s measured by continuous-flow (left; 5 s exchange) and real-time (right; 50 min exchange) HDX/MS. A,B: Relation between T_m_ and NUS_t_. The straight line and 95% confidence interval (colored ribbons) are the result of linear fitting. Open symbols indicate an EX1 contribution, while filled symbols correspond to pure EX2 exchange. C,D: Same as A,B but between the apparent free energy of unfolding at 22°C and NUS_t_.

The correlation does not extend across different secondary structures: the duplex-forming ds26 exchanges much faster than T95-2T despite having a larger *T*_*m*_ and reasonably close 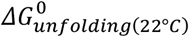. Similarly, 2-tetrad G4s have similar exchange rates (and number of sites) to ds26 despite having much lower *T*_*m*_ values and free energies. This shows the ability of tetrads to protect nucleobases from exchange much more efficiently than base pairs undergoing breathing motions.^32,56^ Even within G4s, two- and three-tetrad species follow different trends due to the increased protection of internal tetrads.

Within the 3-tetrad G4 group, the correlation remains limited but can highlight certain G4 characteristics. c-kit87up is quite unstable (*T*_*m*_ = 33°C, θ = 0.89) but displays a high protection at short incubation times (*NUS*_5*s*_ = 24), comparable to those of the more stable 26CEB, T95-2T, T30177TT and NMyc. However, at longer exchange times, the number of non-exchanged sites decreases to levels similar to less stable G4s such as BCl2 (*NUS*_80*min*_ = 1.7). c-kit87up contains a 5-nucleotide snapback loop forming two A·G base pairs (Figures **1** and S**1**). These base pairs may provide additional H-bonding sites and further protection to the 3’-end tetrad on which they are stacked, explaining the high initial *NUS* levels. However, this protection is likely to be weaker than that of stable internal tetrads, hence the extensive exchange at long incubation times. This example highlights the importance of selecting a wide range of exchange times in order to sample different types of exchange events, and thus different structural characteristics.^62^

The correlation between solution stability and exchange rates also exists for a given conformer in different conditions. A reference sample of T95-2T containing 1.0 mM KCl (*T*_*m*_ = 66.8°C) was compared with samples containing 0.75, 0.50, 0.25, and 0.10 mM KCl in which the G4 is less stable (*T*_*m*_ = 65.1—52.2°C) but remains fully folded at room temperature (θ_22°*C*_ = 1.0) and retains a parallel topology (Figure S**22**). A decrease in protection occurs at lower [K^+^], especially at longer incubation times (Figure **5**A). A fairly linear trend of *NUS*_*t*_ = *f*([*K*^+^]) is observed between 1.00 and 0.25 mM KCl, where the magnitude of the decrease is limited but significant (*ΔNUS* = 1.30; Figure **5**B). The decrease of [K^+^] (at least in this range), and thus the thermodynamic stability of T95-2T, leads to a significant increase in the local dynamics of the G4 without visibly altering its secondary structure. Consistently, the apparent protection against exchange scales linearly with the free energy of unfolding (Figure **5**C). Strikingly, the decrease in protection is much steeper and results in a bimodal isotopic distribution when [KCl] is reduced to 0.10 mM. This suggests a concerted exchange of several sites following the unfolding of G4, which will be further explored below. This decrease in protection also leads to a loss of linearity between *NUS*_*t*_ and both [*K*^+^] and 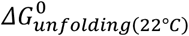 (dotted lines in Figure **5**B,C).

**Figure 5:**
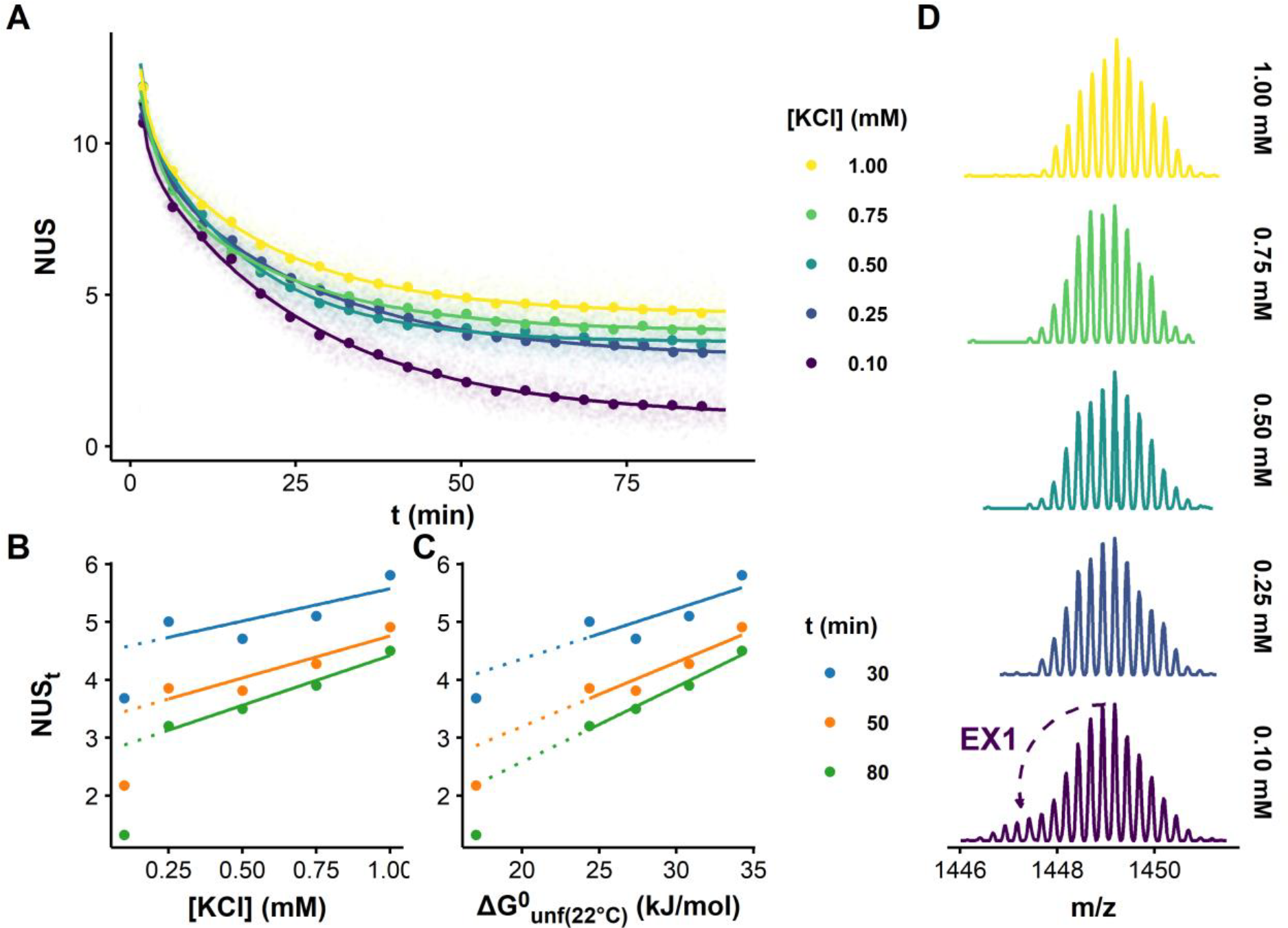
Influence of the stability of T95-2T on its exchange rate and mechanism: A. HDX/MS kinetics in presence of 0.10—1.00 mM KCl (4-species binding 2 K^+^). The transparent points correspond to single scans, the non-transparent points to moving averages (width: 50 scans, every 250 scans). The fitting procedure was performed on the single-scan data using Equation (4), where j = 2. B. Change in NUS_t_ (t = 30, 50 and 80 min.) as a function of [KCl], showing a linear decrease of the protection with [KCl] in the 0.25—1.00 mM range, and a large loss of protection at 0.10 mM, and C. Change in NUS_t_ as a function of the apparent free energy of unfolding, showing a linear decrease of the protection with the decrease in stability. D. HDX/MS spectra highlighting the shift from a pure EX2 to a mixed EX1/EX2 exchange mechanism upon decreasing [KCl] (13 scans combined across 12.6 s centered at 6.2 min; 4-species binding 2 K^+^).

### 3.2. Beyond averages: exploring exchange mechanisms

#### 3.2.1. Stable G4s exchange without visiting unfolded states

The exchange of stable three-tetrad G4s is characterized by mass spectra with single isotopic populations that gradually shift to lower *m*/*z* over time (Figures S**23**—S**30**). They are all very unlikely to unfold in the assay conditions (*T*_*m*_ ≥ 52.0°C) and the exchange likely occurs by discrete exchange events occurring via local fluctuations only. In this exchange regime, termed EX2, many opening events must occur before the exchange itself takes place because *k*_*cl*_ ≫ *k*_*ch*_. The apparent rate defined in Equation (3) can therefore be approximated by Equation (9), where *K*_*op*_ is the equilibrium constant of the opening reaction. In this regime, the free energy of the unfolding reaction depends on the local structural stability, and can be related to *k*_*HDX*_ (Equation (10)).^23,63^

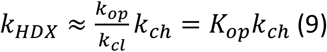

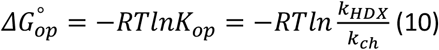

Tetramolecular G4s formed by TG_4_T, AG_4_T, and TG_5_T exchange *via* EX2 although they are technically in equilibrium with single strands that exchange very quickly. In reality, these G4s have very slow hybridization rates and very large half-lives, indicating no contribution from the single strand (further discussed in supporting information; Figure S**27**).^64,65^ The control duplex-forming sequence ds26 also exchanges by an EX2 mechanism (Figure S**31**). Although the shortest exchange time points (< 5 s) yield distributions slightly broader than what is theoretically possible with a single population, the use of a second population did not yield a statistically better fit. This is consistent with an exchange occurring via double-helix breathing motions.^32,56^ It is possible that stable G4s may exchange through similar opening of the outer tetrad, although this phenomenon is still not well understood.^59^ This would also be consistent with the strong protection provided by internal tetrads, for which breathing motions are likely to be restricted. The exchange of these buried sites may require larger unfolding events, which are unlikely to happen in stable G4s and are therefore kinetically limiting.

#### 3.2.2. Mixtures of conformers can produce multimodal isotopic distributions

Oligonucleotides forming less stable G4s exhibit a markedly different behavior; exchange data often present bimodal isotopic distributions that reflect either:

i. The presence of two different topologies that exchange at significantly different rates. The abundance of their respective isotopic population should remain unchanged.
ii. The exchange of multiple sites via an unfolded state, known as the EX1 regime. The low-exchange isotopic population converts to the high-exchange population with increasing mixing time, leading to concerted changes in isotopic population abundances (Figure S**11**). This is discussed in section 3.2.3.

The 3-tetrad G4 formed by VEGF (*T*_*m*_ = 49°C) exhibits a bimodal distribution in the first few minutes of exchange (Figure **6**A,B and S**32**). The relative abundances of the low-exchange and high-exchange isotopic populations during this time frame remain constant (Figure **6**C). This suggests that this G4 does not visit a long-lived unfolded state under these experimental conditions. The low-*m*/*z* distribution is most likely a contribution of small amounts of unfolded species non-specifically binding two K^+^, or two-tetrad G4 binding non-specifically binding one K^+^. This also means that the equilibration between the 3- and 2-tetrad species (likely requiring complete unfolding) is slow compared to the HDX time frame.

**Figure 6:**
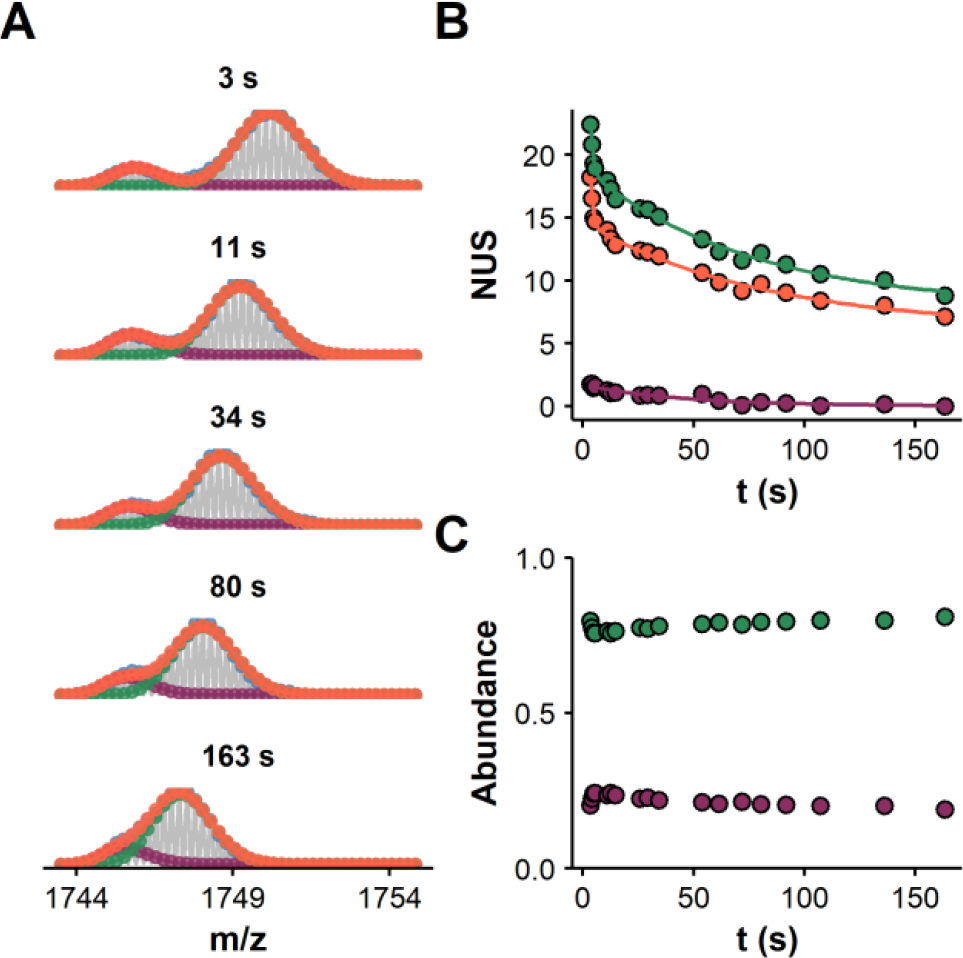
Bimodal HDX data modeling of the exchange data of VEGF (continuous-flow mixing, initial DC = 90%, final DC = 9%). A. The experimental data is shown in grey, the peak picking in blue, the overall fit in orange, and individual population in green and purple (z = 4-, K = 2, time labeled in s above panels). B. Exchange kinetics of the overall (orange) and deconvoluted distributions (green and purple). C. Relative abundance of the deconvoluted distributions as a function of the exchange time.

The bimodal isotopic distributions of c-kit87 and Bcl2 (Figures S**33** and S**34**) are also likely to be largely caused by the presence of unprotected unfolded strands that bind potassium non-specifically, given the presence of 0 and 1 K^+^ species (Figure S**35**). Very slow EX1 contributions cannot be completely ruled out given the slight changes in isotope population abundances, but it would contribute very little to the total exchange. 24TTG is another example of possibly “contaminated” EX2 kinetics, characterized by a very small and constant contribution of a low-mass population (∼3%; Figure S**36**). In contrast, similar but less stable human telomeric sequences (22AG, 23TAG, 26TTA) exchange partially by unfolding (see section 3.2.3).

In native MS, different intramolecular conformers, and their unfolded counterpart, can have identical *m*/*z* if their total number of bound cations (specific or non-specific) is the same. Native HDX/MS allows us to resolve these species by their exchange protection, and to deconvolute the spectra to extract the exchange kinetics of the pure species. The exchange behavior discussed here indicates that the conformer interconversion is slower than the exchange time range.

#### 3.2.3. Low stability G4s can exchange via partially unfolded states

The EX1 exchange regime requires the unfolded regions of the biomacromolecule to close at a much slower rate than chemical exchange. A single visit in the ‘open’ configuration is then sufficient for complete labeling. With *k*_*cl*_ ≪ *k*_*ch*_, Equation (3) becomes Equation (11), where *k*_*op*_ is the opening reaction rate constant. Typically, this results in a bimodal distribution in which the abundance of the highly-exchanged population increases at the expense of the weakly-exchanged population as a function of time (Figure S**11**). It could be seen as an isotopic stamp on molecules as they visit an open conformation. Even if they close again, the previous opening translates in a highly exchanged population that reveals the extent and rate of opening.

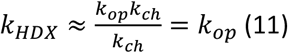

##### 3.2.3.1. The model case of 23TAG

Figure **7** shows the example of 23TAG (global *T*_*m*_ = 37°C) folding in 1 mM KCl solutions into hybrid 3-tetrad G4s binding two potassium cations (23TAG·2K^+^), in mixture with small amounts of 2-tetrad antiparallel G4 (23TAG·1K^+^), and even smaller amounts of unfolded strands (23TAG·0K^+^; Figure S**8**).^13,48^ For 23TAG·2K^+^, two distributions are required to adequately model all time points (Figure S**37**). At longer exchange times, the distribution *appears* to be monomodal, but is too broad to account for a single population; thus, modeling with two populations is necessary. Moreover, the abundances of the isotopic populations change over time: an increase of the low-mass, highly exchanged population, and a decrease of the low-exchange counterpart (Figure **7**C). Thus, the isotopic distribution of 23TAG•2K^+^ is generated by the cooperative exchange of multiple sites via unfolded states.

**Figure 7:**
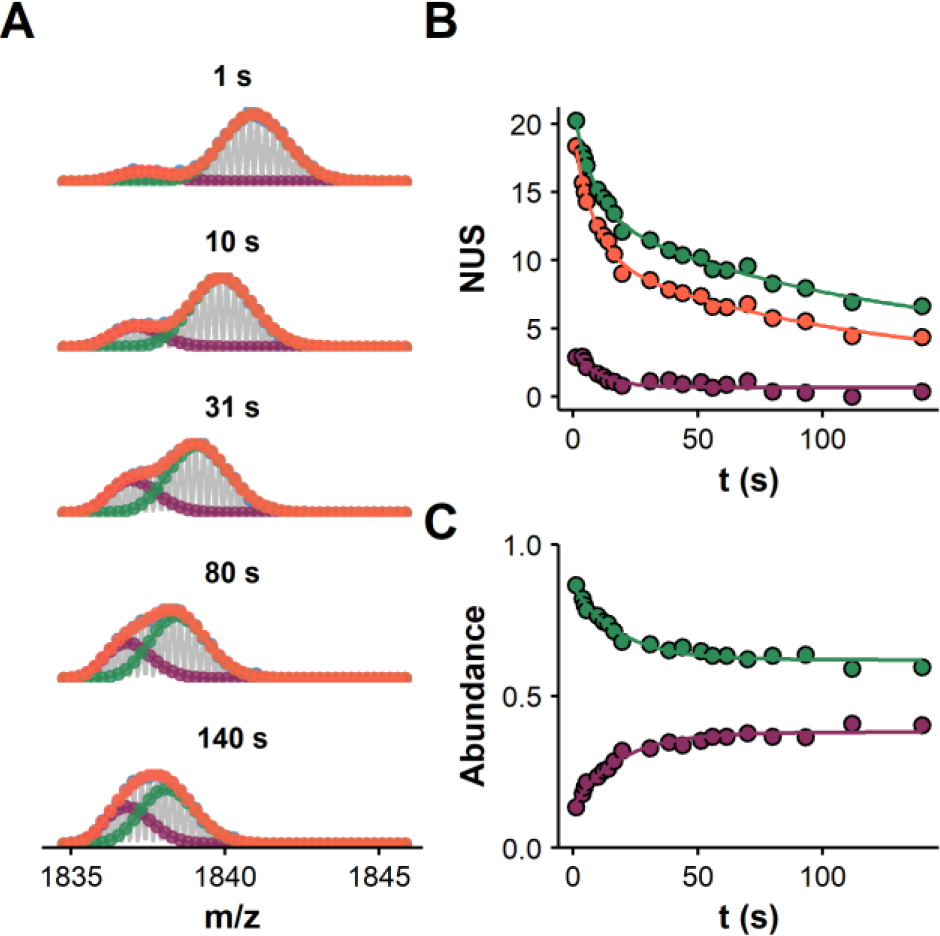
Bimodal HDX data modeling of the exchange data of 23TAG (continuous-flow mixing, initial DC = 90%, final DC = 9%). A. The experimental data is shown in grey, the peak picking in blue, the overall fit in orange, and individual population in green and purple (z = 4-, K = 2, time labeled in s above panels). B. Exchange kinetics of the overall (orange) and deconvoluted distributions (green and purple). C. Relative abundance of the deconvoluted distributions as a function of the exchange time.

Non-linear fitting of the abundance data as a function of time yields an apparent EX1 contribution rate 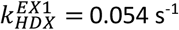 (Table S**7**), which reflects an opening rate *k*_*op*_ (Equation (11)). Despite this relatively fast unfolding (*t*_1/2_ = 12.8 s), only small amounts of 23TAG•0K^+^ species are detected (Figure S**8**). 23TAG·2K^+^ also undergoes HDX *via* the EX2 mechanism, as evidenced by the exponential decay of the *NUS*_*t*_ for both isotopic populations (Figure **7**B). The high rate of EX2 exchange relative to EX1 results in the entire exchange being completed before all 23TAG•2K^+^ complexes have had a chance to visit the ‘open’ conformation, resulting in the incomplete EX1 exchange in Figure **7**C. G4s continue to unfold/refold and eventually all will have visited an unfolded state at least once, but unfolding events are no longer detectable due to the complete exchange of all sites. The mass-resolved 23TAG·1K^+^ also exchanges quickly, owing to its only two tetrads, by both EX1 and EX2 (Figure S**38**). The extensive exchange of the low-mass population in a short exchange time also points to significant unfolding events.

We will now discuss the nature of these partially unfolded states in the light of the G-quadruplex unfolding landscapes. Several teams have evidenced the formation of distinct 3-tetrad hybrid G4s but also of 2-tetrad antiparallel structures, via different intermediates.^66–72^ In this context, our team had monitored the binding kinetics of potassium to 23TAG by native MS (with non-specific cation adduct subtraction).^48^ We proposed that the folding pathway from the unfolded strands (‘*0*’) to the preferred 2K^+^-binding 3-tetrad hybrid conformers (‘*2a*’ and ‘*2b*’) involves a 1K^+^-binding intermediate ‘*1a*’ (*e.g*., G-triplexes), while the rapidly forming ‘*1b*’ ensemble is off-pathway (Figure **8**B). We reproduced here these experiments (Figure **8**A) and dynamic fitting of the binding kinetics with a model reflecting the interplay of these five ensembles yielded the folding and unfolding rate constants (Figure **8**C).

**Figure 8:**
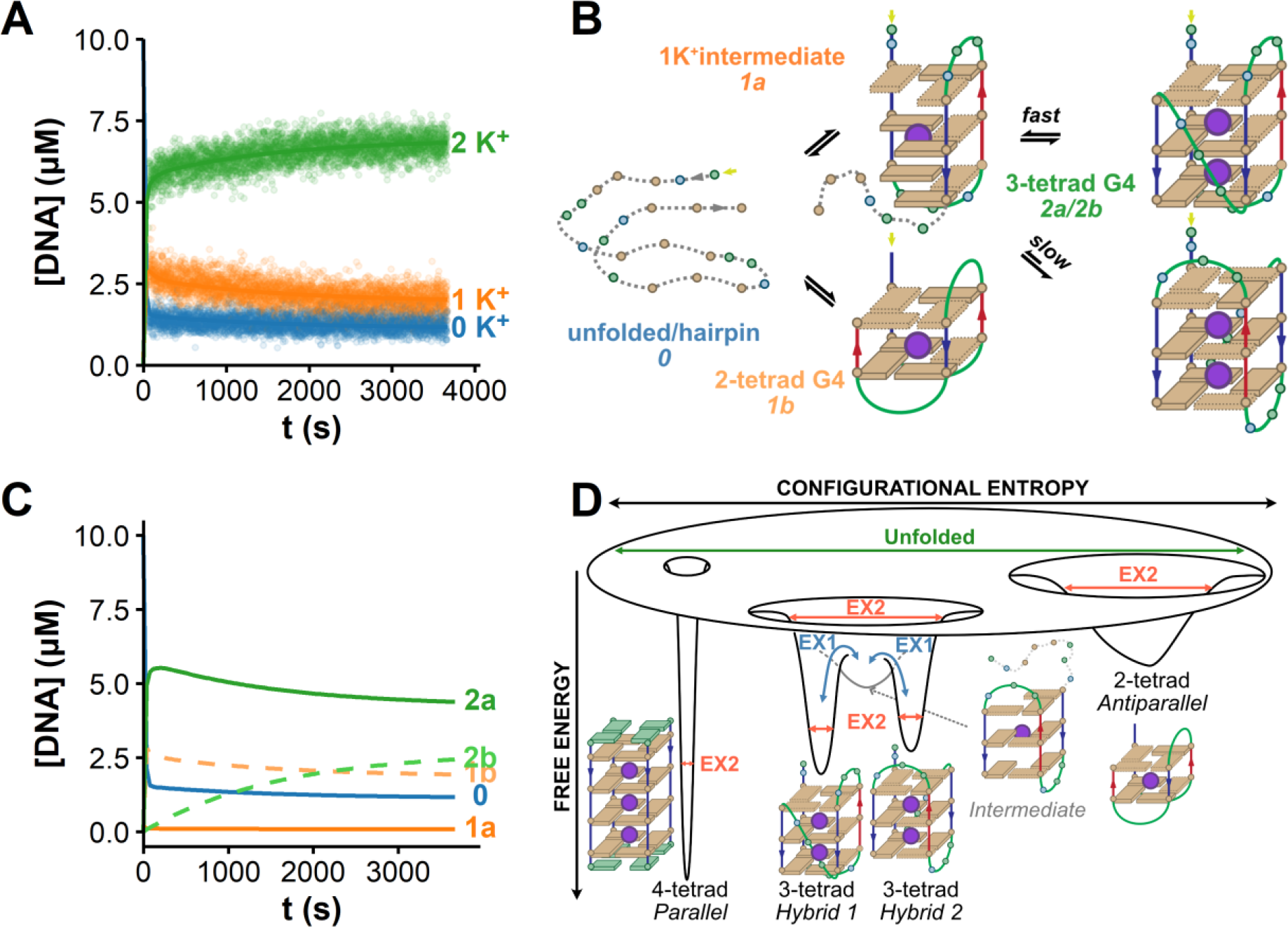
Folding and unfolding of 23TAG. A. Binding kinetics of K^+^ (1 mM) and 23TAG (10 µM) in 100 mM TMAA (pH 7.0). Lines are the result of dynamic fitting using the folding pathway in panel B. C. Simulation of concentration of individual ensembles from the optimized rate constants. D. Relation between the energy landscape of G4s and the exchange mechanism. Increasing number of tetrads give deeper and narrower funnels; the narrower the funnel, the slower the EX2 exchange. EX1 requires to visit other states that can be folding intermediates (e.g. G-triplex) between different topologies.

Interestingly, the rate of unfolding from *2a* to the intermediate *1a* is close to that of the HDX opening rate (*k*_2*a*→1*a*_ = 0.032 s^-1^ vs. 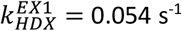). The other conformer (*2b*) unfolds (and folds) much slower and gradually replaces *2a*.^48,72^ It is therefore likely that *2b* contributes significantly less than *2a* to the EX1 exchange, limiting the proportion of overall 3-tetrad G4 visiting the less-folded state on the time scale of the experiment (Figure **7**C). The 0 → 1*a* reaction is much slower (*k*_0→1*a*_ = 5.6 × 10^−5^ μM^-1^ s^-1^) than 1*a* → 2*a* (*k*_1*a*→2*a*_ = 1.7 × 10^−3^ μM^-1^ s^-1^), and therefore *1a* is never significantly populated, and is not detected once the equilibrium is reached (Figure **8**C). 2-tetrad conformers are also possible contributors to the isotopic population of 23TAG•2K^+^. However, the 1*b* → 0 → 1*a* → 2*a*/2*b* reaction pathway occurs at a much slower rate than the isotopic population interconversion due to the very slow rate-limiting step 0 → 1*a*. In addition, the conversion from 2-to 3-tetrad species requires visiting fully unfolded states. This is inconsistent with the detectable protection of the highly-exchanged isotopic population of 23TAG•2K^+^ that suggests only partial unfolding. We also verified the absence of significant shift in the equilibrium across the HDX/MS data points, which could have contributed (by e.g., concentration increase of the 2-tetrad G4 binding a second potassium cation non-specifically (Figure S**8**B). Two-tetrad G4s and unfolded strands are therefore unlikely to significantly contribute to the exchange of 23TAG•2K^+^.

A molecular mechanism compatible with the observed EX1/EX2 exchange kinetics is therefore the unfolding of 23TAG•2K^+^ into a partially unfolded ensemble,^70^ which may resemble the folding intermediate *1a*, followed by cooperative exchange of many sites before refolding can occur, as often observed for proteins (Figure **8**D).^23^ Although this ensemble is invisible at equilibrium, it is here revealed by H/D exchange. The exact nature of these unfolded states is a matter of debate.^16,72,74^ Our results suggest they are likely to bind specifically one potassium cation, and to have H-bonds in a few base pairs, triplets, or tetrads that provide some, but not much, protection against exchange. Conformers such as G-hairpins,^67,69,75–78^ G-triplexes,^48,67–69,76,77,79–82^ or strand-slipped conformers^77,81,83^ that all can bind K^+^ and contain fewer, less stable and more accessible H-bonds between nucleobases are likely candidates. Simultaneously, 23TAG•2K^+^ can exchange *via* EX2 by visiting microstates corresponding to local rearrangements such as breathing motions.

##### 3.2.3.2. Other instances of EX1 exchange

Other oligonucleotides exchange partially by EX1: both the 2- and 3-tetrad species formed by 22AG, 26TTA, and 222T as well as the 2-tetrad TBA and 22CTA G4s (Figures S**39**—S**49** and Tables S**7** and S**8**). Overall, these G4s exchange faster than the stable, pure EX2 species, have lower apparent *T*_*m*_ (below 42°C), and some are not fully folded at 22°C (e.g., 26TTA, TBA, 22CTA). However, there are very large differences in the extent of EX1 exchange among these species that cannot be explained by the *T*_*m*_ alone. The 2-tetrad G4s rapidly have mostly highly exchanged populations, and more than 3-tetrad G4s (e.g., compare 222T•2K^+^ and 222T•1K^+^).

The unfolding leading to an EX1-type exchange is always extensive, as few sites remain deuterated at the earliest measured times. In some cases (e.g., 26TTA·2K^+^, TBA), the low-mass population is even completely exchanged during the dead time. Thus, unfolding is complete enough to make all sites available for exchange and/or the few potentially protected sites exchange too fast to be observed, which is consistent with their low stability. After deconvolution, the “pure” EX2 contributions to these species are more in-line with the *NUS* vs. *T*_*m*_ trend of the more stable G4s, although still faster (Figure S**50**). This further shows that EX1 kinetics is a major contributor to the overall exchange of low-*T*_*m*_ G4s.

In section 3.1.3, we showed that a decrease in stability for a given G4 is accompanied by an increase in EX2 exchange rates. A sufficiently large decrease in stability (*ΔG*°_*unfolding*_ < 20 *kJ*/*mol*) also promotes EX1. This can be seen for T95-2T (Figure **5**), 222T•2K^+^ (Figure S**45**) and T30177TT (Figure S**51**) in 0.1 mM KCl solutions. The EX2 exchange is slightly faster, but the EX1 contribution accounts for most of the apparent increase in the global exchange rate. This suggests a greater propensity of these G4s to unfold cooperatively (decrease of *ΔΔG*° between the G4 and its unfolded conformers) rather than only an increase in their local dynamics. Both T95-2T and T30177TT appear fully folded under these conditions, further showing that HDX/MS reveals unfolded states that are not significantly populated and therefore invisible to most other techniques.

## 4. Conclusions

HDX/native MS was employed to study G4-forming oligonucleotides with varying topologies and stabilities. The salient points of this work are:

i. A time range spanning several orders of magnitudes (seconds to hours) is necessary to observe the exchange of all structured sites in G4 species. In particular, some structural features alter protection in specific time windows. The recommended time range is thus similar for nucleic acids as for protein HDX/MS.^62^
ii. By coupling HDX to native MS, one can obtain HDX data for conformers that bind different numbers of cations, providing access to exchange kinetics of several ensembles in a single experiment.
iii. The number of protected sites depends on the oligonucleotides’ secondary structure, not the primary structure. This is essential for structural applications. The number of protected sites indicates the number of tetrads, based on our reference dataset.^60^
iv. The exchange rates decrease with increasing G4 stability 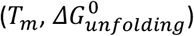 but are also sensitive to conformational dynamics and specific structural motifs. For stable conformers, the exchange proceeds through local breathing rather than a macroscopic 2-state unfolding, and thus HDX should not be used to predict stability, but to contextualize the results. For example, outlying results may reveal a special folding. Compared to bulk *T*_*m*_ and *ΔG*^0^ values derived from melting experiments, HDX/MS provides an assessment of G4 stability that is isothermal, avoiding the formation of alternative conformers at higher temperatures.^84^
v. The exchange scenario depends on the stability, number of tetrads, and topology of the G4. Two limiting cases are identified:
  i. Stable G4s exchange rarely and one site at a time according to local fluctuations (EX2 exchange; microstates in **8**D). Molecular dynamics may be used to characterize the breathing modes that favor this scenario.
  ii. Less stable G4s, especially those with two tetrads, can furthermore exchange several sites simultaneously via partial or complete unfolding (EX1 exchange; Figure **8**D). The topology of G4s likely influences the nature of the less-folded states they visit. These intermediate states can thus be detected by HDX, acting as an isotopic stamp when they are visited, even if they are overall sparsely populated. Exploiting the extent and rate of EX1 kinetics thanks to the deconvolution of isotope distributions thus opens the door to further exploring the folding landscapes of G4s.

The first three points demonstrate that native HDX/MS can help assign topologies to polymorphic G4s. For example, native HDX/MS clearly shows that human telomeric sequences such as 22AG, 23TAG, and 26TTA can coexist in equilibrium as 2- and 3-tetrad G4s. However, the current method is limited to the separation of conformers of different masses. Characterization of conformers of the same mass (e.g., the hybrid-1 and hybrid-2 topologies of 23TAG•2K^+^) requires an additional separation method, such as ion mobility spectrometry, which we will describe in due course. Direct infusion of exchanging samples also limits the scope of analysis to room temperature and the use of MS-compatible buffers. To perform HD exchange in a wider range of experimental conditions, bottom-up and/or on-line buffer exchange approaches need to be developed.

## 5. Data availability

The processed HDX/MS data of the oligonucleotide panel is deposited on Zenodo.^60^

## 6. Supporting Information

Methods and results of spectroscopic methods, MS tune softness, HDX/native MS raw data examples, processing principle and processed data, HDX/NMR methods and results, complete HDX/MS panel results including non-linear fitting results and diagnostics, isotope distribution deconvolution principle and results, and exchange scenario summary.

## 7. Acknowledgments

This work was supported by the Agence Nationale de la Recherche [ANR-21-CE29-0004 “DNA-HDXMS” to E.L.]. We thank Frédéric Rosu for invaluable support, and the structural biophysico-chemistry platform of the Institut Européen de Chimie et Biologie for the access to mass spectrometers.

