## Supporting Information for "A General Framework to Interpret Hydrogen-Deuterium Exchange Native Mass Spectrometry of G-Quadruplex DNA"

#### G4 structures

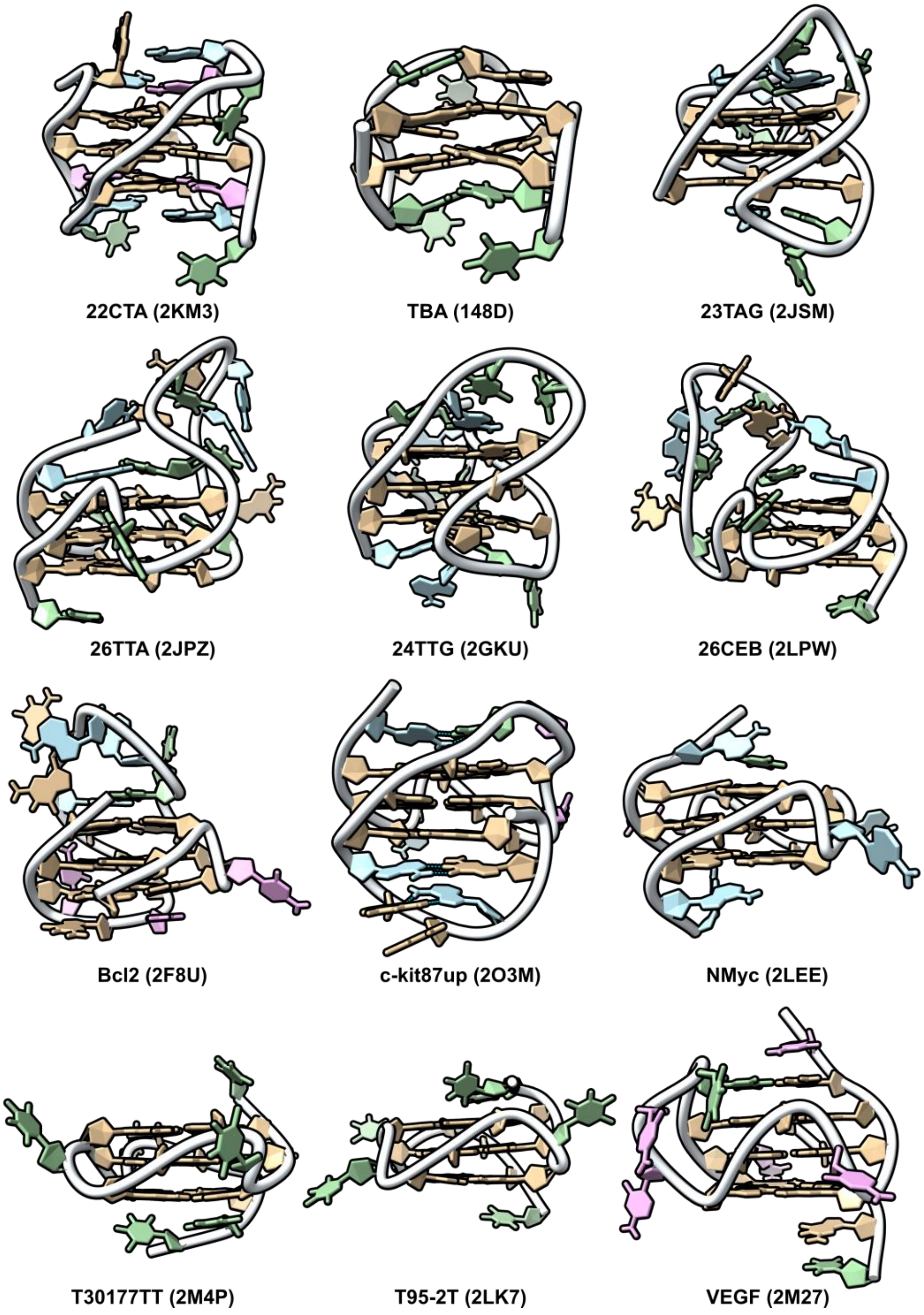


***Figure S1:*** *PDB-deposited NMR structures of selected G4s*

### UV-melting

UV/vis spectra were acquired from 220 to 350 nm with an interval of 1 nm, a bandwidth of 2 nm, and an averaging time of 0.1 s, then blank and baseline corrected. The molar concentrations in DNA strands were determined from the UV absorbance at 260 nm and the molar extinction coefficients calculated using the nearest-neighbor model, as described in reference[1].

UV-melting experiments were performed on a UVmc2 double-beam spectrophotometer (SAFAS, Monte Carlo, Monaco) equipped with a high-performance Peltier temperature controller and a thermostatable 10-cell holder, with 400-µl, 1-cm pathlength quartz cuvettes (115B-QS, Hellma GmbH & Co. KG, Müllheim, Germany). The samples were prepared by diluting the oligonucleotides to 10 µM in 100 mM TMAA, 1 mM KCl buffers (unless otherwise mentioned). The absorbance was monitored at 260, 295 and 335 nm on a cycle composed of a cooling ramp from 90°C to 4°C at 0.2°C min$^{-1}$, followed by a plateau of 15 minutes and a heating ramp back to 90°C (0.2°C min$^{-1}$). Data points were acquired every 60 s with an averaging time of 0.5 s.

UV-melting data was processed using the in-house *MeltR* software as described previously[1], by non-linear fitting of the raw data with Equation (12), where $T$ is the temperature (in K), $T_m$ is the melting temperature (in K), $a$ and $b$ are the slopes and intercepts, respectively, of the folded ($F$) and unfolded ($U$) baselines, and $R$ is the gas constant (in J K$^{-1}$ mol$^{-1}$). The wavelength of interest was 295 nm for G4s and 260 nm for ds-DNA, and 335 nm was used as a blank. The data from a blank sample containing only the buffer was subtracted as well. The folded fraction $\theta_T$ at a given temperature was derived from the optimized baseline parameters, as described by Mergny and Lacroix[2].

$$A_T = (a^F \times T + b^F) \times \frac{1}{1+exp\left(-\frac{\Delta H^\circ\left(1-\frac{T}{T_m}\right)}{RT}\right)} + \left(a^U \times T + b^U \times \frac{exp\left(-\frac{\Delta H^\circ\left(1-\frac{T}{T_m}\right)}{RT}\right)}{1+exp\left(-\frac{\Delta H^\circ\left(1-\frac{T}{T_m}\right)}{RT}\right)}\right) \quad (12)$$

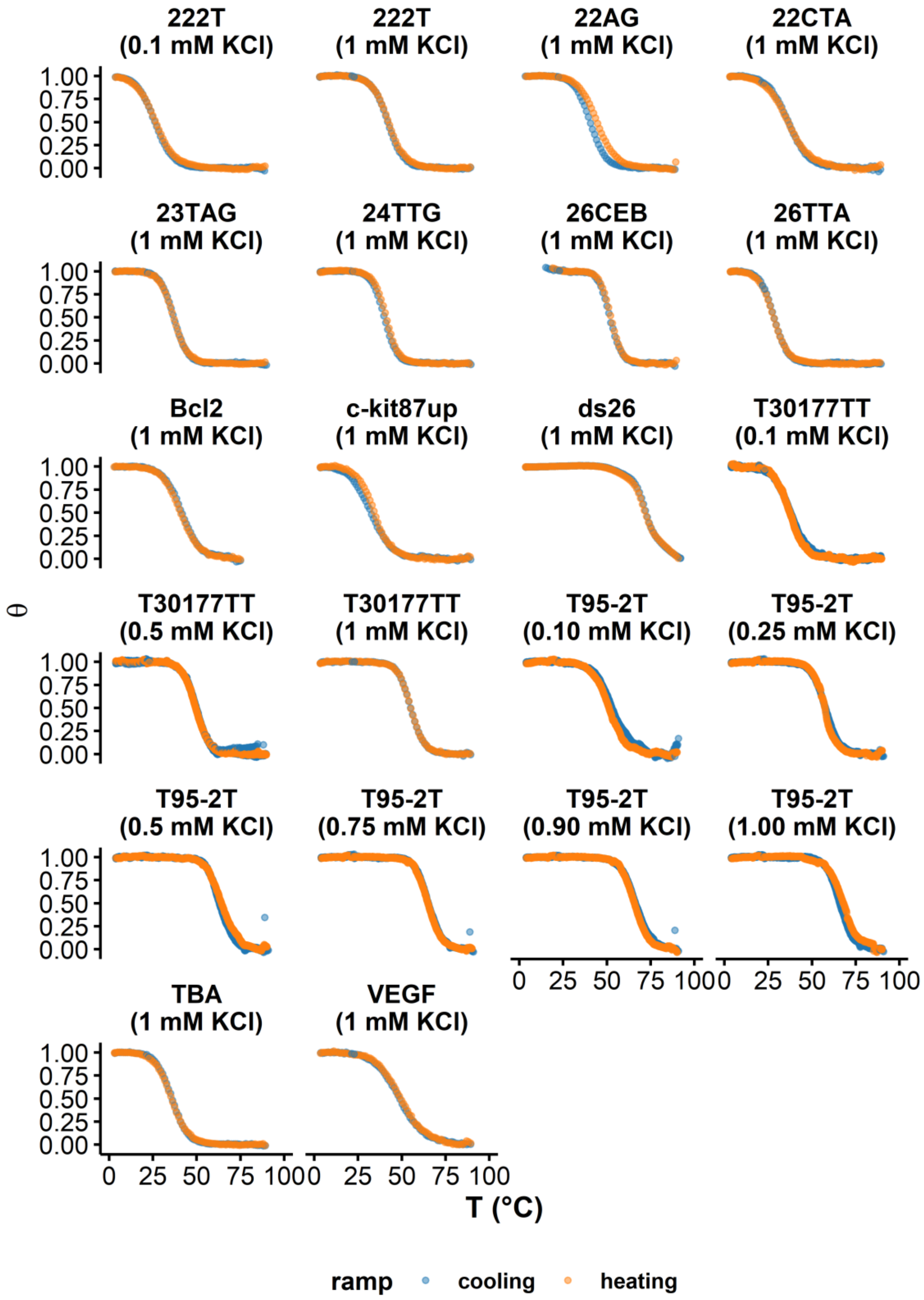


***Figure S2:*** *Folded fraction as function of the temperature, determined from UV-melting experiments at 295 nm (except ds26: 260 nm)*

#### Circular dichroism

Circular dichroism (CD) experiments were performed on a Jasco J-815 spectrophotometer equipped with a Jasco CDF 426S Peltier temperature controller using quartz cuvettes (10 mm path length) at 22°C. The samples were prepared by diluting the oligonucleotides to 10 µM in 100 mM TMAA, 1 mM KCl buffers (unless otherwise mentioned). The scanning range was 220—350 nm with a 1 nm data interval, 2 nm bandwidth, and 1 s response. Three accumulations were acquired with a scan speed of 50 nm/min. Raw data was blank and zero corrected (using the baseline between 325 and 350 nm), then converted to molar ellipticities $\Delta\varepsilon$ using equation (13), where $\theta$ is the ellipticity in mdeg and $C$ the molar concentration in strand, which was obtained as described above.

$$\Delta\varepsilon = \frac{\theta}{32980 \times C} \quad (13)$$

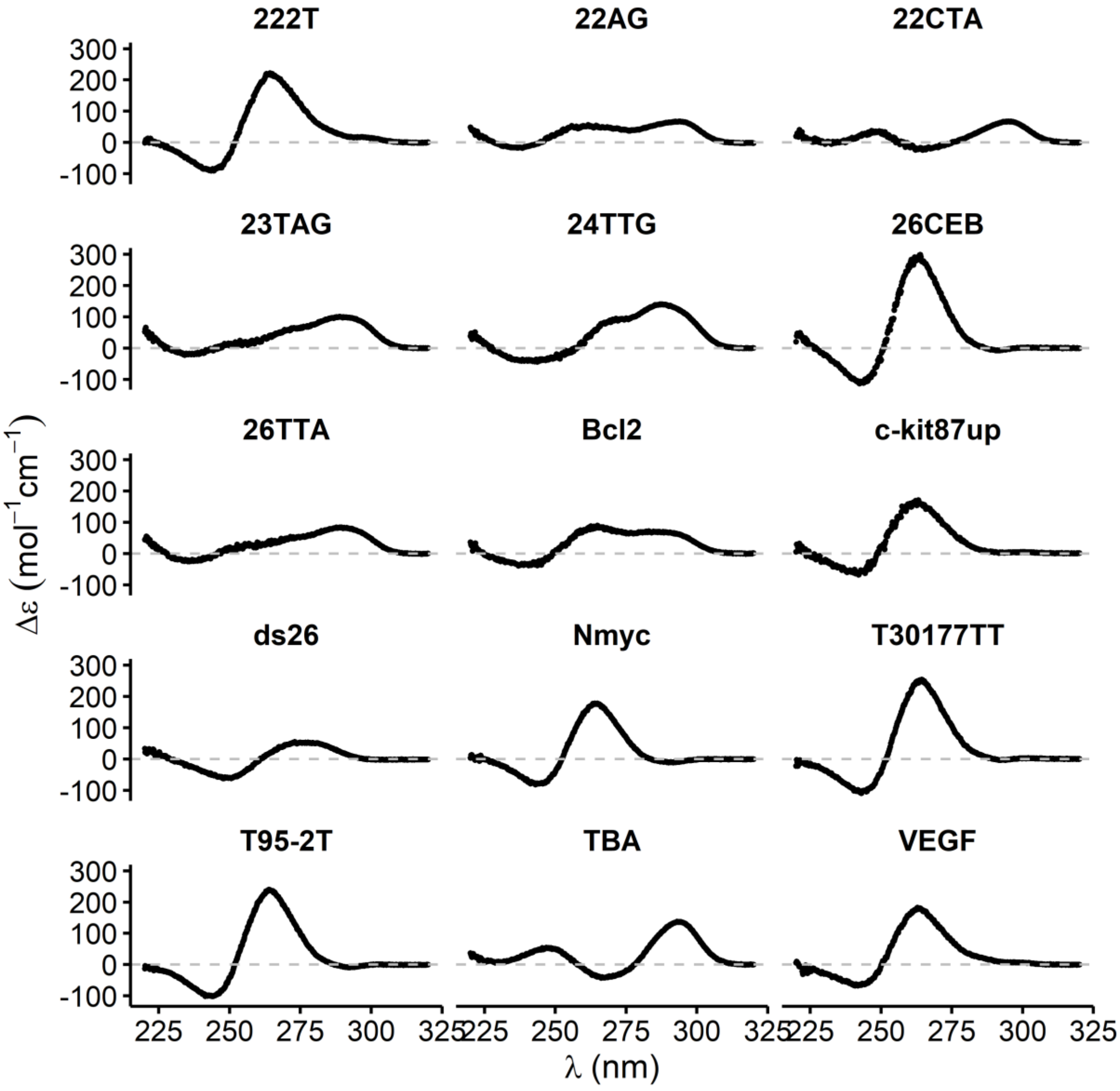


*__Figure S3:__ CD spectra acquired in 100 mM TMAA + 1 mM KCl.*

### HDX/NMR experiment

#### Sample preparation

The 500 µL exchange sample contained 150 µM T30177TT, 100 mM TMAA, 1mM KCl in 9% $D_2O$.

#### NMR parameters

All 1H NMR spectra were acquired on a 700 MHz NMR Bruker Biospin instrument with a TXI probe at 25°C, in 5 mm diameter NMR tubes (Wilmad from CortecNet, France), with a proton relaxation time of 6 s and 32k points/spectrum. The Watergate W5 pulse sequence combined to the "jump and returns" water suppression technique was used[3]. The number of scans was set using to 8 (acquisition time = 58 s) for time points until 28 min of exchange, then 48 (acquisition time = 5.8 min) afterwards, with no dummy scans. The *multizg* function was used to chain these acquisition steps. TopSpin 4.1 (Bruker) was used for data acquisition and processing. The lock and shim steps were achieved on a 200 µM oligonucleotide sample.

#### Reference

The non-exchanged reference (t = 0) spectrum of T30177TT in the conditions described above was acquired one day before the kinetics experiment (both with 8 and 48 scans). The 500 µL sample was frozen in liquid nitrogen then lyophilized overnight in a BenchTop Pro freeze-dryer at 180 µbar and −84°C.

#### Exchange kinetics

The dried sample (9% D2O) was dissolved in 500 µL of $D_2O$. The sample was mixed for 142 s then introduced in the instrument. 61 spectra were recorded for a total of seventeen hours, with a total dead time of around 4 minutes (*i.e.*, manual mixing time + time of first spectrum acquisition). The exchange time was defined as the time at which the acquisition starts plus the mean of its duration and the manual mixing time.

#### Data processing

After manual phase-adjusting and baseline correction, the imino peaks maxima which had enough signal at 8 scans (i.e G5+9, G13, G17 and G10+6) were picked at every time point. In two cases (G5/G9, G10/G6), the NMR peak resolution was not sufficient to be processed separately. The raw intensities ($I$) for each peak were normalized by the number of scans ($ns$).

$$I' = \frac{I}{ns}$$

The fully-exchanged peak intensities at 90% $D_2O$ ($I_{90}'$) was defined as 10% of the corresponding peak intensities of the non-exchanged reference sample ($I_9'$). For a given imino site, the fraction of remaining protons $f_H$ at exchange time $t$ is the difference between the peak intensity $I'(t)$ and that of the fully-exchanged reference ($I_{90}'$), normalized by the difference between the 9% and 90% $D_2O$ peak intensities.

$$f_H(t) = \frac{I'(t) - I_{90}'}{I_9' - I_{90}'}$$

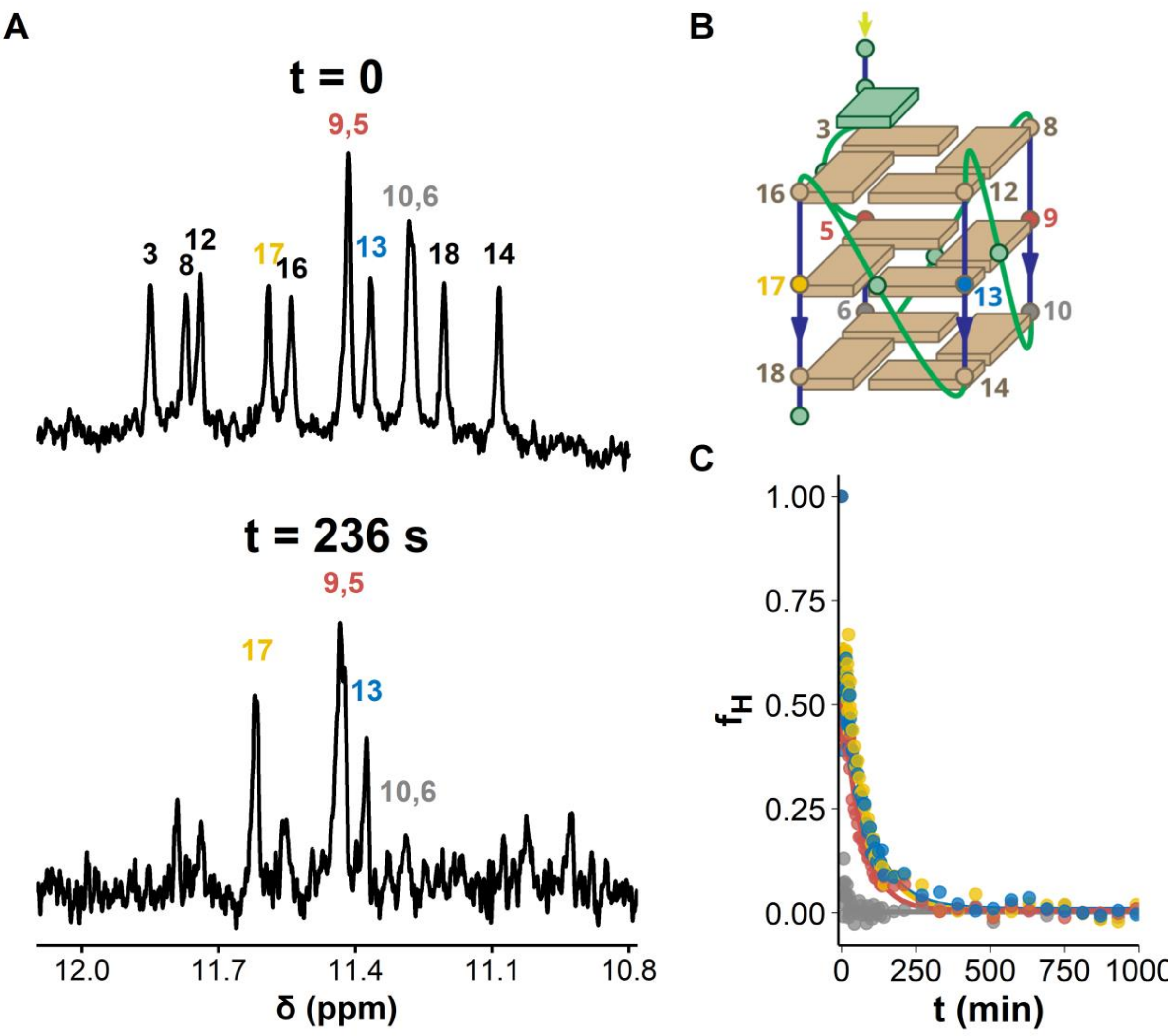


***Figure S4:*** *HDX/NMR kinetics of T30177TT. A. 1H-NMR spectra before exchange (t = 0) and after 236 seconds of exchange in D$_2$O. Peak are labeled with guanine numbers as described in panel B. C. fraction of non-exchanged imino protons against exchange time. External tetrads are mostly exchanged within the dead time (4 min). Internal tetrads exchange over the course of several hours with little differences between guanines. A slightly faster exchange is visible for dG5/dG9, which may be due to the presence of a bulging dT4.*

#### Tune softness

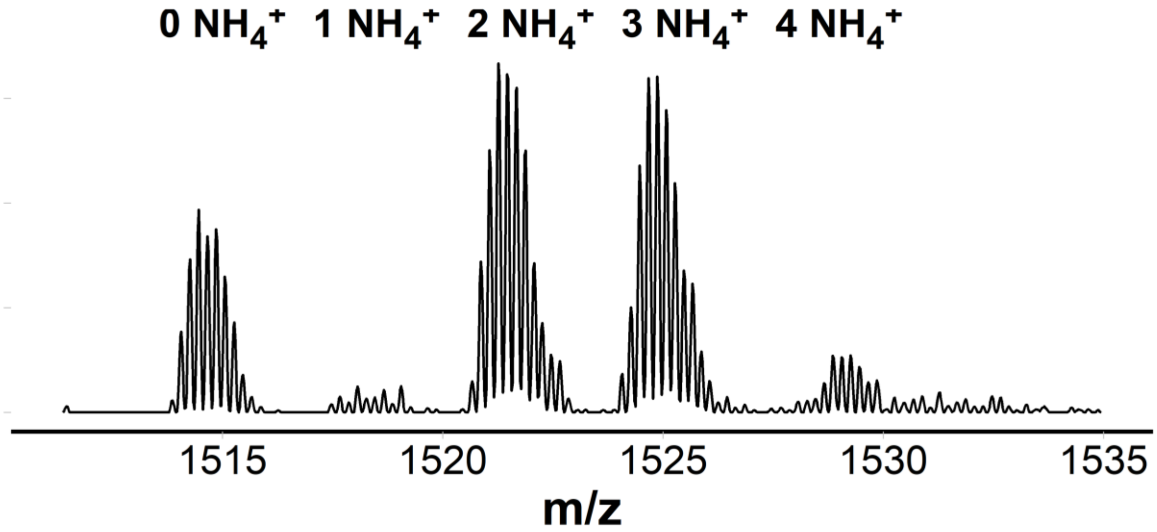


***Figure S5:*** *Mass spectrum of a 10-µM solution of [G4T4G4]2 G-quadruplex in 100 mM ammonium acetate, zoomed on the 5- charge state.*

#### Raw HDX data examples

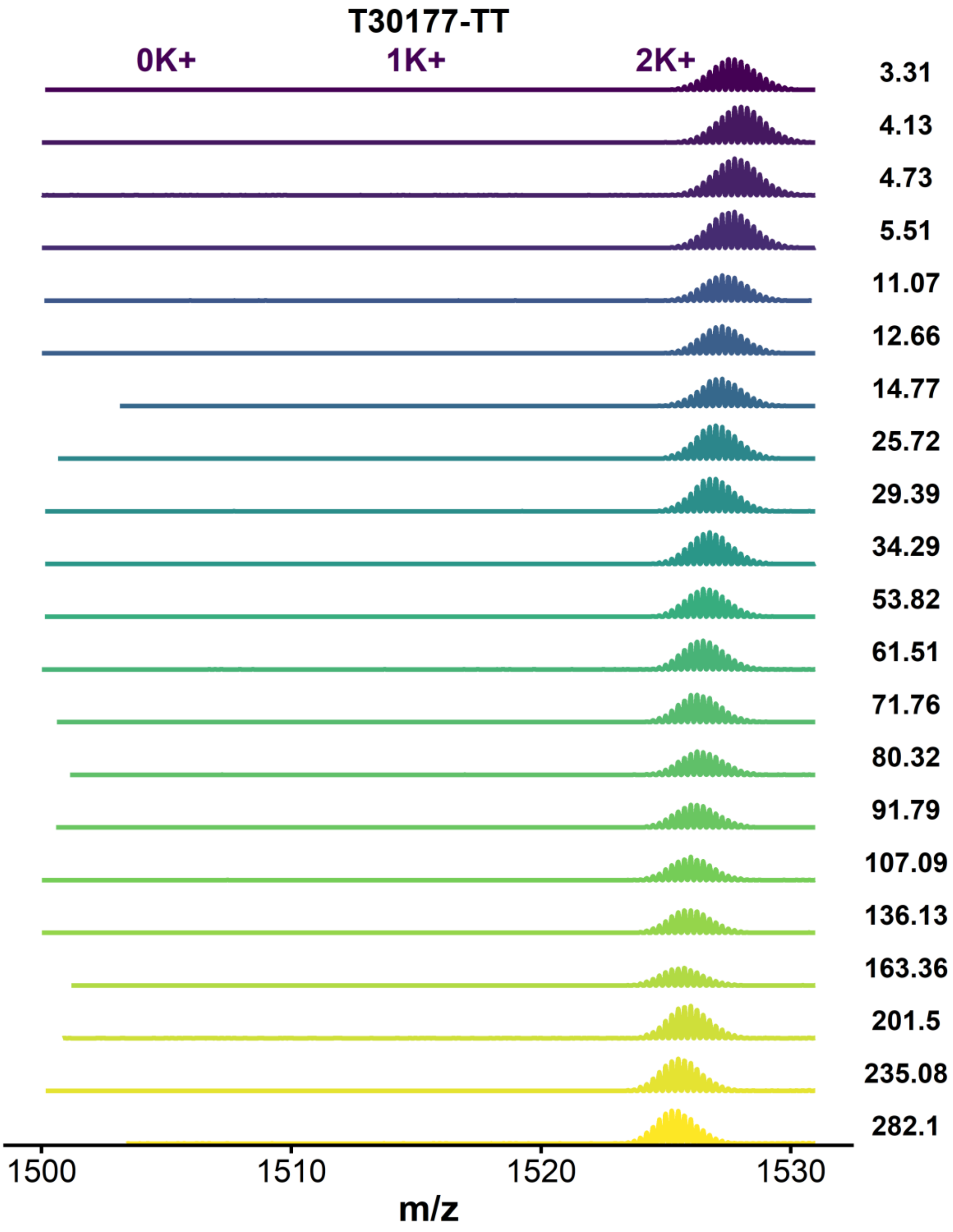


***Figure S6:*** *MS spectra of T30177-TT acquired by continuous-flow HDX experiments, focused on the 4- charge state species*

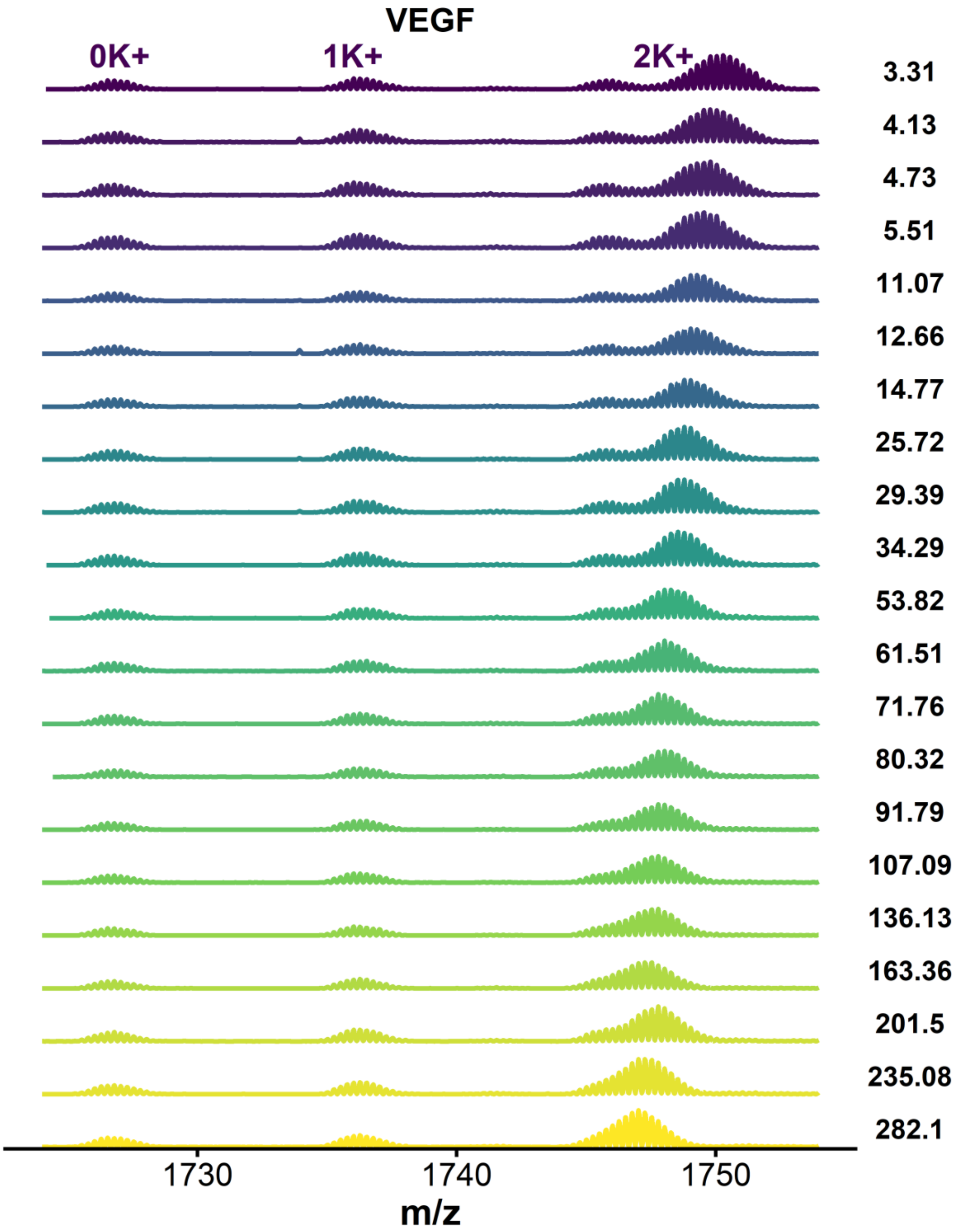


***Figure S7:*** *MS spectra of VEGF acquired by continuous-flow HDX experiments, focused on the 4- charge state species*

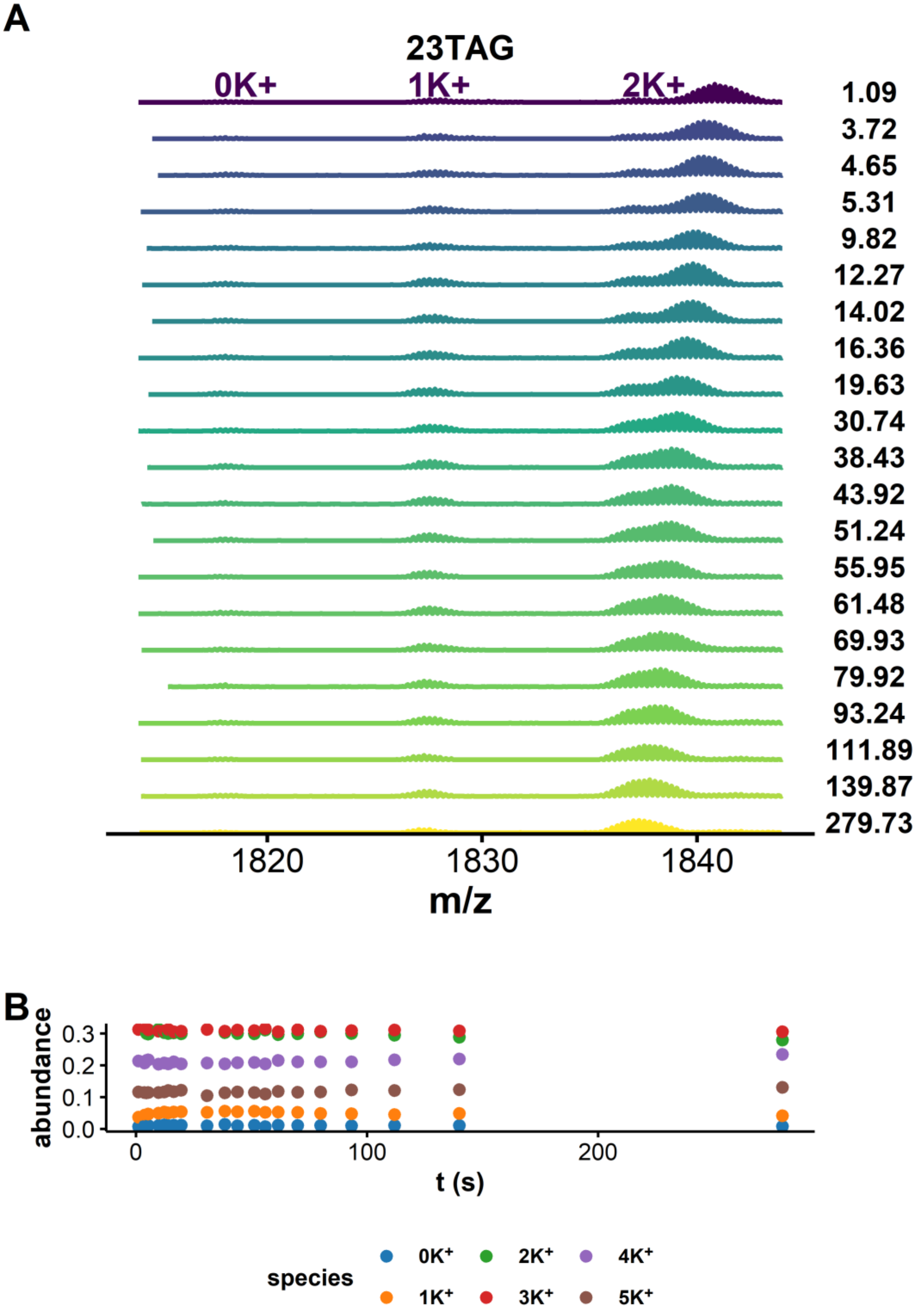


***Figure S8:*** *A. MS spectra of 23TAG acquired by continuous-flow HDX experiments, focused on the 4- charge state species. B. Relative abundance of the 4- species with potassium-binding stoichiometry from 0 to 5.*

#### Data processing

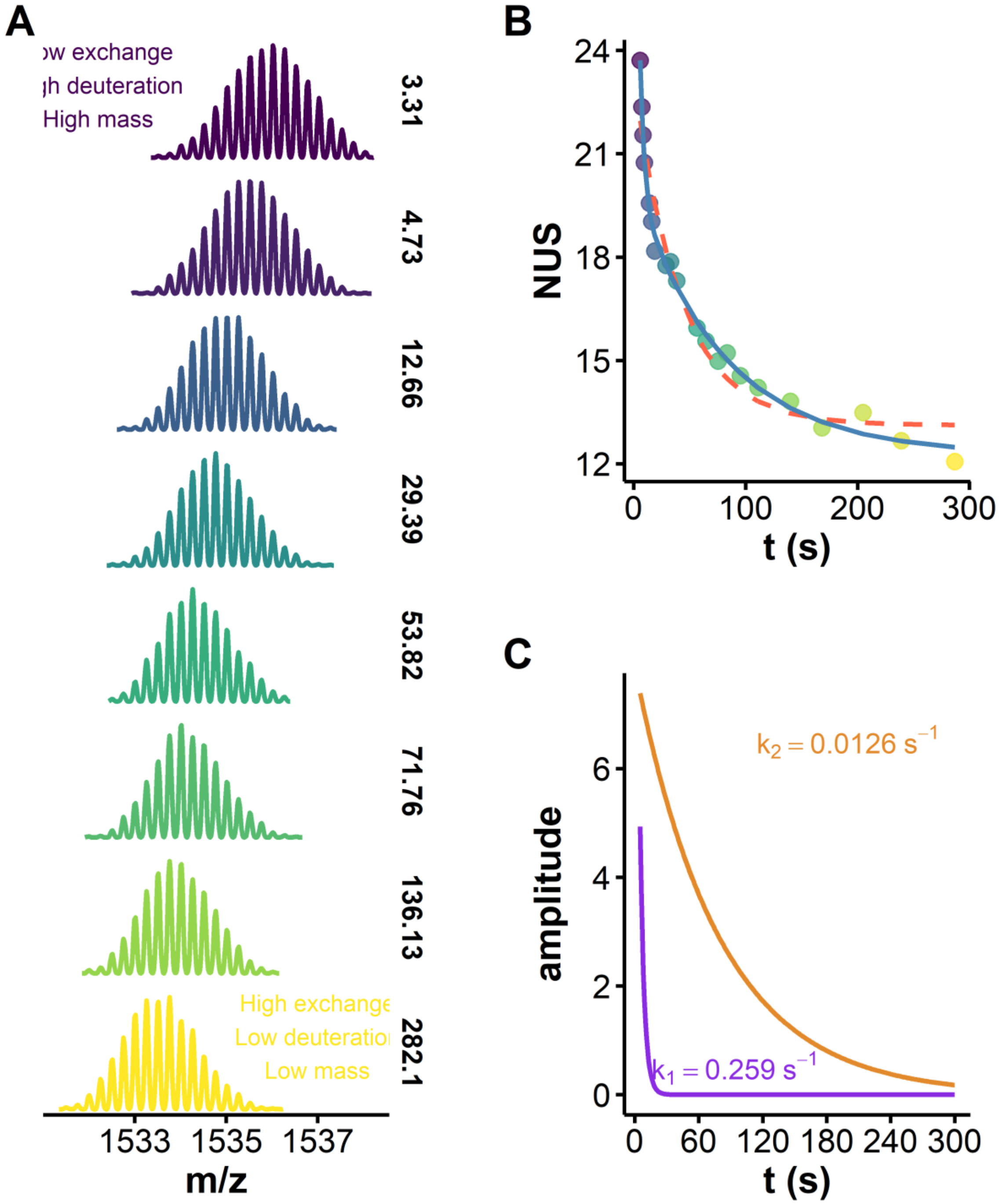


***Figure S9:*** *A. Isotopic distribution of the 4- charge state of NMyc as a function of the exchange time (D-to-H direction from top to bottom; time given in seconds, not all time points shown). B. Corresponding exchange kinetics plot, wherein centroids have been converted to NUS, and non-linear fitting was performed using equation (4) (blue: j = 2, dashed orange: j = 1). C. Decomposition of the two exponential regimes (faster exchange: purple, slower exchange: orange)*

#### HDX/MS exchange plots for the panel oligonucleotides

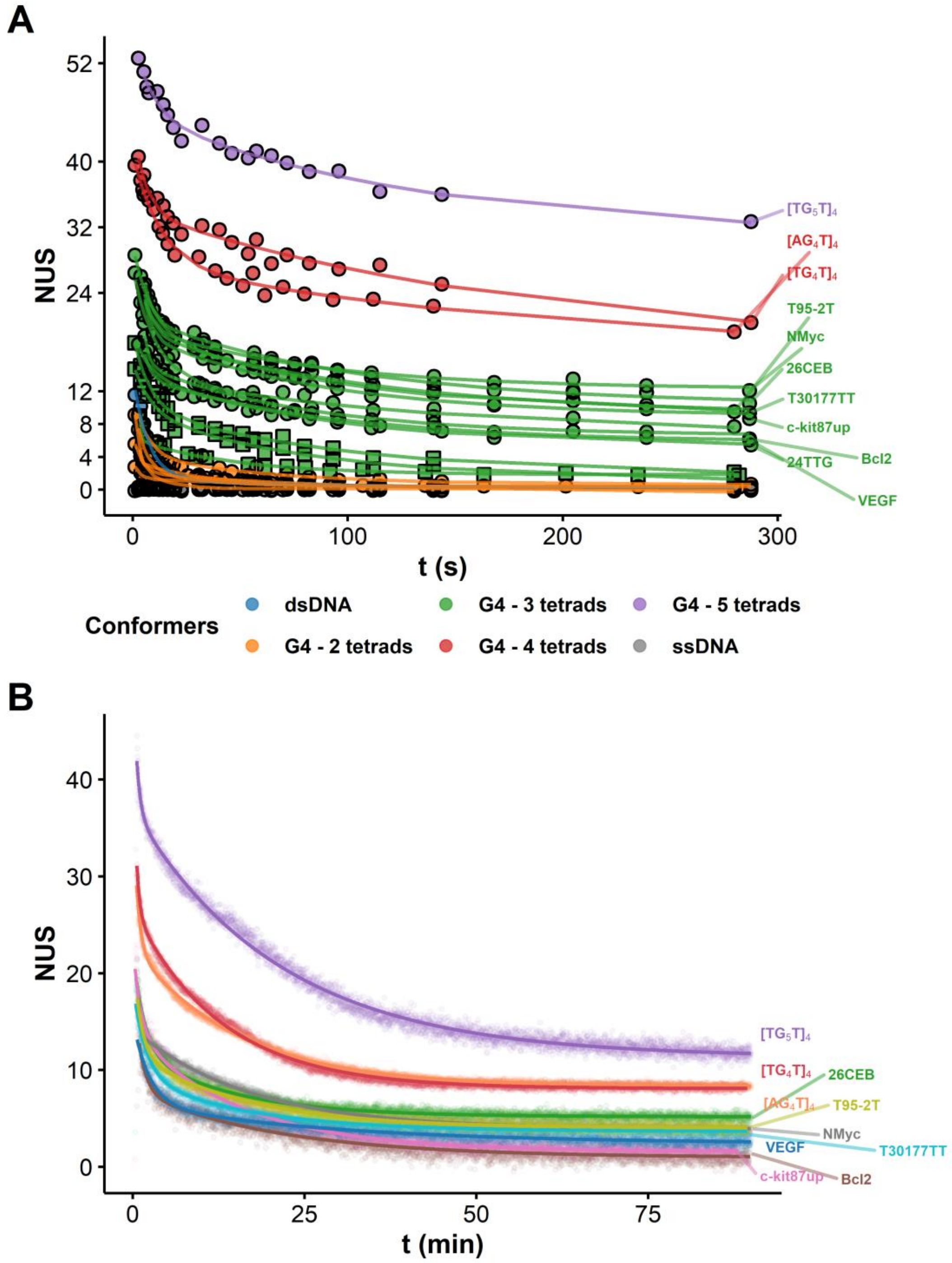


*Figure S10:* *Apparent HDX/MS kinetics of a panel of oligonucleotides. A: continuous-flow measurements, colored by number of tetrads, B: real-time measurements, colored by oligonucleotides. The lines are the results of non-linear fitting with equation (4), with j = 2. Some species are not labelled for the sake of clarity.*

#### Deconvolution of multimodal isotopic distributions

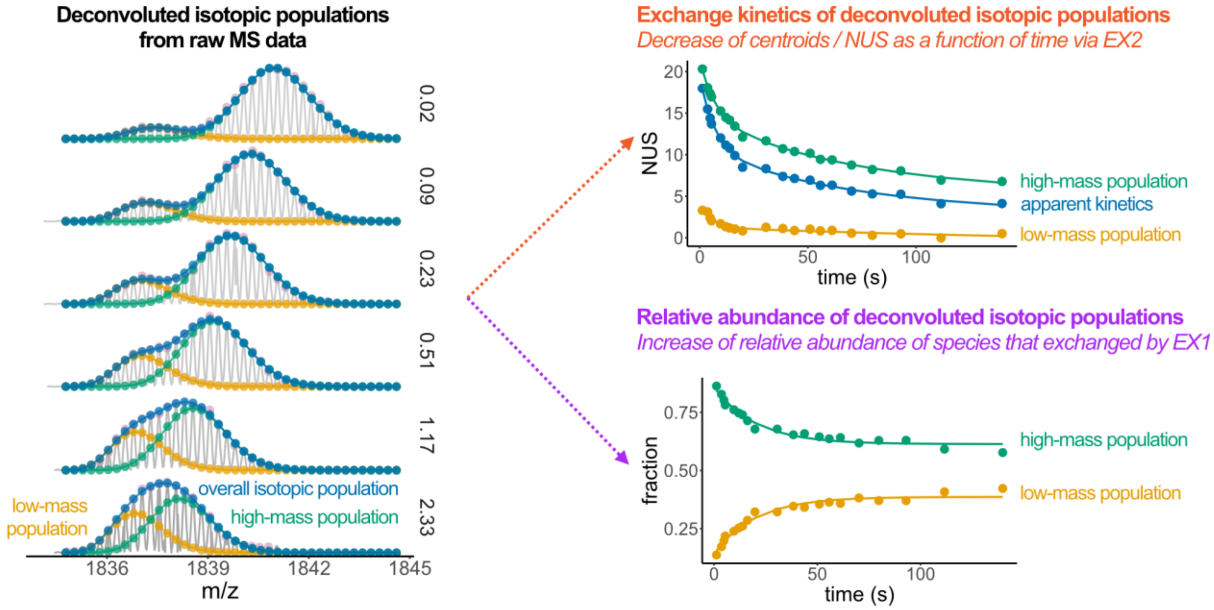


***Figure S11:*** *Principle of EX1/EX2 kinetics extraction from bimodal isotopic distributions.*[4]

### Non-linear fitting of the HDX/MS exchange data (continuous-flow)

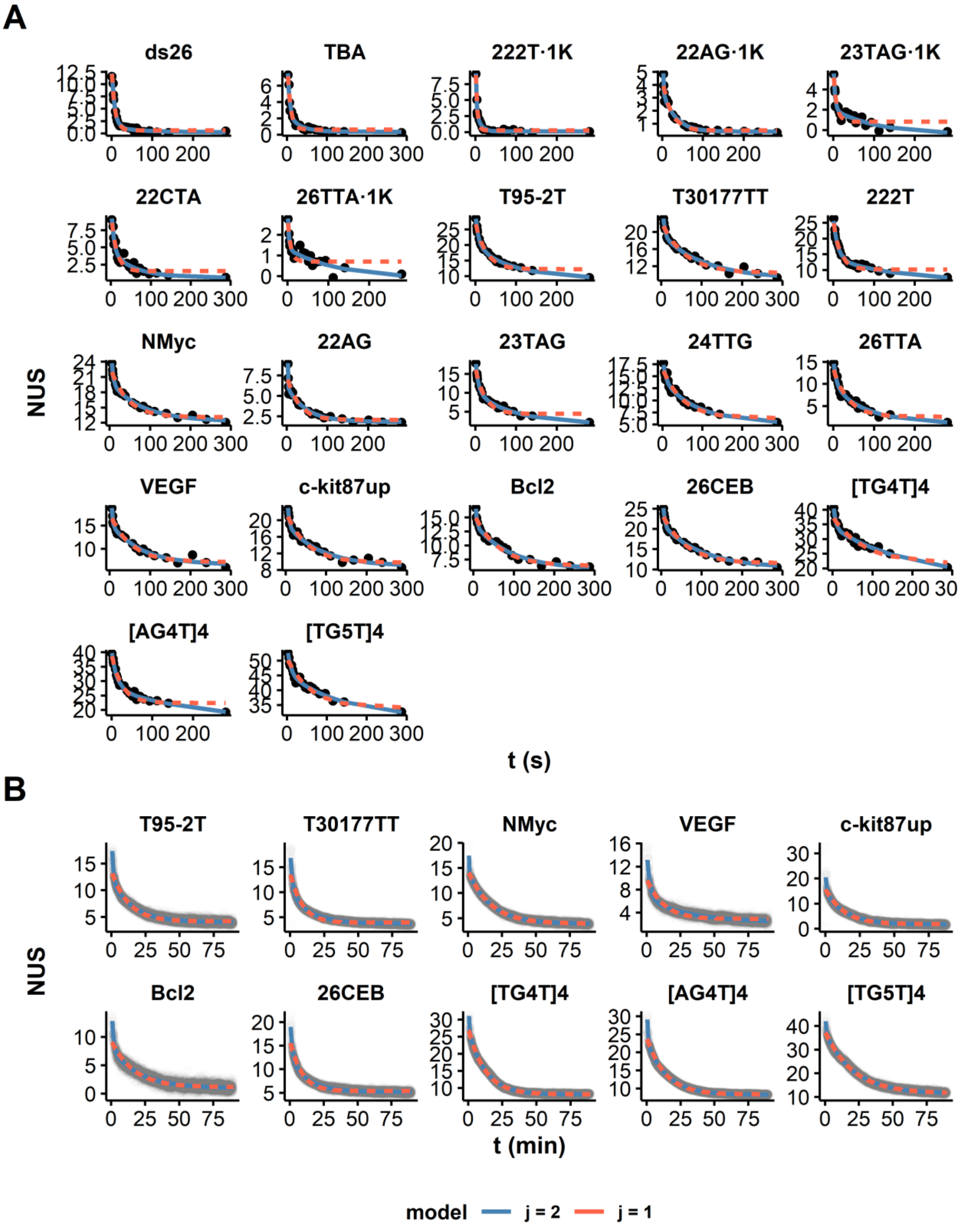


***Figure S12:*** *Comparison of the non-linear fitting the exchange data, using equation (4), with j = 1 (blue) or 2 (orange). A: Continuous-flow MS, B: Real-time MS*

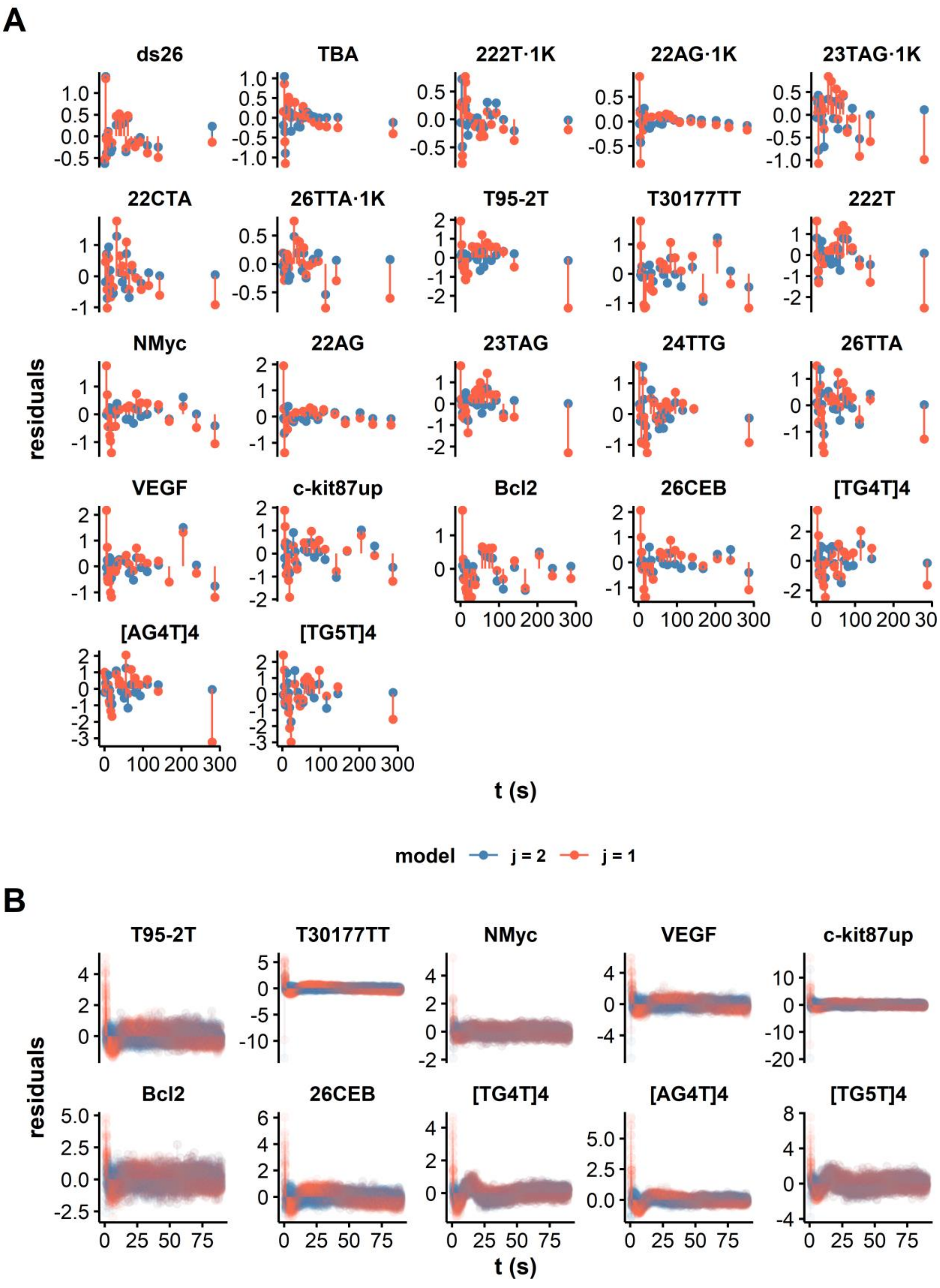


***Figure S13:*** *Comparison of non-linear fitting residuals of the exchange data, using equation (4), with i = 1 (blue) or 2 (orange). A: Continuous-flow MS, B: Real-time MS*

***Table S1:*** *Double exponential fitting parameters (continuous-flow MS)*

| **Analyte** | | **Estimates** | | | | | **Standard Errors** | | | | | **Significance (Pr(>\|t\|))** | | | | |
|---|---|---|---|---|---|---|---|---|---|---|---|---|---|---|---|---|
| **Oligonucleotide** | **Tetrads** | $N_1$ | $k_1$ ($s^{-1}$) | $N_2$ | $k_2$ ($s^{-1}$) | $NUS_\infty$ | $N_1$ | $k_1$ ($s^{-1}$) | $N_2$ | $k_2$ ($s^{-1}$) | $NUS_\infty$ | $N_1$ | $k_1$ ($s^{-1}$) | $N_2$ | $k_2$ ($s^{-1}$) | $NUS_\infty$ |
| $[TG_5T]_4$ | 5 | 9.5 | 0.119 | 15.2 | 0.007 | 30.8 | 1.6 | 0.044 | 1.5 | 0.003 | 2.1 | *** | * | *** | * | *** |
| $[AG_4T]_4$ | 4 | 13.4 | 0.077 | 11.4 | 0.005 | 16.1 | 1.4 | 0.014 | 3.7 | 0.004 | 4.9 | *** | *** | ** | | ** |
| $[TG_4T]_4$ | 4 | 10.5 | 0.167 | 17.5 | 0.005 | 16.4 | 2.0 | 0.058 | 3.2 | 0.002 | 3.8 | *** | * | *** | * | *** |
| 222T | 3 | 13.2 | 0.095 | 7.4 | 0.007 | 6.4 | 1.2 | 0.016 | 1.4 | 0.005 | 2.1 | *** | *** | *** | | ** |
| 22AG | 3 | 298.5 | 1.380 | 4.4 | 0.022 | 1.8 | 321.3 | 0.331 | 0.2 | 0.003 | 0.1 | | *** | *** | *** | *** |
| 23TAG | 3 | 8.9 | 0.133 | 8.5 | 0.010 | 1.6 | 0.7 | 0.021 | 0.6 | 0.002 | 0.6 | *** | *** | *** | *** | * |
| 24TTG | 3 | 4.9 | 0.147 | 9.1 | 0.012 | 5.2 | 1.3 | 0.085 | 1.0 | 0.003 | 0.8 | ** | | *** | ** | *** |
| 26CEB | 3 | 19.1 | 0.255 | 10.9 | 0.010 | 10.3 | 5.3 | 0.049 | 0.3 | 0.001 | 0.4 | ** | *** | *** | *** | *** |
| 26TTA | 3 | 6.0 | 0.123 | 8.5 | 0.012 | 1.0 | 1.3 | 0.052 | 1.2 | 0.004 | 0.9 | *** | * | *** | * | |
| Bcl2 | 3 | 20.4 | 0.336 | 8.3 | 0.011 | 5.8 | 13.1 | 0.112 | 0.4 | 0.002 | 0.4 | | ** | *** | *** | *** |
| NMyc | 3 | 18.0 | 0.259 | 7.9 | 0.013 | 12.3 | 5.2 | 0.052 | 0.3 | 0.001 | 0.3 | ** | *** | *** | *** | *** |
| T30177TT | 3 | 13.9 | 0.206 | 10.3 | 0.009 | 9.1 | 4.9 | 0.066 | 0.5 | 0.002 | 0.6 | * | ** | *** | *** | *** |
| T95-2T | 3 | 10.2 | 0.121 | 10.6 | 0.010 | 9.1 | 0.6 | 0.015 | 0.6 | 0.002 | 0.5 | *** | *** | *** | *** | *** |
| VEGF | 3 | 53.4 | 0.467 | 9.1 | 0.013 | 6.6 | 50.4 | 0.166 | 0.5 | 0.002 | 0.4 | | * | *** | *** | *** |
| c-kit87up | 3 | 20.6 | 0.274 | 10.3 | 0.012 | 8.9 | 12.1 | 0.105 | 0.6 | 0.002 | 0.6 | | * | *** | *** | *** |
| 222T·1K | 2 | 9.6 | 0.488 | 3.8 | 0.093 | 0.1 | 1.7 | 0.211 | 2.4 | 0.047 | 0.1 | *** | * | | - | |
| 22AG·1K | 2 | 115.2 | 1.289 | 3.2 | 0.032 | 0.4 | 151.8 | 0.411 | 0.2 | 0.004 | 0.1 | | ** | *** | *** | *** |
| 22CTA | 2 | 7.8 | 0.165 | 3.8 | 0.016 | 0.6 | 1.5 | 0.067 | 1.1 | 0.010 | 0.6 | *** | * | ** | | |
| 23TAG·1K | 2 | 4.7 | 0.259 | 2.5 | 0.009 | -0.5 | 0.7 | 0.073 | 0.6 | 0.006 | 0.7 | *** | ** | *** | | |
| 26TTA·1K | 2 | 2.2 | 0.354 | 1.5 | 0.008 | -0.1 | 0.5 | 0.137 | 0.4 | 0.005 | 0.5 | *** | * | ** | | |
| TBA | 2 | 9.2 | 0.221 | 2.2 | 0.033 | 0.4 | 1.4 | 0.068 | 1.3 | 0.024 | 0.2 | *** | ** | - | | |
| ds26 | 0 | 12.7 | 0.141 | 1.1 | 0.014 | 0.2 | 1.0 | 0.022 | 0.9 | 0.028 | 0.6 | *** | *** | | | |

***Table S2:*** *Single exponential fitting parameters (continuous-flow MS)*

| Analyte | | Estimates | | | Standard Errors | | | Significance (Pr(>\|t\|)) | | |
|---|---|---|---|---|---|---|---|---|---|---|
| **Oligonucleotide** | **Tetrads** | $N_1$ | $k_1$ ($s^{-1}$) | $NUS_\infty$ | $N_1$ | $k_1$ ($s^{-1}$) | $NUS_\infty$ | $N_1$ | $k_1$ ($s^{-1}$) | $NUS_\infty$ |
| $[TG_5T]_4$ | 5 | 16.8 | 0.017 | 34.1 | 1.1 | 0.003 | 1.2 | *** | *** | *** |
| $[AG_4T]_4$ | 4 | 16.8 | 0.039 | 22.5 | 0.8 | 0.005 | 0.6 | *** | *** | *** |
| $[TG_4T]_4$ | 4 | 16.1 | 0.013 | 21.6 | 1.5 | 0.003 | 1.6 | *** | *** | *** |
| 222T | 3 | 15.6 | 0.058 | 10.2 | 0.8 | 0.007 | 0.4 | *** | *** | *** |
| 22AG | 3 | 5.6 | 0.035 | 2.1 | 0.4 | 0.008 | 0.2 | *** | *** | *** |
| 23TAG | 3 | 12.2 | 0.042 | 4.4 | 0.7 | 0.007 | 0.5 | *** | *** | *** |
| 24TTG | 3 | 10.3 | 0.020 | 6.3 | 0.6 | 0.003 | 0.6 | *** | *** | *** |
| 26CEB | 3 | 12.6 | 0.017 | 11.5 | 0.6 | 0.002 | 0.5 | *** | *** | *** |
| 26TTA | 3 | 11.0 | 0.027 | 2.6 | 0.6 | 0.004 | 0.6 | *** | *** | *** |
| Bcl2 | 3 | 9.4 | 0.017 | 6.4 | 0.5 | 0.002 | 0.4 | *** | *** | *** |
| NMyc | 3 | 10.1 | 0.024 | 13.1 | 0.5 | 0.004 | 0.4 | *** | *** | *** |
| T30177TT | 3 | 12.0 | 0.017 | 10.4 | 0.6 | 0.003 | 0.5 | *** | *** | *** |
| T95-2T | 3 | 15.1 | 0.037 | 12.2 | 0.7 | 0.005 | 0.5 | *** | *** | *** |
| VEGF | 3 | 10.5 | 0.019 | 7.2 | 0.6 | 0.003 | 0.5 | *** | *** | *** |
| c-kit87up | 3 | 12.3 | 0.019 | 9.8 | 0.7 | 0.003 | 0.5 | *** | *** | *** |
| 222T·1K | 2 | 11.2 | 0.248 | 0.3 | 0.7 | 0.023 | 0.1 | *** | *** | * |
| 22AG·1K | 2 | 4.0 | 0.047 | 0.5 | 0.3 | 0.007 | 0.1 | *** | *** | *** |
| 22CTA | 2 | 8.5 | 0.073 | 1.6 | 0.8 | 0.013 | 0.3 | *** | *** | *** |
| 23TAG·1K | 2 | 5.0 | 0.124 | 0.8 | 0.6 | 0.028 | 0.2 | *** | *** | *** |
| 26TTA·1K | 2 | 2.2 | 0.136 | 0.7 | 0.4 | 0.045 | 0.1 | *** | ** | *** |
| TBA | 2 | 9.2 | 0.130 | 0.6 | 0.8 | 0.015 | 0.1 | *** | *** | *** |
| ds26 | 0 | 13.1 | 0.127 | 0.6 | 0.5 | 0.009 | 0.1 | *** | *** | *** |

***Table S3:*** *Model selection parameters (continuous-flow MS): lower Akaike information criterion (AIC) and Bayesian information criterion (BIC) indicate which model describe best the experimental data.*

| Data | | Double exponential (j = 2) | | | | | Single exponential (j = 1) | | | | |
|---|---|---|---|---|---|---|---|---|---|---|---|
| **Oligonucleotide** | **n** | **Degrees freedom** | **Sigma** | **Log-likelihood** | **AICc** | **BIC** | **Degrees freedom** | **Sigma** | **Log-likelihood** | **AICc** | **BIC** |
| $[TG_5T]_4$ | 21 | 16 | 0.85 | -23.54 | 65.07 | 65.34 | 18 | 1.34 | -34.36 | 79.22 | 80.90 |
| $[AG_4T]_4$ | 21 | 16 | 0.66 | -18.28 | 54.55 | 54.82 | 18 | 1.19 | -31.87 | 74.23 | 75.91 |
| $[TG_4T]_4$ | 21 | 16 | 0.88 | -24.34 | 66.69 | 66.95 | 18 | 1.48 | -36.38 | 83.25 | 84.93 |
| 222T | 21 | 16 | 0.62 | -16.80 | 51.60 | 51.87 | 18 | 1.02 | -28.58 | 67.67 | 69.34 |
| 22AG | 21 | 16 | 0.25 | 2.17 | 13.66 | 13.93 | 18 | 0.63 | -18.34 | 47.18 | 48.85 |
| 23TAG | 21 | 16 | 0.37 | -6.30 | 30.59 | 30.86 | 18 | 0.99 | -28.04 | 66.58 | 68.26 |
| 24TTG | 21 | 16 | 0.61 | -16.40 | 50.80 | 51.06 | 18 | 0.76 | -22.36 | 55.21 | 56.89 |
| 26CEB | 21 | 16 | 0.31 | -2.05 | 22.10 | 22.37 | 18 | 0.88 | -25.43 | 61.37 | 63.04 |
| 26TTA | 21 | 16 | 0.64 | -17.49 | 52.97 | 53.24 | 18 | 0.90 | -25.96 | 62.43 | 64.10 |
| Bcl2 | 21 | 16 | 0.37 | -6.29 | 30.57 | 30.84 | 18 | 0.67 | -19.80 | 50.11 | 51.79 |
| NMyc | 21 | 16 | 0.28 | -0.51 | 19.02 | 19.28 | 18 | 0.75 | -22.20 | 54.90 | 56.58 |
| T30177TT | 21 | 16 | 0.47 | -10.89 | 39.78 | 40.04 | 18 | 0.88 | -25.56 | 61.63 | 63.31 |
| T95-2T | 21 | 16 | 0.33 | -3.38 | 24.76 | 25.03 | 18 | 1.06 | -29.41 | 69.32 | 70.99 |
| VEGF | 21 | 16 | 0.52 | -13.26 | 44.53 | 44.79 | 18 | 0.86 | -24.90 | 60.30 | 61.98 |
| c-kit87up | 21 | 16 | 0.58 | -15.66 | 49.32 | 49.58 | 18 | 0.96 | -27.34 | 65.18 | 66.85 |
| 222T·1K | 21 | 16 | 0.31 | -2.41 | 22.81 | 23.08 | 18 | 0.43 | -10.57 | 31.64 | 33.32 |
| 22AG·1K | 21 | 16 | 0.16 | 11.15 | -4.30 | -4.04 | 18 | 0.32 | -4.43 | 19.37 | 21.05 |
| 22CTA | 21 | 16 | 0.61 | -16.47 | 50.94 | 51.21 | 18 | 0.74 | -21.81 | 54.11 | 55.79 |
| 23TAG·1K | 21 | 16 | 0.40 | -7.59 | 33.18 | 33.44 | 18 | 0.57 | -16.42 | 43.33 | 45.01 |
| 26TTA·1K | 21 | 16 | 0.24 | 2.67 | 12.67 | 12.94 | 18 | 0.36 | -6.50 | 23.50 | 25.18 |
| TBA | 21 | 16 | 0.40 | -7.61 | 33.23 | 33.49 | 18 | 0.44 | -10.99 | 32.48 | 34.16 |
| ds26 | 21 | 16 | 0.48 | -11.61 | 41.22 | 41.49 | 18 | 0.48 | -12.82 | 36.13 | 37.81 |

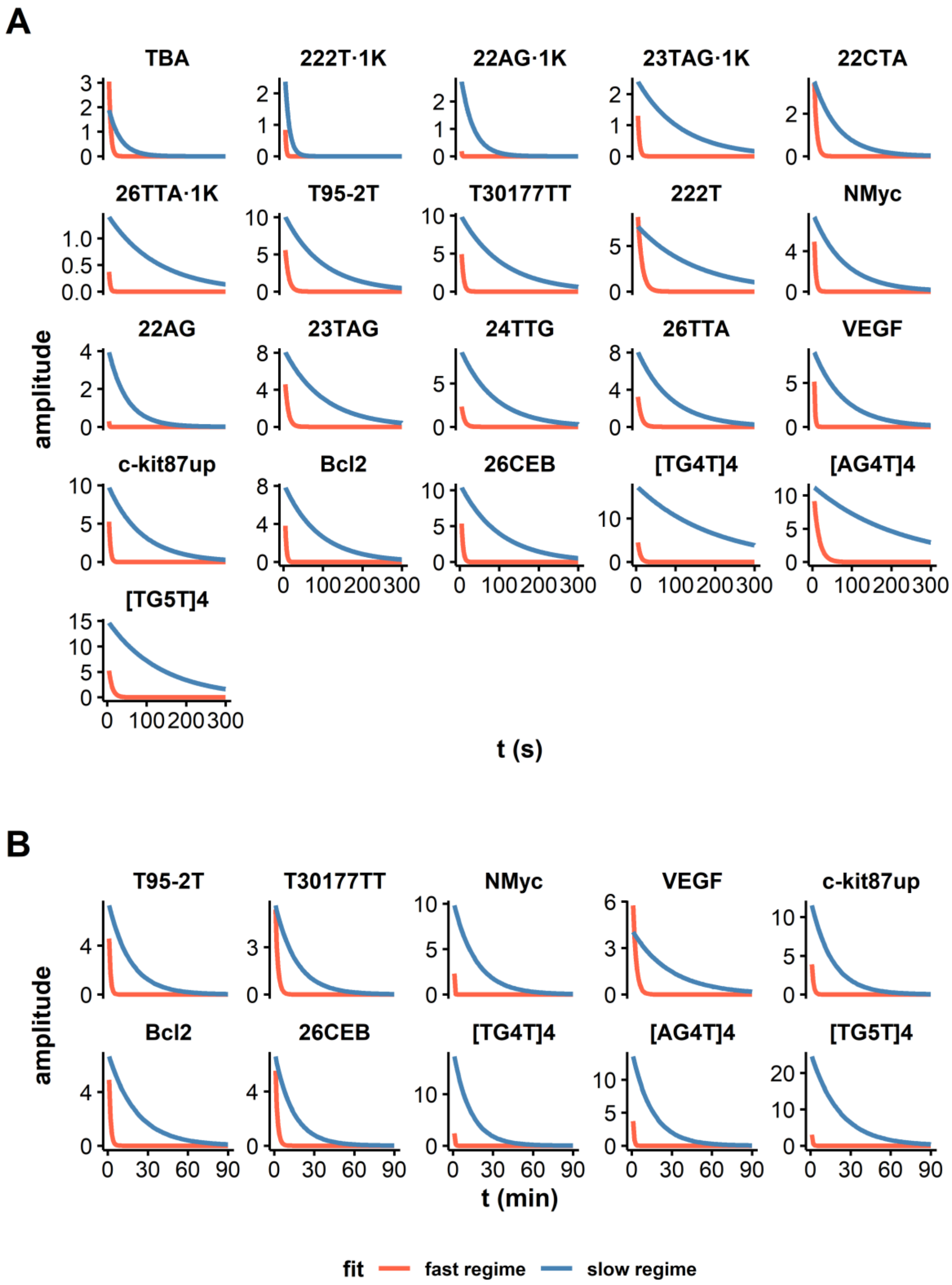


***Figure S14:*** *Decomposition of the bi-exponential fitting of 3-tetrad G4 exchange plots obtained by A. continuous-flow and B. real-time MS.*

#### NUS areas

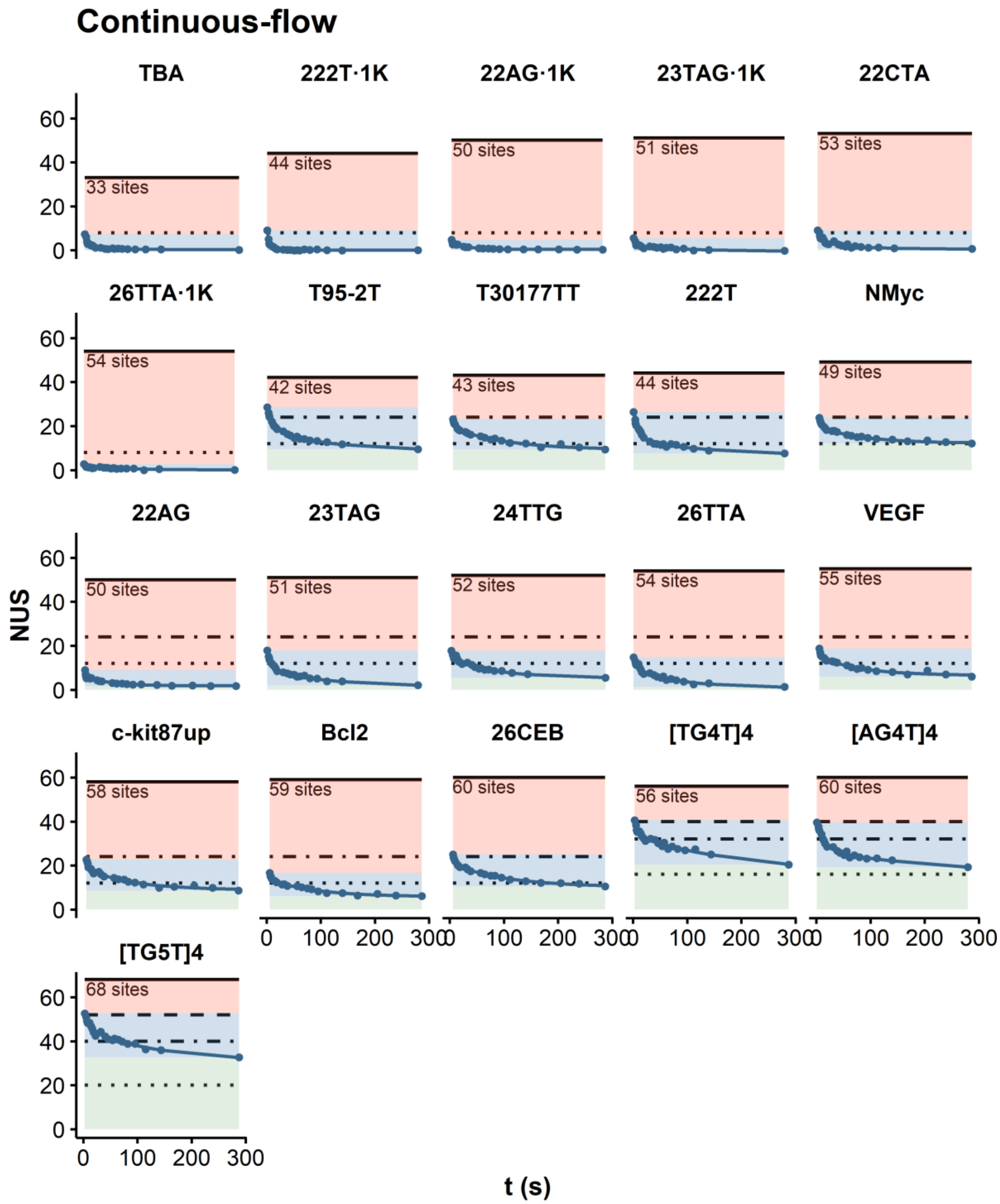


*Figure S15: Continuous-flow exchange plots divided in three areas along the y-axis: Red: subsecond scale exchange, invisible in this assay, blue: visible exchange events, green: sites remaining non-exchanged in this time-frame. Four horizontal bars indicate remarkable site numbers: total number of exchangeable sites (solid), total number of imino and amino protons in tetrads (12 per tetrads; dashed), total number of H-bonds (8 per tetrads; dot-dashed), and total number of imino protons in tetrads (4 per tetrads; dotted)*

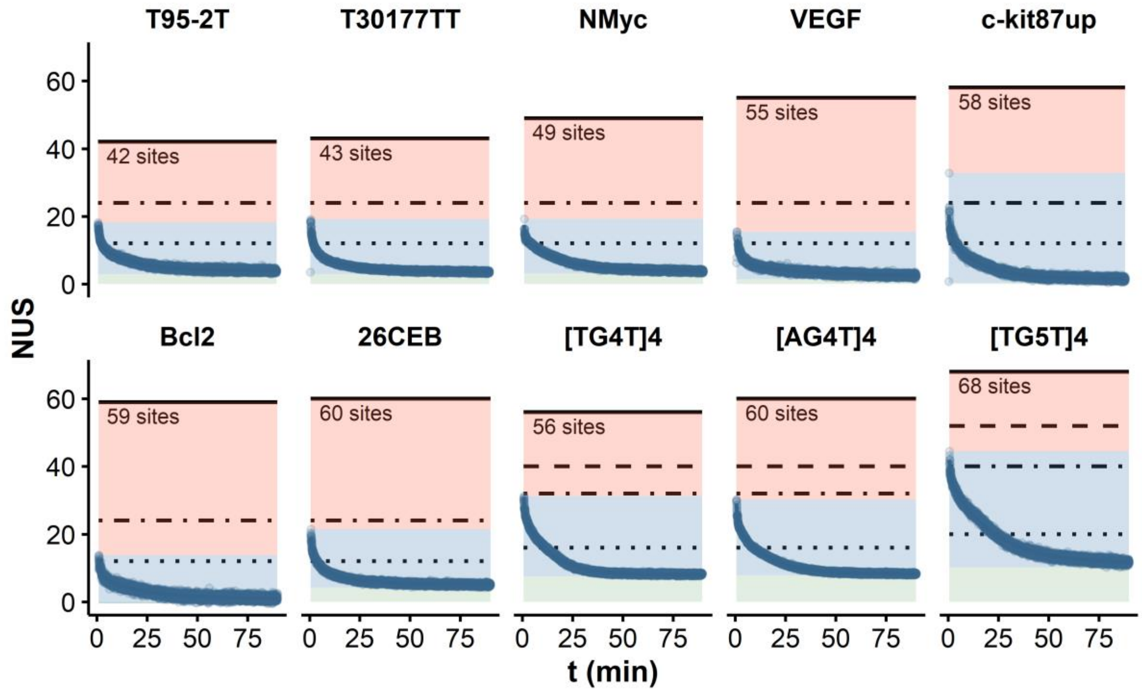


***Figure S16:*** *Exchange plots (t < 5 min) divided in three areas along the y-axis: Red: subsecond scale exchange, invisible in this assay, blue: visible exchange events, green: sites remaining non-exchanged in this time-frame. Four horizontal bars indicate remarkable site numbers: total number of exchangeable sites (solid), total number of imino and amino protons in tetrads (12 per tetrads; dashed), total number of H-bonds (8 per tetrads; dot-dashed), and total number of imino protons in tetrads (4 per tetrads; dotted)*

### Non-linear fitting of the HDX/MS exchange data (real time)

***Table S4:*** *Double exponential fitting parameters (real-time MS)*

| Analyte | | Estimates | | | | | Standard Errors | | | | | Significance (Pr(>\|t\|)) | | | | |
|---|---|---|---|---|---|---|---|---|---|---|---|---|---|---|---|---|
| Oligonucleotide | Tetrads | $N_1$ | $k_1$ ($min^{-1}$) | $N_2$ | $k_2$ ($min^{-1}$) | $NUS_\infty$ | $N_1$ | $k_1$ ($min^{-1}$) | $N_2$ | $k_2$ ($min^{-1}$) | $NUS_\infty$ | $N_1$ | $k_1$ ($min^{-1}$) | $N_2$ | $k_2$ ($min^{-1}$) | $NUS_\infty$ |
| $[TG_5T]_4$ | 5 | 16.1 | 1.675 | 25.7 | 0.047 | 11.3 | 1.7 | 0.117 | 0.1 | 0.000 | 0.0 | *** | *** | *** | *** | *** |
| $[AG_4T]_4$ | 4 | 16.6 | 1.488 | 14.5 | 0.066 | 8.3 | 0.4 | 0.031 | 0.0 | 0.000 | 0.0 | *** | *** | *** | *** | *** |
| $[TG_4T]_4$ | 4 | 21.1 | 2.169 | 18.7 | 0.078 | 8.1 | 1.7 | 0.095 | 0.0 | 0.000 | 0.0 | *** | *** | *** | *** | *** |
| 26CEB | 3 | 10.6 | 0.644 | 7.1 | 0.073 | 5.2 | 0.2 | 0.017 | 0.1 | 0.001 | 0.0 | *** | *** | *** | *** | *** |
| Bcl2 | 3 | 11.5 | 0.853 | 7.0 | 0.048 | 1.0 | 0.8 | 0.053 | 0.1 | 0.001 | 0.0 | *** | *** | *** | *** | *** |
| NMyc | 3 | 182.7 | 4.373 | 10.4 | 0.059 | 3.9 | 55.0 | 0.302 | 0.0 | 0.000 | 0.0 | *** | *** | *** | *** | *** |
| T30177TT | 3 | 9.4 | 0.566 | 6.0 | 0.063 | 3.7 | 0.1 | 0.013 | 0.1 | 0.001 | 0.0 | *** | *** | *** | *** | *** |
| T95-2T | 3 | 12.0 | 0.959 | 7.8 | 0.062 | 4.0 | 0.4 | 0.029 | 0.0 | 0.001 | 0.0 | *** | *** | *** | *** | *** |
| VEGF | 3 | 8.9 | 0.438 | 4.2 | 0.035 | 2.4 | 0.2 | 0.014 | 0.1 | 0.001 | 0.0 | *** | *** | *** | *** | *** |
| c-kit87up | 3 | 9.3 | 0.866 | 12.3 | 0.063 | 1.6 | 0.4 | 0.046 | 0.1 | 0.001 | 0.0 | *** | *** | *** | *** | *** |

***Table S5:*** *Single exponential fitting parameters (real-time MS)*

| Analyte | | Estimates | | | Standard Errors | | | Significance (Pr(>\|t\|)) | | |
|---|---|---|---|---|---|---|---|---|---|---|
| Oligonucleotide | Tetrads | $N_1$ | $k_1$ ($min^{-1}$) | $NUS_\infty$ | $N_1$ | $k_1$ ($min^{-1}$) | $NUS_\infty$ | $N_1$ | $k_1$ ($min^{-1}$) | $NUS_\infty$ |
| $[TG_5T]_4$ | 5 | 26.3 | 0.049 | 11.5 | 0.1 | 0.000 | 0.0 | *** | *** | *** |
| $[AG_4T]_4$ | 4 | 15.8 | 0.074 | 8.4 | 0.0 | 0.000 | 0.0 | *** | *** | *** |
| $[TG_4T]_4$ | 4 | 19.5 | 0.082 | 8.1 | 0.0 | 0.000 | 0.0 | *** | *** | *** |
| 26CEB | 3 | 11.0 | 0.118 | 5.4 | 0.1 | 0.001 | 0.0 | *** | *** | *** |
| Bcl2 | 3 | 8.3 | 0.062 | 1.1 | 0.1 | 0.001 | 0.0 | *** | *** | *** |
| NMyc | 3 | 10.6 | 0.061 | 3.9 | 0.0 | 0.000 | 0.0 | *** | *** | *** |
| T30177TT | 3 | 10.0 | 0.114 | 3.9 | 0.1 | 0.001 | 0.0 | *** | *** | *** |
| T95-2T | 3 | 9.5 | 0.080 | 4.1 | 0.0 | 0.001 | 0.0 | *** | *** | *** |
| VEGF | 3 | 7.0 | 0.082 | 2.9 | 0.1 | 0.001 | 0.0 | *** | *** | *** |
| c-kit87up | 3 | 14.2 | 0.076 | 1.8 | 0.1 | 0.001 | 0.0 | *** | *** | *** |

***Table S6:*** *Model selection parameters (real-time MS): lower Akaike information criterion (AIC) and Bayesian information criterion (BIC) indicate which model describe best the experimental data.*

| **Data** | | **Double exponential (j = 2)** | | | | | **Single exponential (j = 1)** | | | | |
|---|---|---|---|---|---|---|---|---|---|---|---|
| **Oligonucleotide** | **n** | **Degrees freedom** | **Sigma** | **Log-likelihood** | **AICc** | **BIC** | **Degrees freedom** | **Sigma** | **Log-likelihood** | **AICc** | **BIC** |
| $[TG_5T]_4$ | 3,927 | 3,922 | 0.64 | -3,825.22 | 7,662 | 7,700 | 3,924 | 0.70 | -4,197.73 | 8,403 | 8,429 |
| $[AG_4T]_4$ | 3,930 | 3,925 | 0.23 | 162.60 | -313 | -276 | 3,927 | 0.42 | -2,165.82 | 4,340 | 4,365 |
| $[TG_4T]_4$ | 3,899 | 3,894 | 0.32 | -1,079.66 | 2,171 | 2,209 | 3,896 | 0.40 | -1,916.81 | 3,842 | 3,867 |
| 26CEB | 3,928 | 3,923 | 0.34 | -1,297.59 | 2,607 | 2,645 | 3,925 | 0.49 | -2,735.80 | 5,480 | 5,505 |
| Bcl2 | 3,904 | 3,899 | 0.62 | -3,656.90 | 7,326 | 7,363 | 3,901 | 0.69 | -4,091.07 | 8,190 | 8,215 |
| NMyc | 3,915 | 3,910 | 0.28 | -516.28 | 1,045 | 1,082 | 3,912 | 0.30 | -864.13 | 1,736 | 1,761 |
| T30177TT | 3,910 | 3,905 | 0.31 | -1,022.06 | 2,056 | 2,094 | 3,907 | 0.49 | -2,748.85 | 5,506 | 5,531 |
| T95-2T | 4,999 | 4,994 | 0.36 | -2,041.14 | 4,094 | 4,133 | 4,996 | 0.49 | -3,553.32 | 7,115 | 7,141 |
| VEGF | 3,905 | 3,900 | 0.43 | -2,276.45 | 4,565 | 4,603 | 3,902 | 0.57 | -3,332.05 | 6,672 | 6,697 |
| c-kit87up | 3,852 | 3,847 | 0.62 | -3,627.63 | 7,267 | 7,305 | 3,849 | 0.72 | -4,187.84 | 8,384 | 8,409 |

#### Exchange trends, cation binding specificity and number of tetrads

In native MS, the number of tetrads can be determined from the number of specifically-bound cations, assuming that $n$ tetrads specifically bind $n-1$ cations.[5] A typical hurdle in this approach is the concomitant binding of non-specific (diffusely bound) cations, resulting in a convoluted distribution of cation stoichiometries that does not directly reflect that in solution. With oligonucleotides of unknown conformation, one can tentatively subtract the contribution of non-specific adducts using the distribution of an unfolded oligonucleotide control.[5] This assumes that the amount of non-specific binding to the control reflects that of the G4, which is true to some extent if the control is properly designed, but is not necessarily trivial. However, non-specific cations do not alter HDX rates,[6] and therefore HDX/MS provides the number of specific cation(s) simply by comparing the exchange kinetics of different adduct stoichiometries

For instance, 22AG has four repeats of three guanines, which intuitively should form a G4 with three tetrads. However, the native MS spectrum of 22AG in 1 mM KCl solution presents a wide distribution of $K^+$ (Figure S**17**A), suggesting that it could also form G4s with smaller or larger number of tetrads. In HDX/MS experiments, the 1-$K^+$ and 2-$K^+$ species exchange at significantly different rates and therefore originate from different conformers in solutions (Figure S**17**B). In contrast, the exchange plot of 22AG·3$K^+$ is superimposed on that of 22AG·2$K^+$, indicating that they both originate from the same conformer in solution. HDX/MS thus clearly shows that 22AG folds into a mixture of 1-$K^+$ (2-tetrads) and 2-$K^+$ (3-tetrads) conformers in 1 mM KCl solutions, in a single experiment without the need for a control.

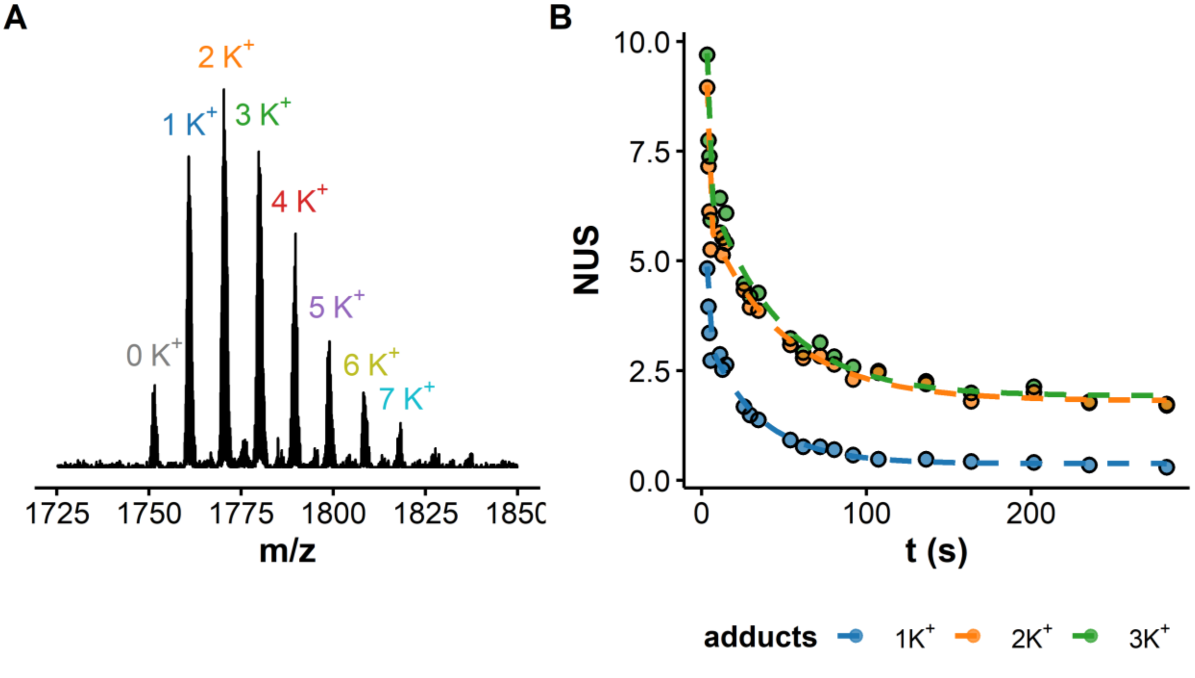


***Figure S17:*** *Cation specificity and HDX rates: A. Native ESI-MS spectrum of 22AG (4- charge state) where the most abundant potassium stoichiometries are annotated, B. Corresponding continuous-flow HDX/MS of the 1—3 $K^+$ species. The lines are the results of non-linear fitting with equation (4) , where j = 2.*

#### NUS vs. $T_m$

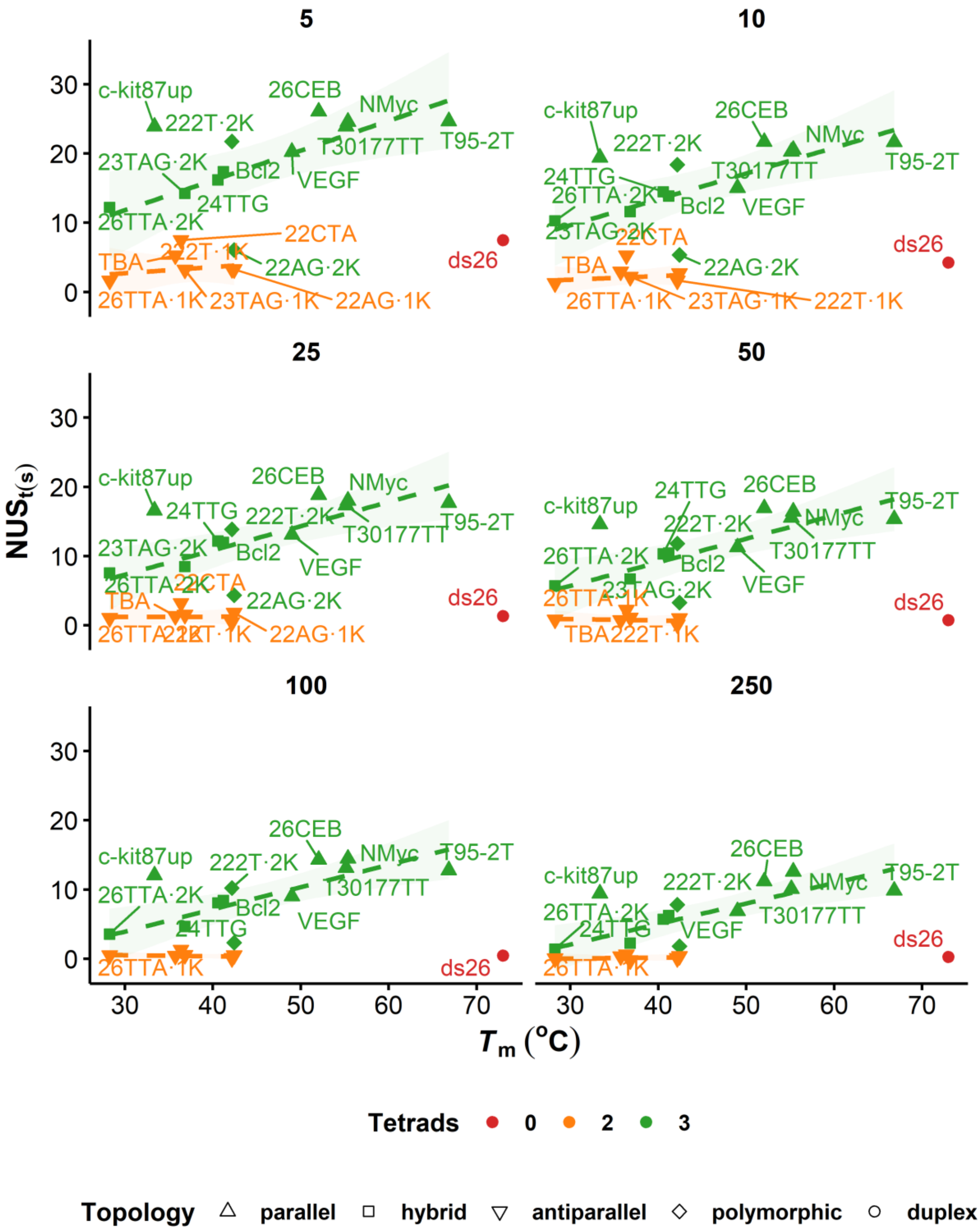


*Figure S18:* *Relation between $T_m$ and $NUS_t$, measured by continuous-flow HDX/MS (panels are labelled with the mixing time in seconds). The straight lines and 95% confidence intervals (colored ribbon) are the result of linear fitting.*

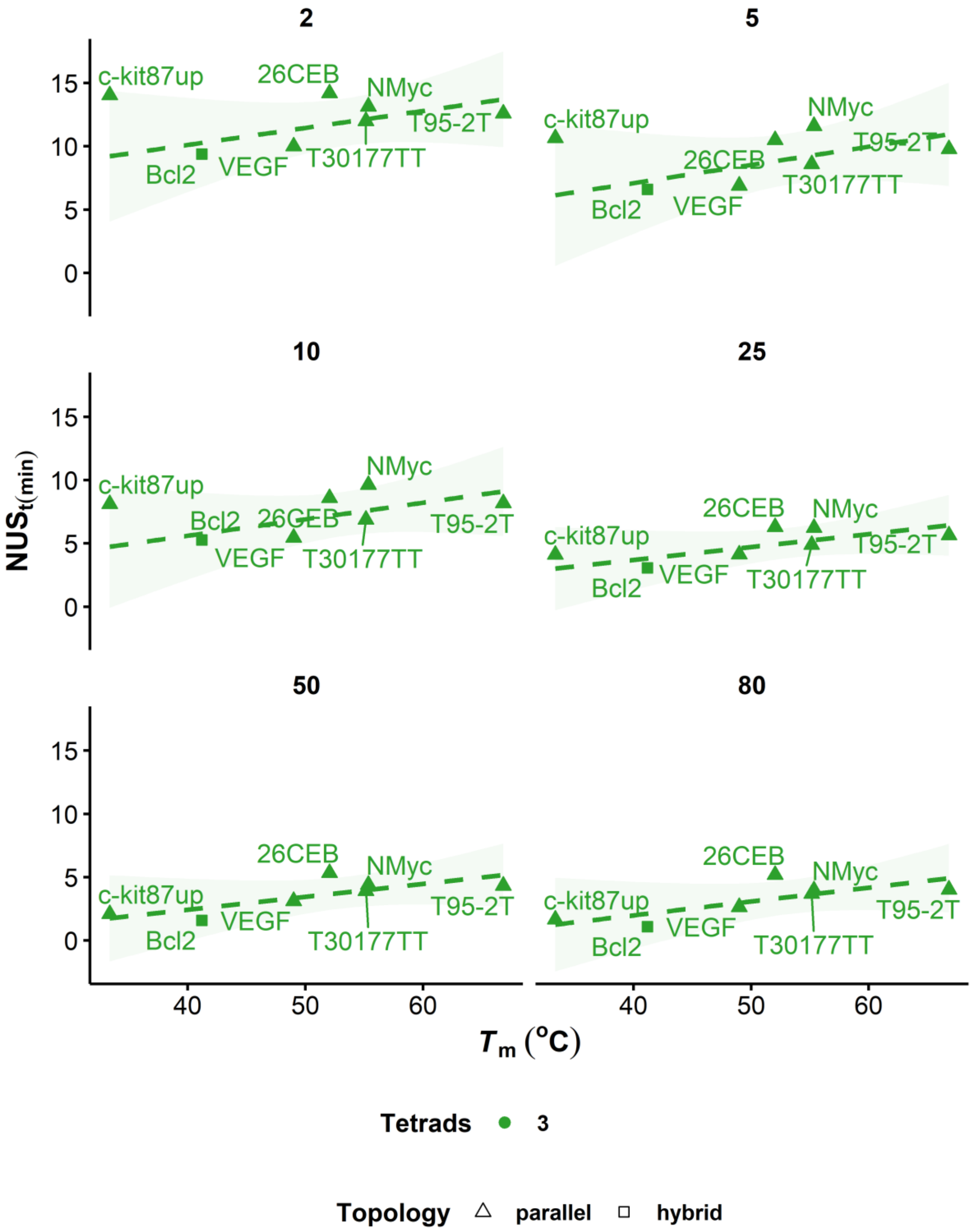


***Figure S19:*** *Relation between $T_m$ and $NUS_t$, measured by real-time HDX/MS (panels are labelled with the mixing time in minutes). The straight lines and 95% confidence intervals (colored ribbon) are the result of linear fitting.*

#### NUS vs. $\Delta G^{0}_{22°C}$

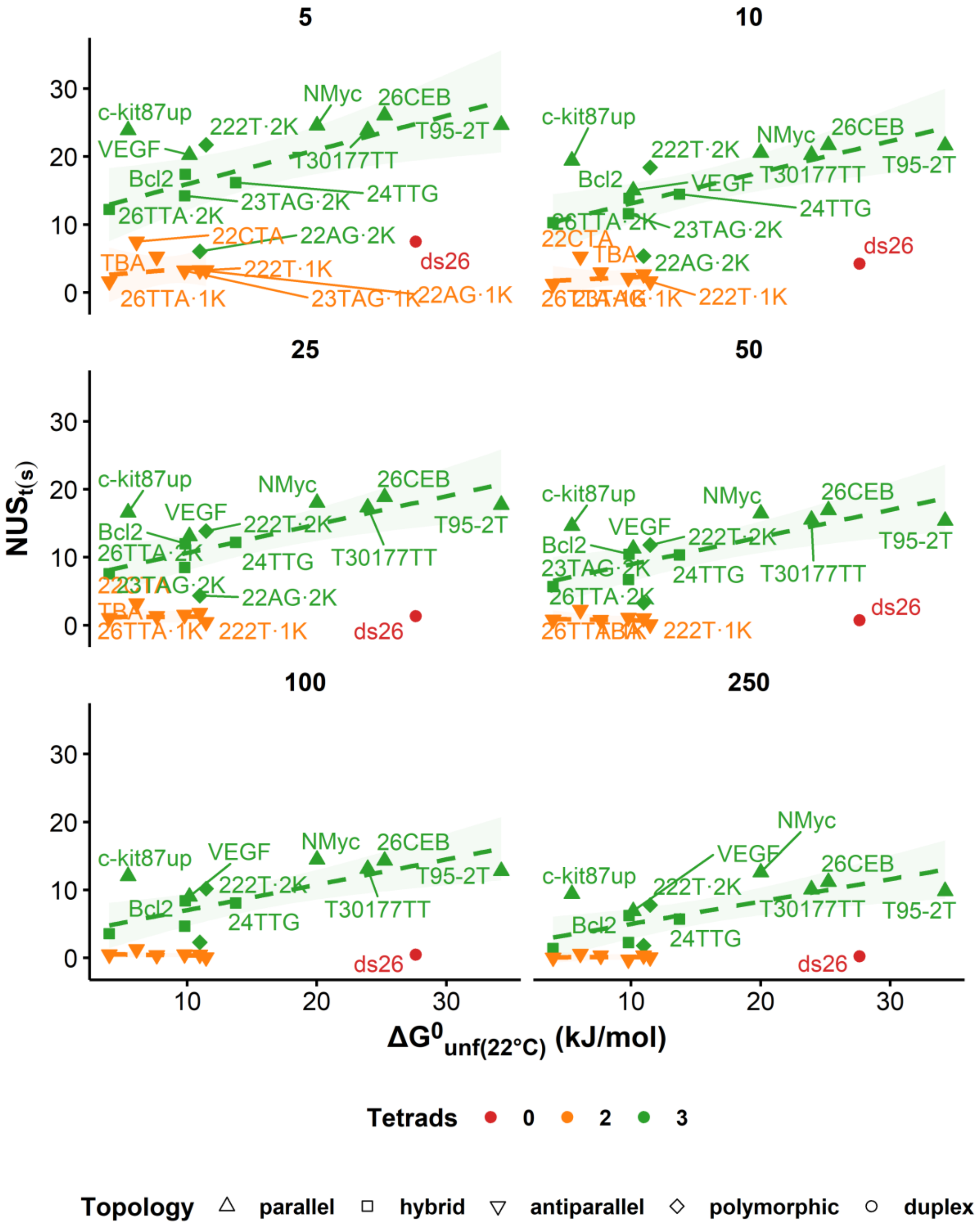


***Figure S20:*** *Relation between the apparent free energy of unfolding at 22°C and $NUS_t$, measured by continuous-flow HDX/MS (panels are labelled with the mixing time in seconds). The straight lines and 95% confidence intervals (colored ribbon) are the result of linear fitting.*

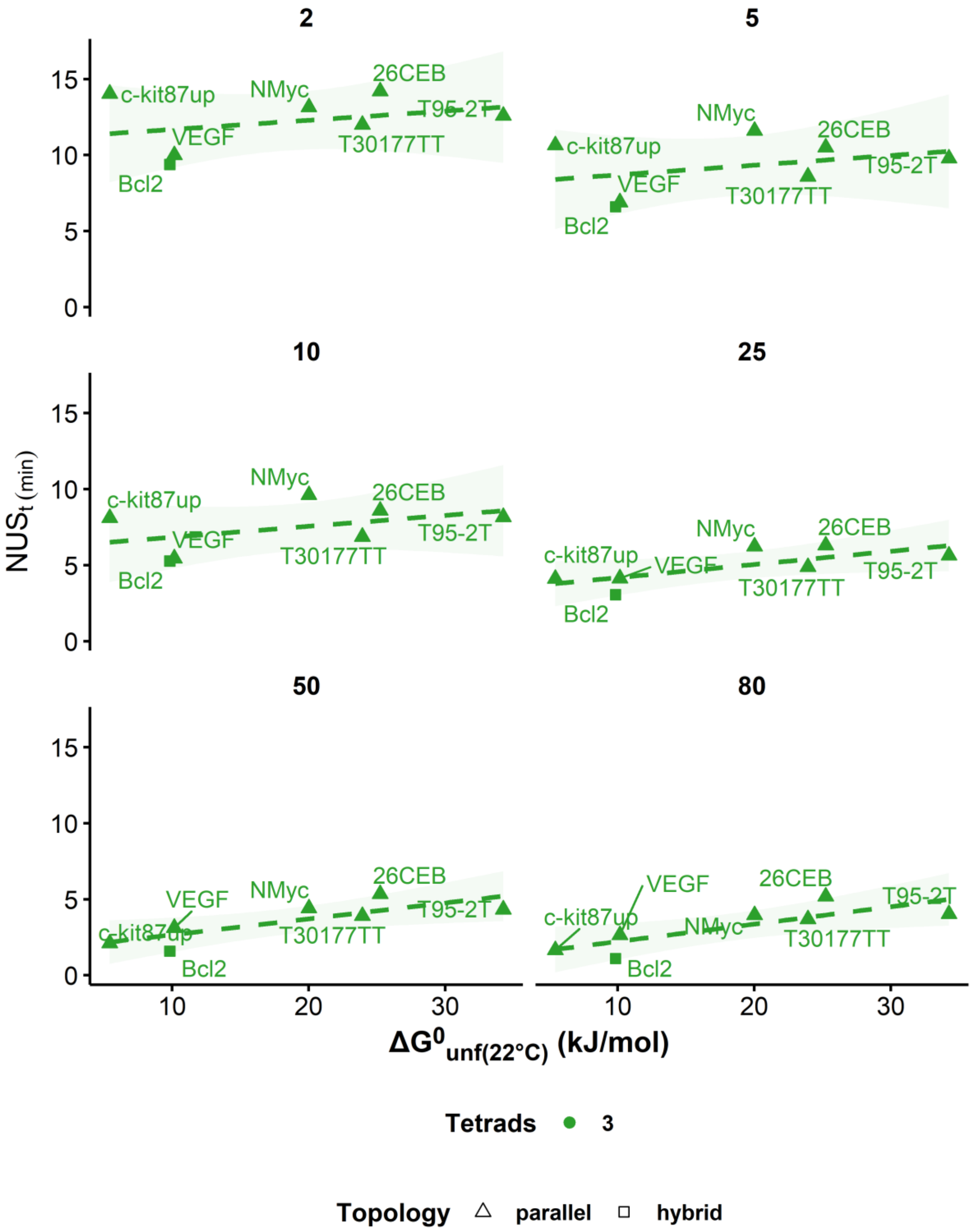


*Figure S21: Relation between the apparent free energy of unfolding at 22°C and $NUS_t$, measured by real-time HDX/MS (panels are labelled with the mixing time in minutes). The straight lines and 95% confidence intervals (colored ribbon) are the result of linear fitting.*

#### T95-2T structure and stability

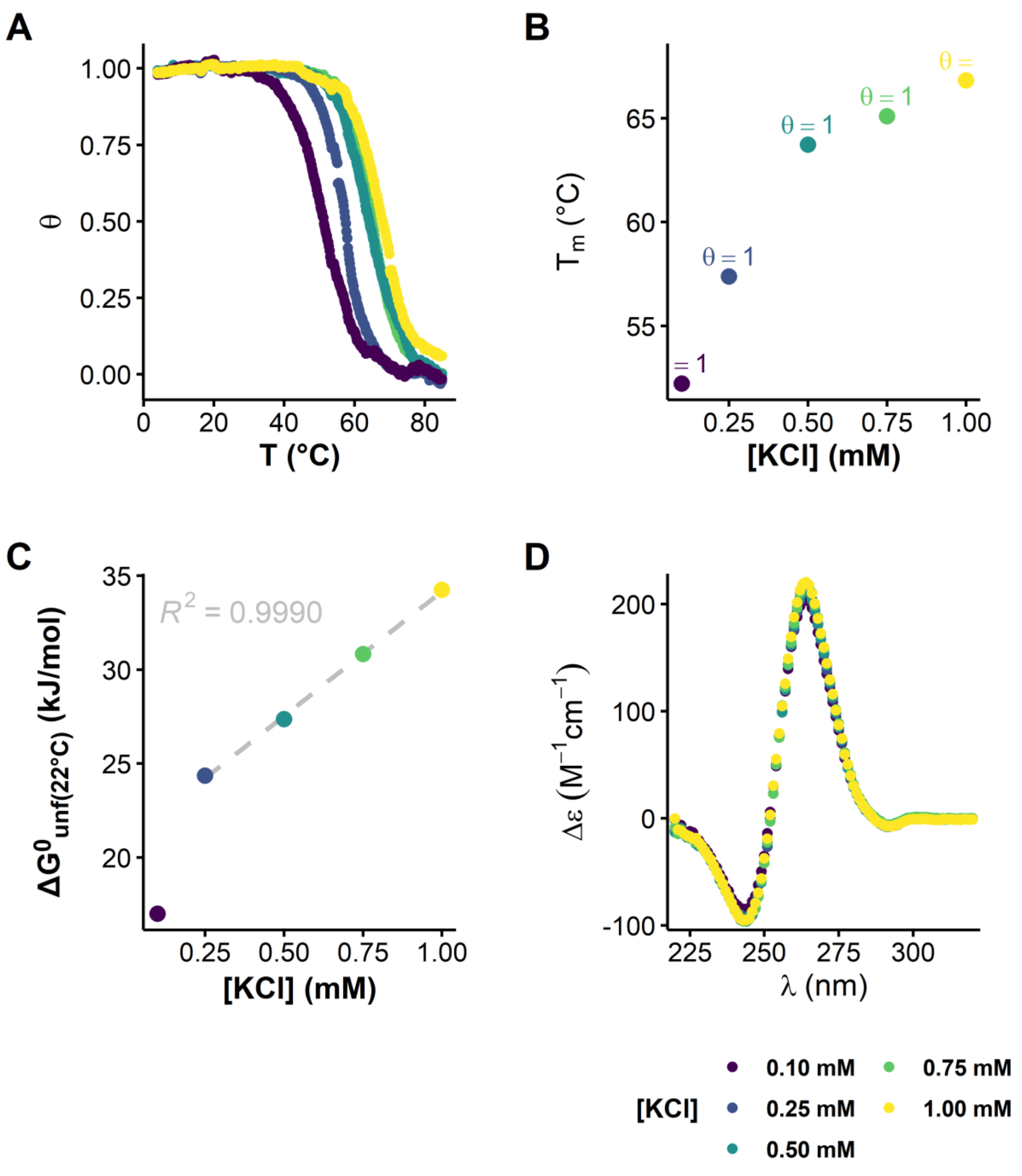


*__Figure S22:__ Structure and stability of T95-2T (10 µM) in presence of various KCl concentrations: A. Processed UV-melting data showing the folded fraction as a function of the solution temperature (heating ramp), B. Melting temperatures (labeled with the corresponding folded fraction) and C. Apparent free energy of unfolding as a function of KCl concentration (the grey dashed line is the result of linear fitting), and D. Circular dichroism spectra obtained for the same solutions.*

### Deconvolution of multimodal isotopic distributions

#### Pure EX2 species

### T30177-TT

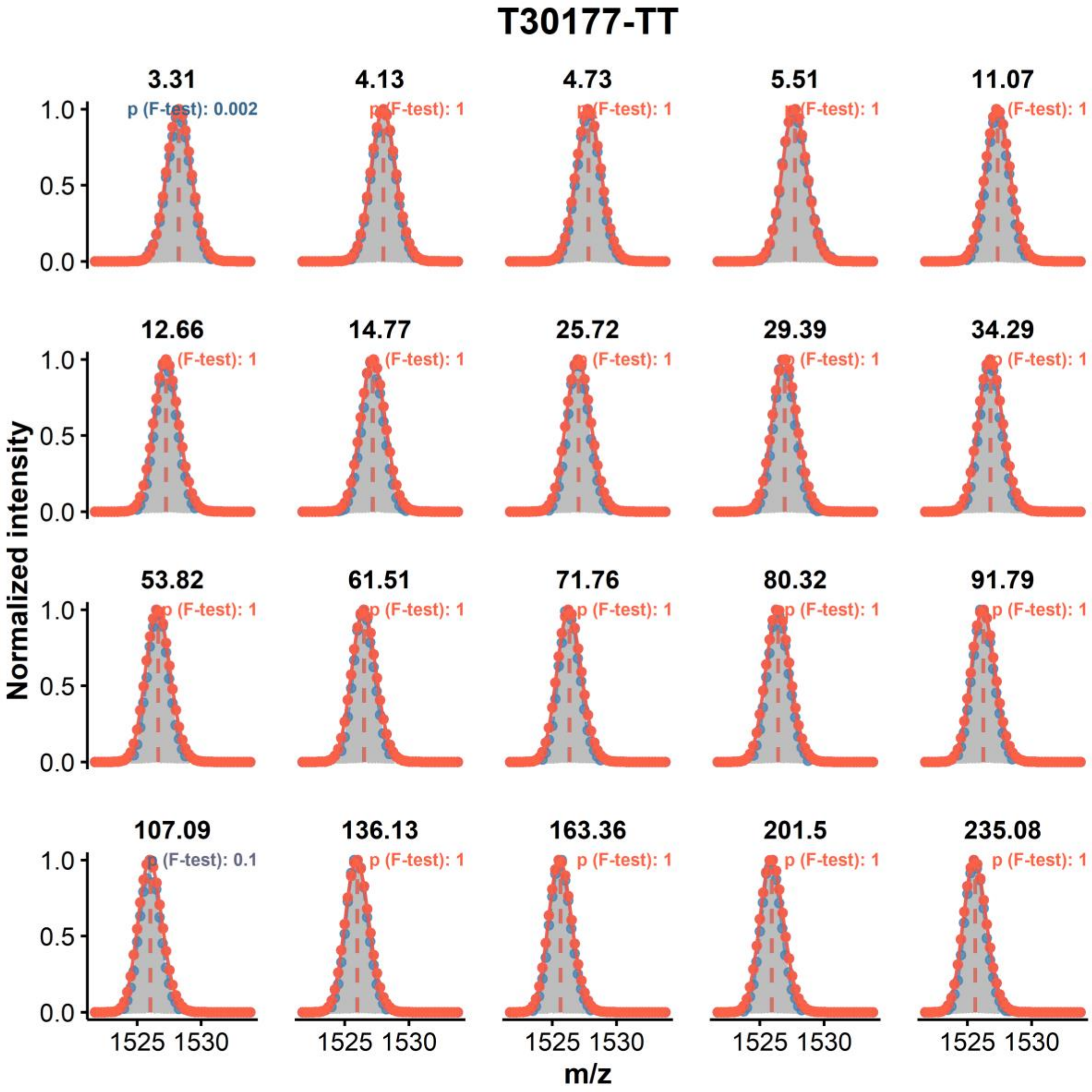


*Figure S23: Monomodal distribution modeling on the peak-picked, continuous-flow exchange data of T30177-TT (z = 4-, K = 2, initial DC = 90%, final DC = 9%). The experimental data is shown in grey, the peak picking in blue, the fit in orange. The orange vertical dashed lines show the position of their respective centroids, the blue one being the apparent experimental value (when not visible: superimposed with the orange line)*

#### 26CEB

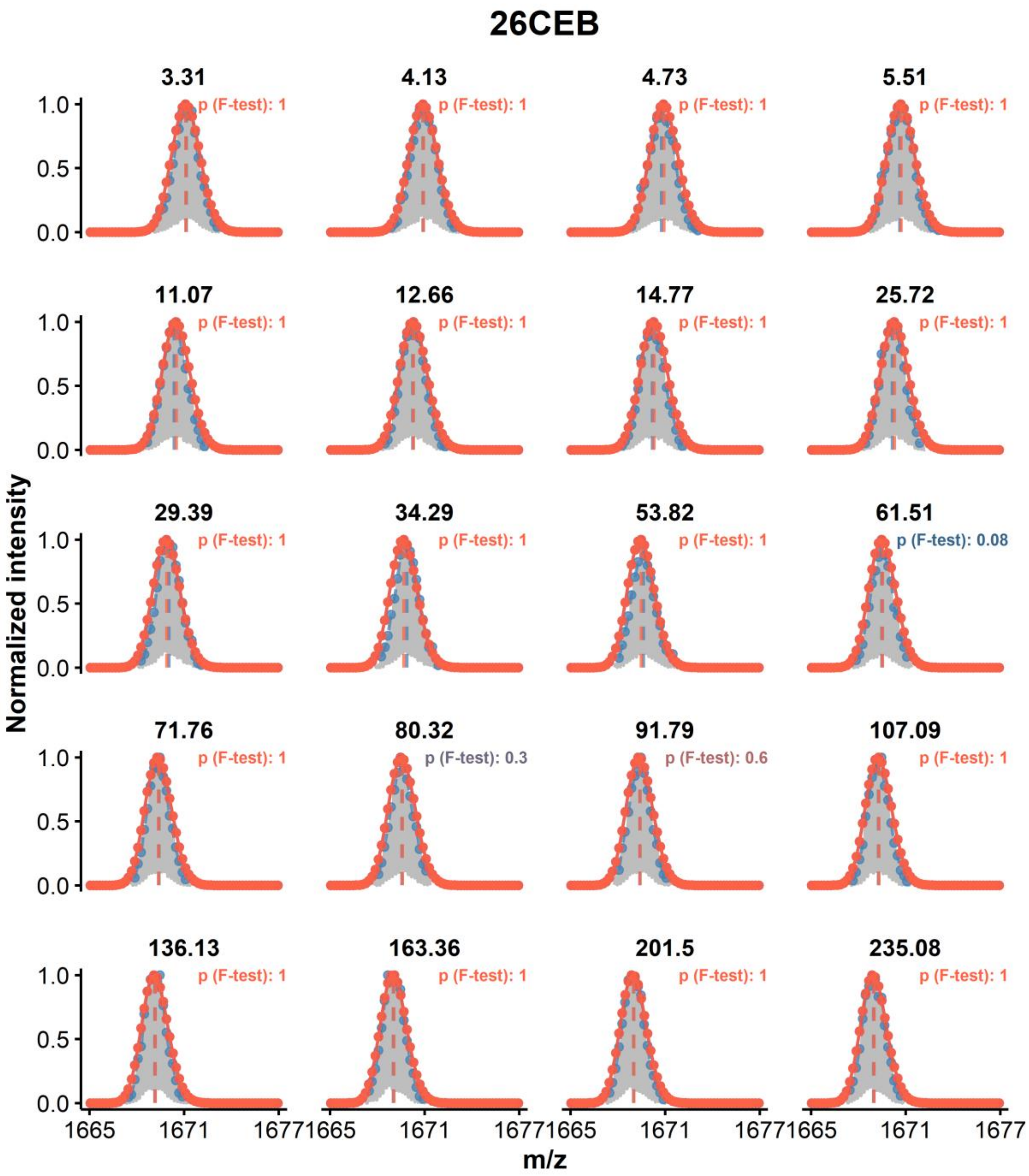


***Figure S24:*** *Monomodal distribution modeling on the peak-picked, continuous-flow exchange data of 26CEB (z = 5-, K = 2, initial DC = 90%, final DC = 9%). The experimental data is shown in grey, the peak picking in blue, the fit in orange. The orange vertical dashed lines show the position of their respective centroids, the blue one being the apparent experimental value (when not visible: superimposed with the orange line)*

## T95-2T

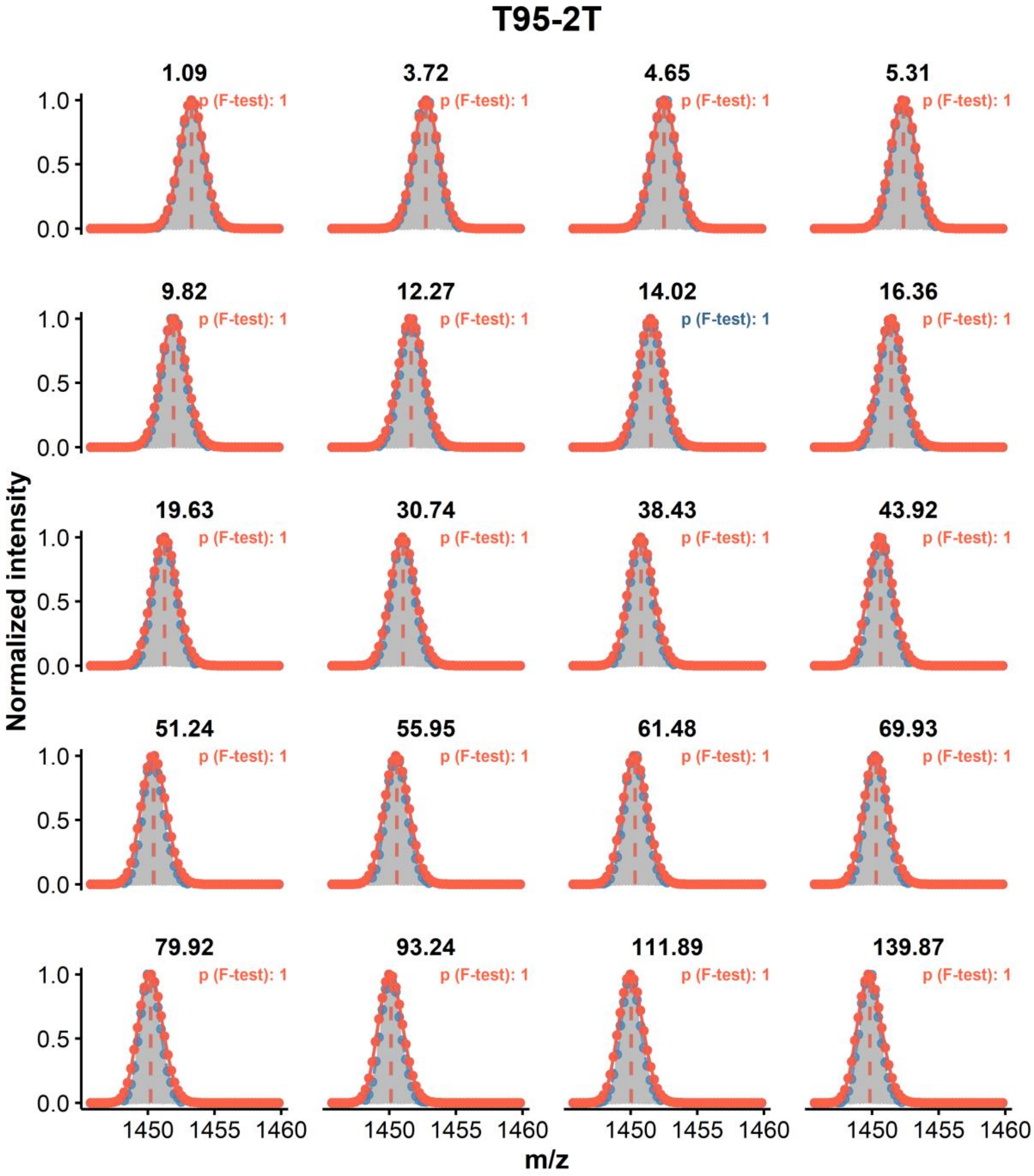


***Figure S25:*** *Monomodal distribution modeling on the peak-picked, continuous-flow exchange data of T95-2T (z = 4-, K = 2, initial DC = 90%, final DC = 9%). The experimental data is shown in grey, the peak picking in blue, the fit in orange. The orange vertical dashed lines show the position of their respective centroids, the blue one being the apparent experimental value (when not visible: superimposed with the orange line)*

#### Nmyc

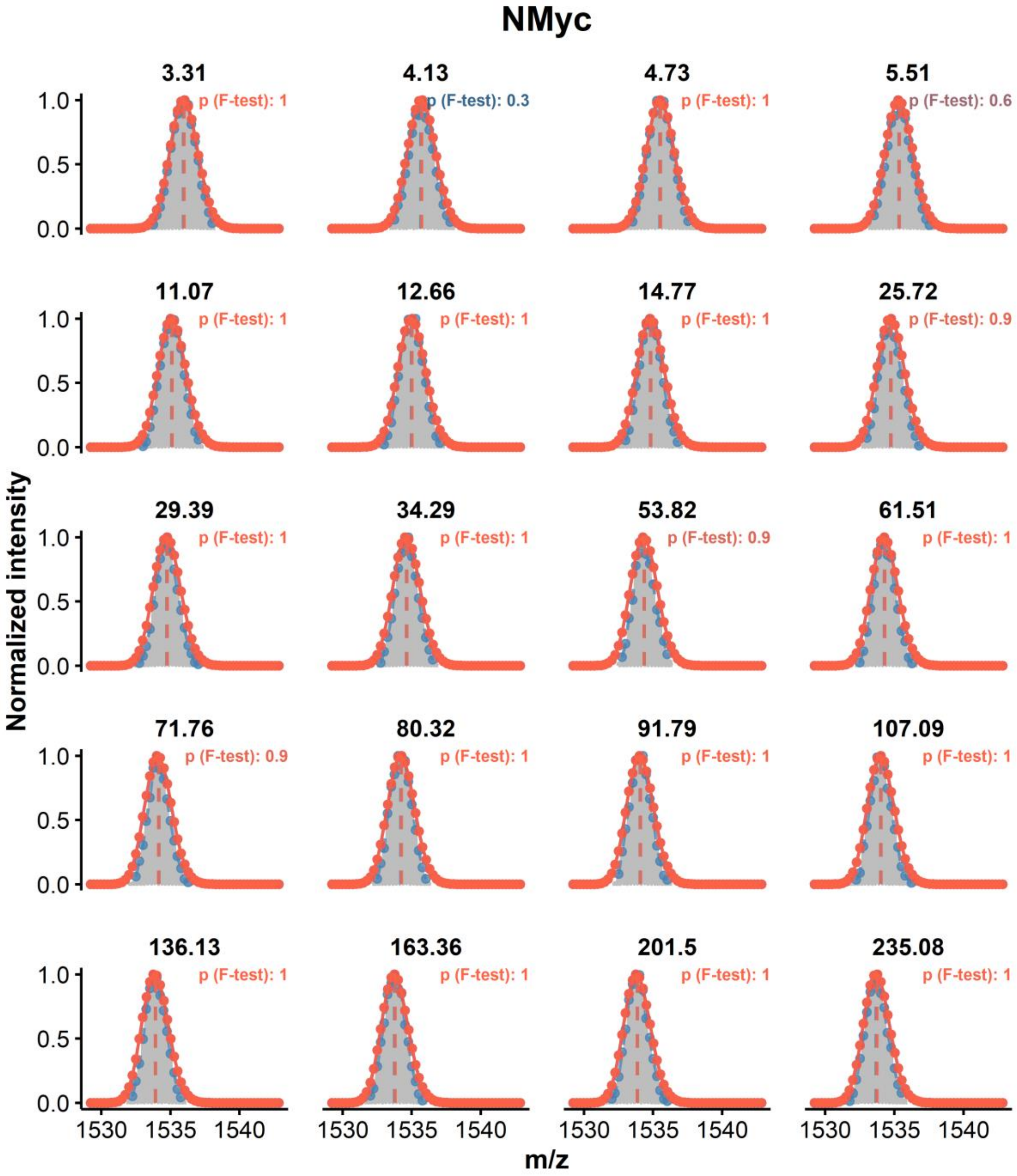


***Figure S26:*** *Monomodal distribution modeling on the peak-picked, continuous-flow exchange data of NMyc (z = 4-, K = 2, initial DC = 90%, final DC = 9%). The experimental data is shown in grey, the peak picking in blue, the fit in orange. The orange vertical dashed lines show the position of their respective centroids, the blue one being the apparent experimental value (when not visible: superimposed with the orange line)*

#### Tetramolecular G4s

The tetramolecular G4s formed by $TG_4T$, $AG_4T$, and $TG_5T$ are in equilibrium with the corresponding unfolded single strands. Although the experiments were performed on samples at equilibrium, wherein the ratio of G4 to single strands is constant (Figure S**27**), the latter could still dynamically substitute protected strands from G4s. Since the unfolded ssDNA exchange fully in the subsecond range, this would yield more extensively exchanged analytes, resulting in multimodal distributions. Yet, the three tetramolecular G4s display unambiguously monomodal distributions (Figures S**28**—S**30**), and therefore exchange exclusively through the EX2 mechanism. This is most likely due to the very large half-life of these quadruplexes (from days to years, depending on the sequence and experimental conditions), not strand substitution over the course of the experiment.

### *$[TG_4T]_4$*

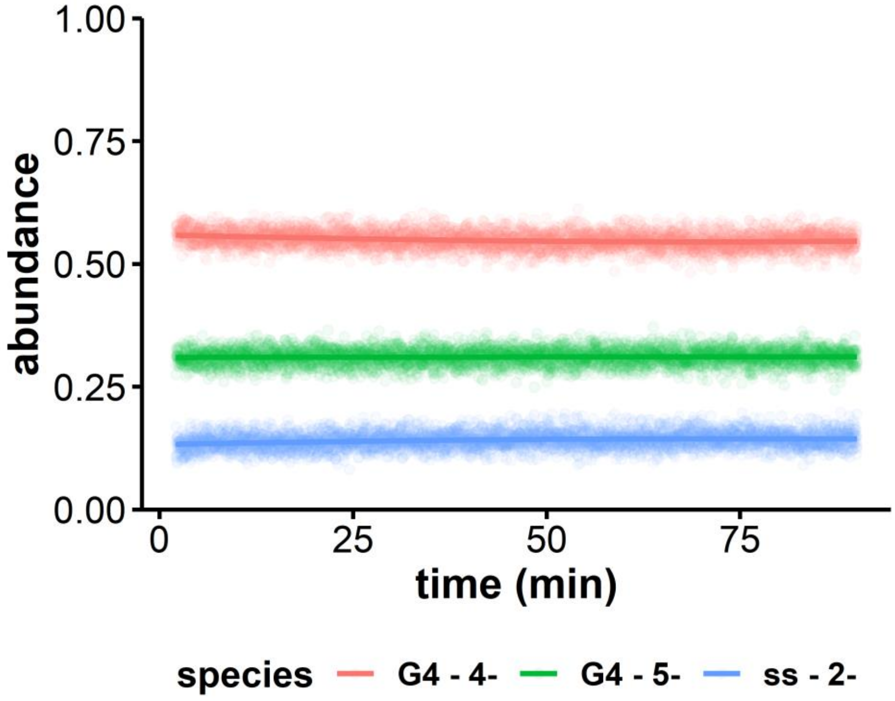


***Figure S27:*** *Relative abundance of the tetramolecular G4 (4- and 5- charge states) and the single strand (2- charge state) during the real-time HDX/MS experiment*

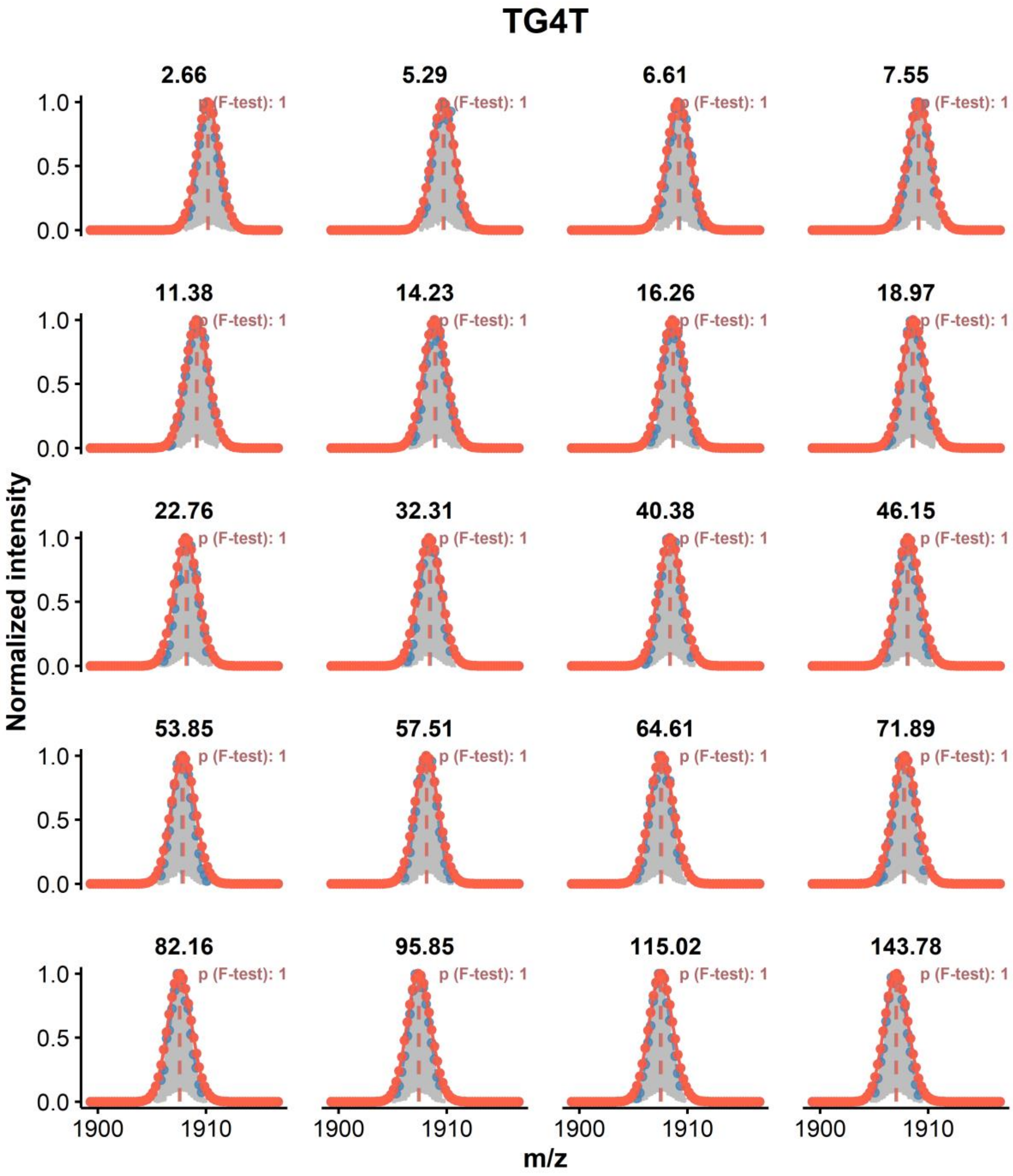


***Figure S28:*** *Monomodal distribution modeling on the peak-picked, continuous-flow exchange data of TG4T (z = 4-, K = 4, initial DC = 90%, final DC = 9%). The experimental data is shown in grey, the peak picking in blue, the fit in orange. The orange vertical dashed lines show the position of their respective centroids, the blue one being the apparent experimental value (when not visible: superimposed with the orange line)*

*[$AG_4T$]$_4$*

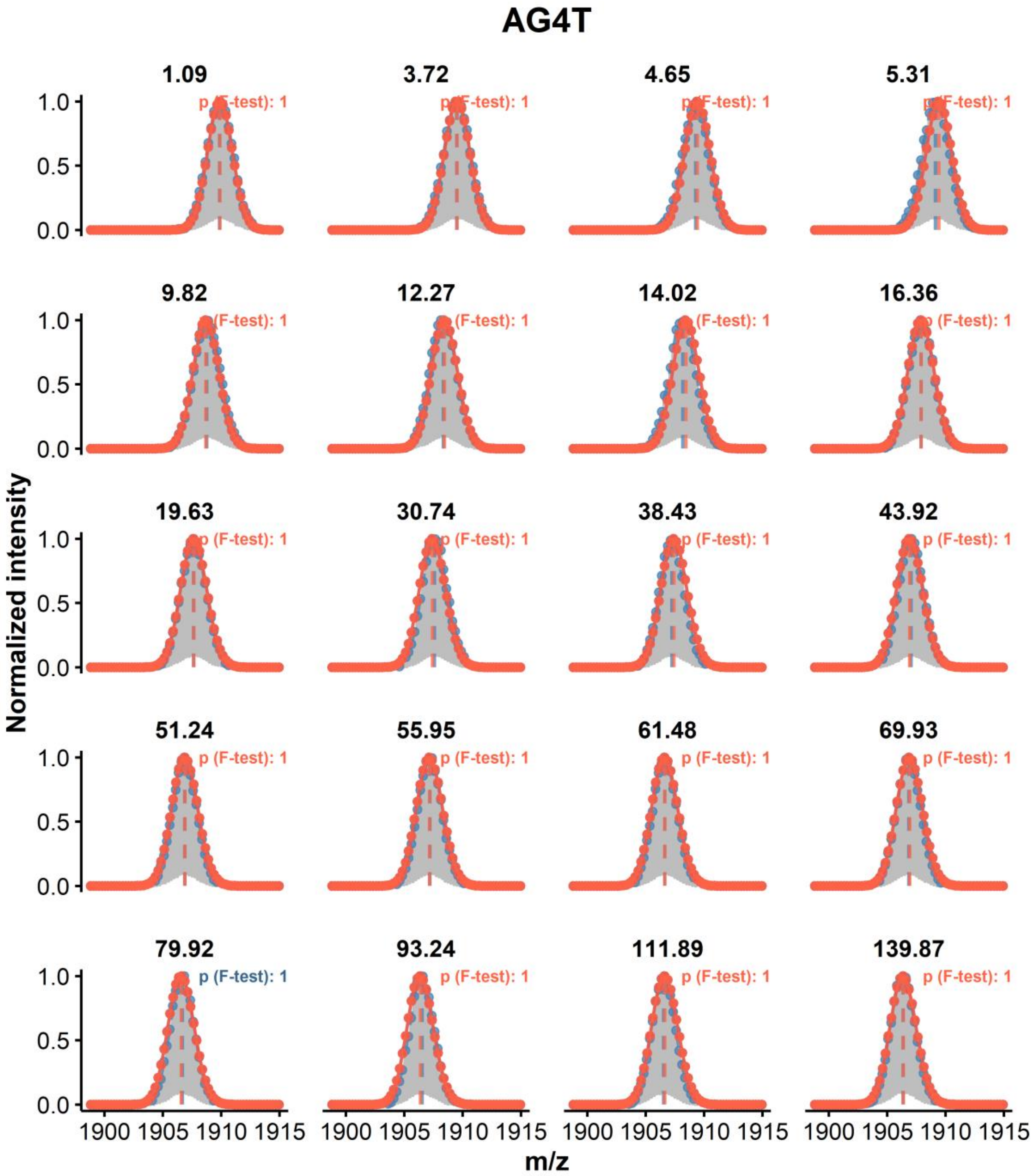


***Figure S29:*** *Monomodal distribution modeling on the peak-picked, continuous-flow exchange data of AG4T (z = 4-, K = 3, initial DC = 90%, final DC = 9%). The experimental data is shown in grey, the peak picking in blue, the fit in orange. The orange vertical dashed lines show the position of their respective centroids, the blue one being the apparent experimental value (when not visible: superimposed with the orange line)*

*[$TG_5T$]$_4$*

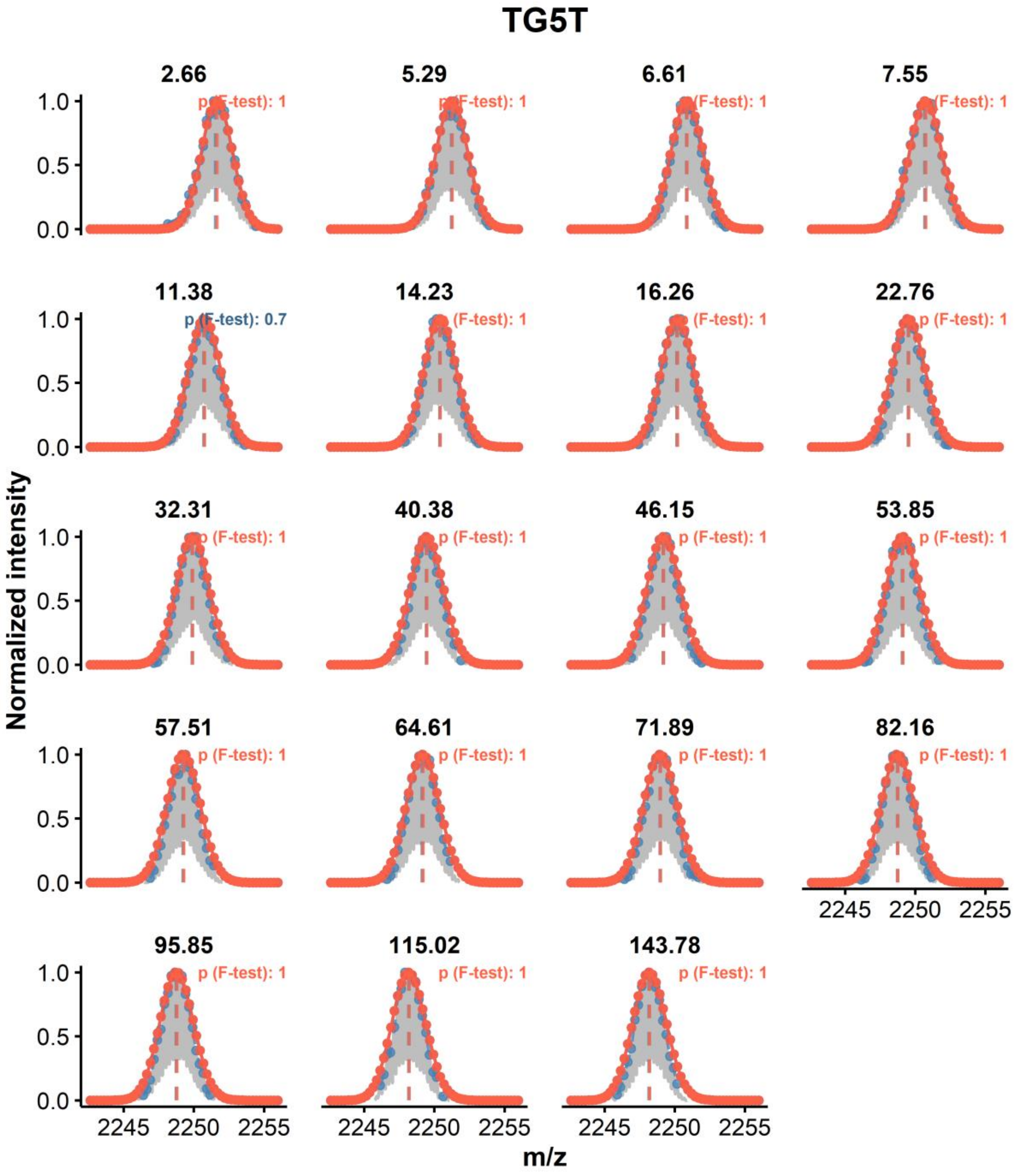


***Figure S30:*** *Monomodal distribution modeling on the peak-picked, continuous-flow exchange data of TG5T (z = 4-, K = 4, initial DC = 90%, final DC = 9%). The experimental data is shown in grey, the peak picking in blue, the fit in orange. The orange vertical dashed lines show the position of their respective centroids, the blue one being the apparent experimental value (when not visible: superimposed with the orange line)*

**ds26**

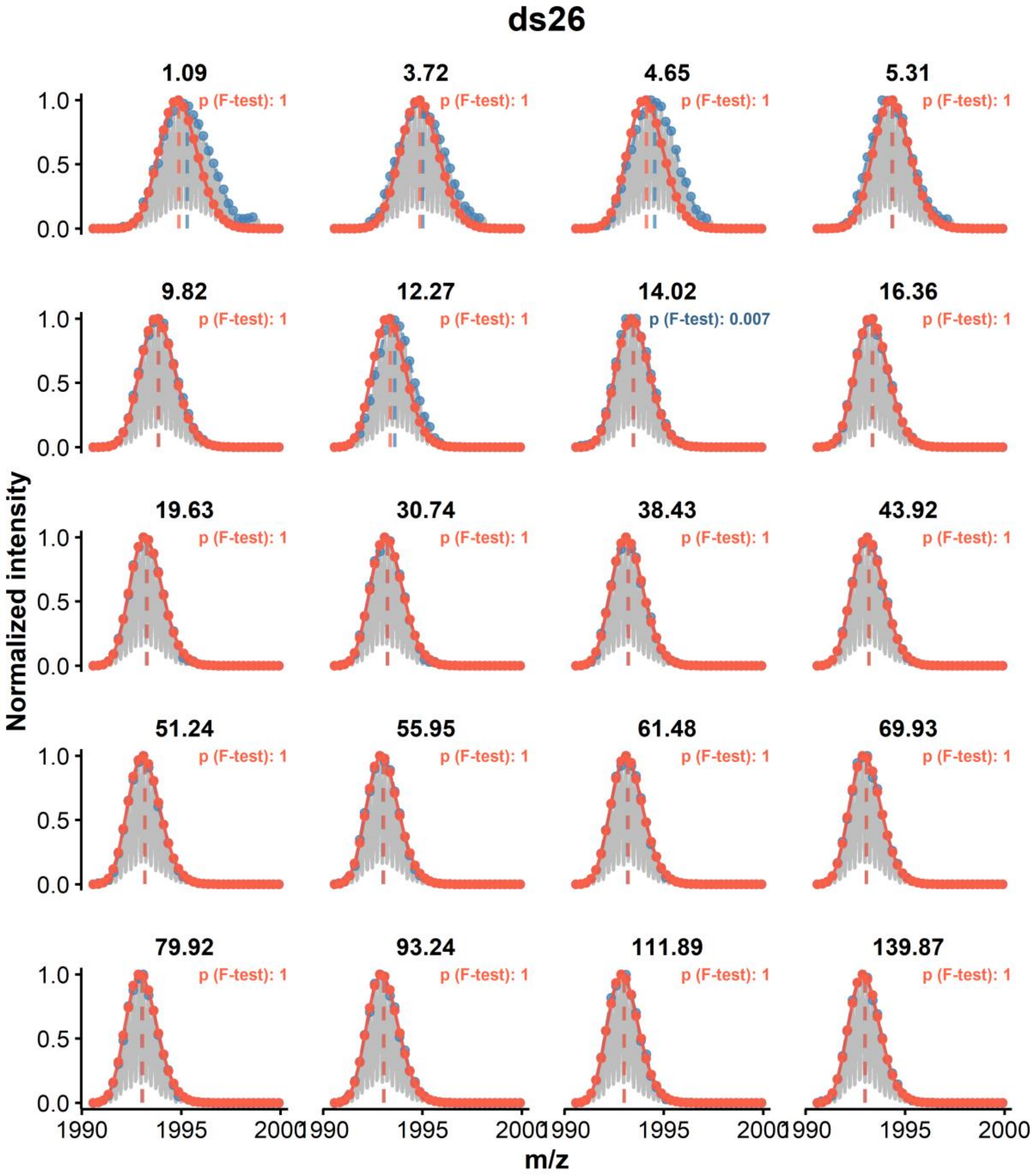


***Figure S31:*** *Monomodal distribution modeling on the peak-picked, continuous-flow exchange data of ds26 (z = 4-, K = 0, initial DC = 90%, final DC = 9%). A. The experimental data is shown in grey, the peak picking in blue, the fit in orange. The orange vertical dashed lines show the position of their respective centroids, the blue one being the apparent experimental value (when not visible: superimposed with the orange line)*

#### Mixed species

##### VEGF

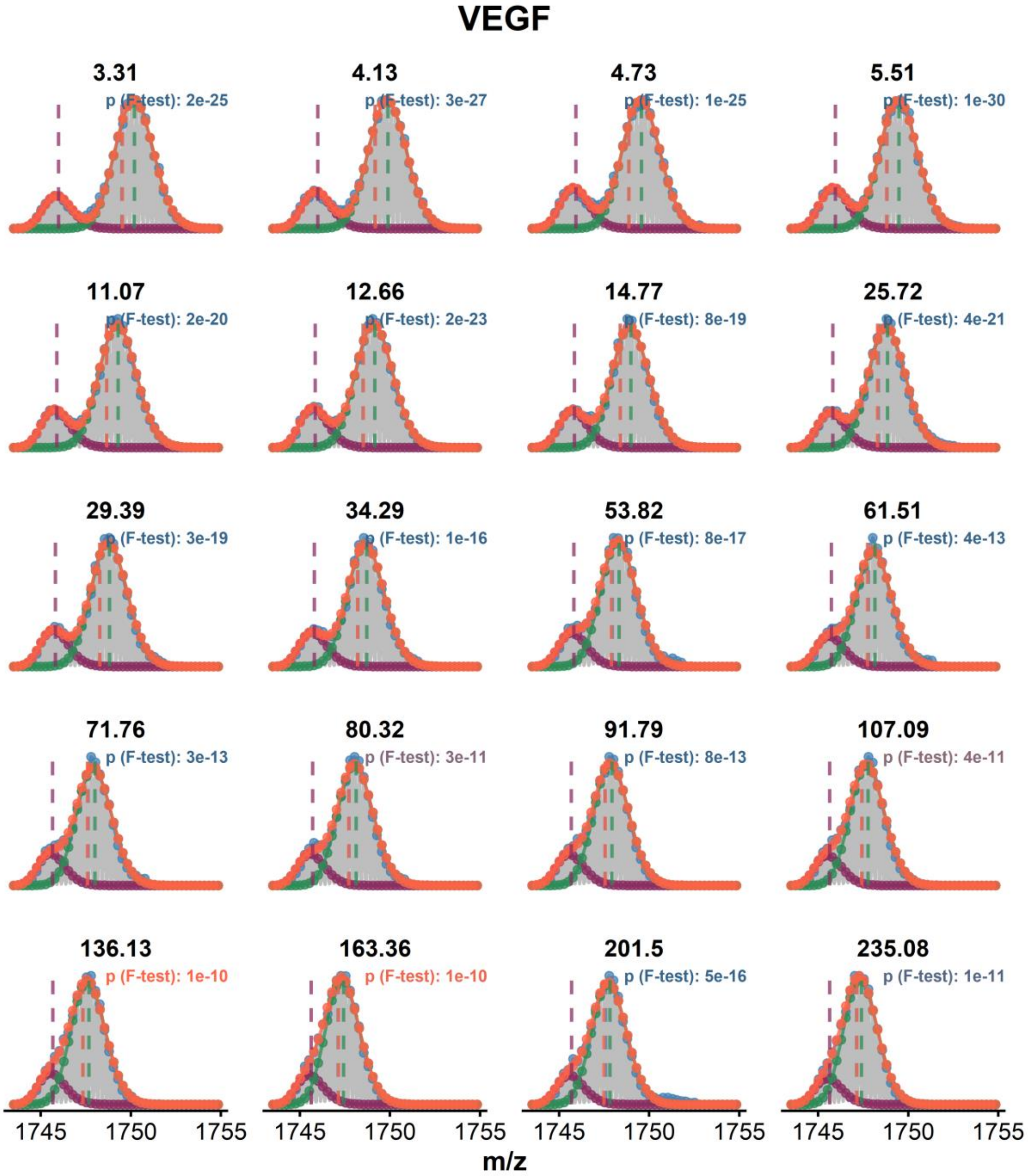


***Figure S32:*** *Bimodal distribution deconvolution on the peak-picked, continuous-flow exchange data of VEGF (z = 4-, K = 2, initial DC = 90%, final DC = 9%). The experimental data is shown in grey, the peak picking in blue, the overall fit in orange, and individual population in green and purple. The vertical dashed lines show the position of their respective centroids, the blue one being the apparent experimental value (when not visible: superimposed with the orange line)*

#### c-kit87up

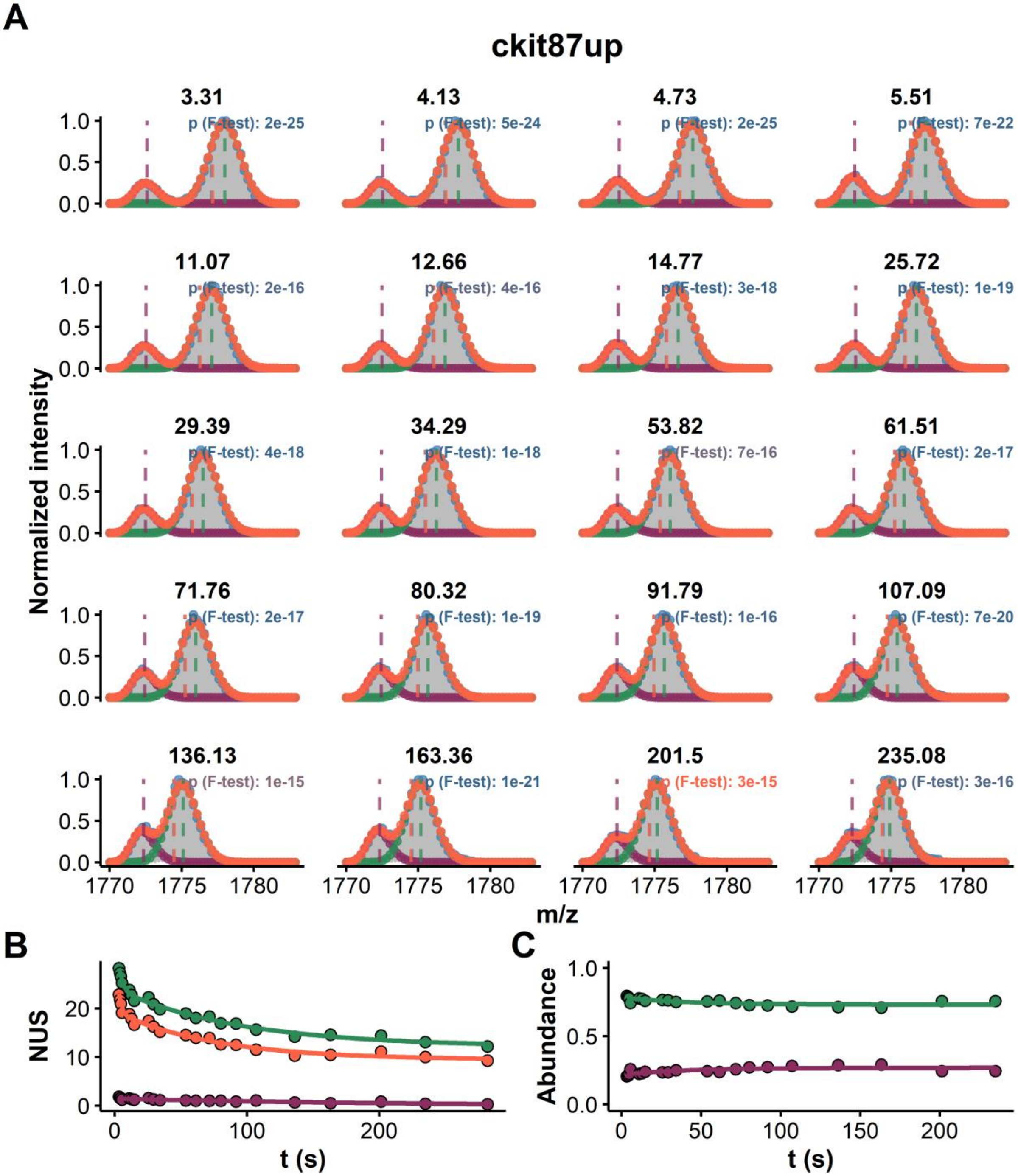


*Figure S33: HDX data modeling and bimodal distribution deconvolution of the continuous-flow exchange data of c-kit87up (z = 4-, K = 2, initial DC = 90%, final DC = 9%). A. The experimental data is shown in grey, the peak picking in blue, the overall fit in orange, and individual population in green and purple. The vertical dashed lines show the position of their respective centroids, the blue one being the apparent experimental value (when not visible: superimposed with the orange line). B. Exchange kinetics of the overall (orange) and deconvoluted distributions (green and purple). C. Relative abundance of the deconvoluted distributions as a function of the exchange time.*

#### Bcl2

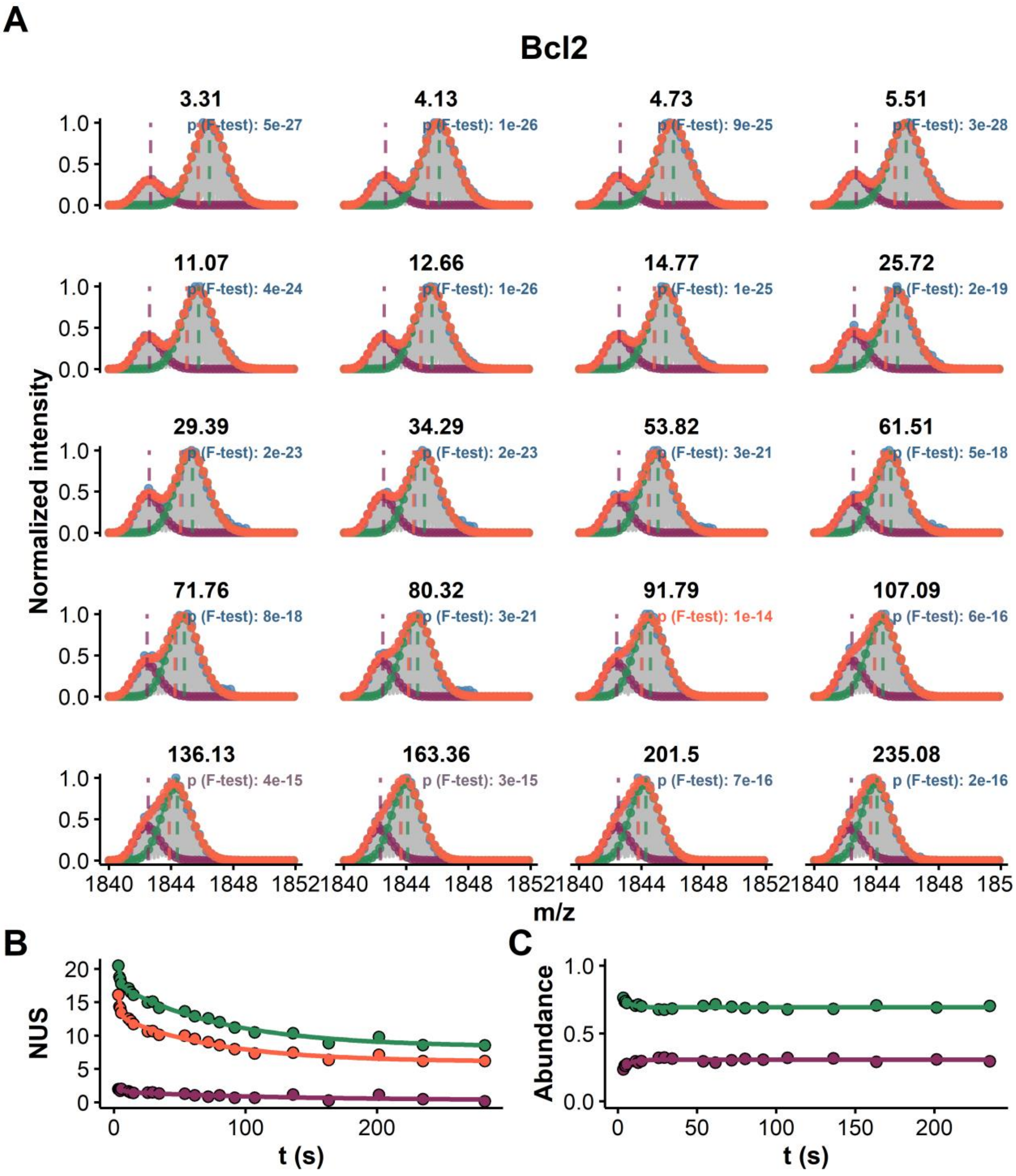


*Figure S34:* *HDX data modeling and bimodal distribution deconvolution of the continuous-flow exchange data of BCl2 (z = 4-, K = 2, initial DC = 90%, final DC = 9%). A. The experimental data is shown in grey, the peak picking in blue, the overall fit in orange, and individual population in green and purple. The vertical dashed lines show the position of their respective centroids, the blue one being the apparent experimental value (when not visible: superimposed with the orange line). B. Exchange kinetics of the overall (orange) and deconvoluted distributions (green and purple). C. Relative abundance of the deconvoluted distributions as a function of the exchange time.*

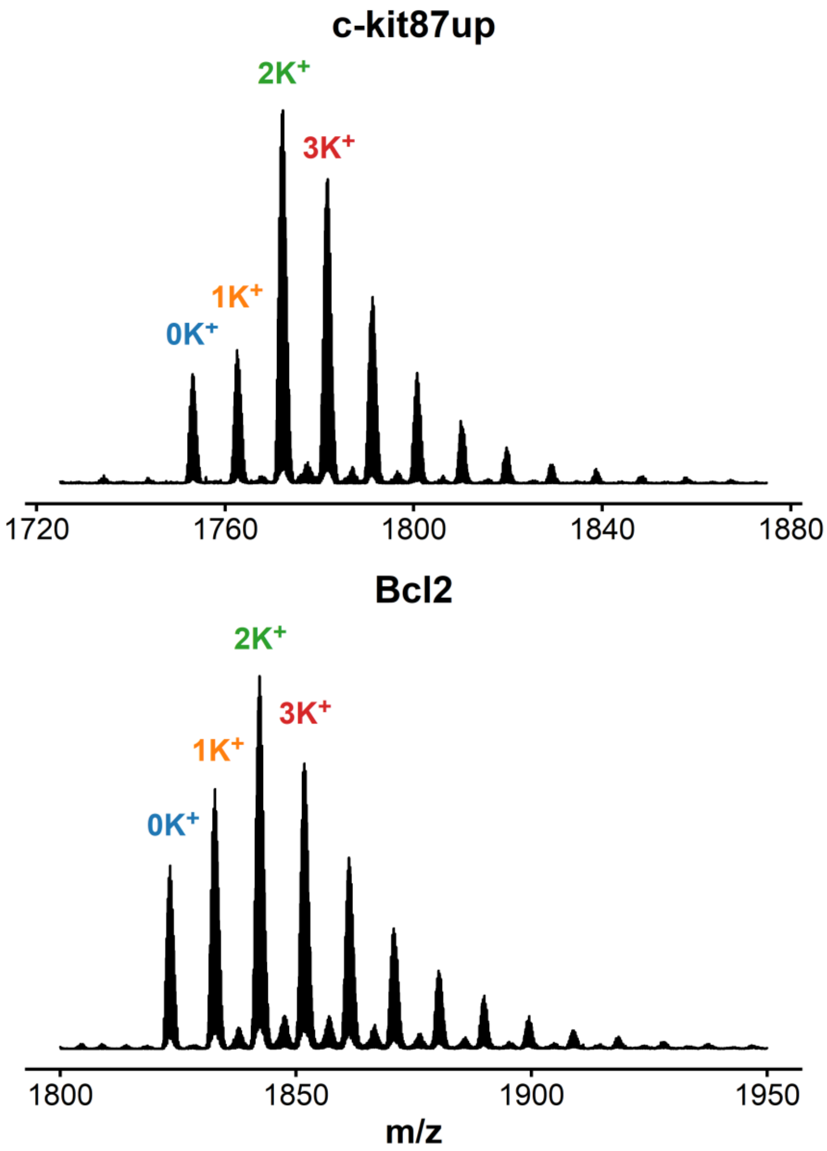


***Figure S35:*** *Native MS spectrum of c-kit87up (top) and Bcl2 (bottom) in 100 mM TMAA, 1 mM KCl (pH = 7.0), zoomed on the 4- charge state.*

#### 24TTG

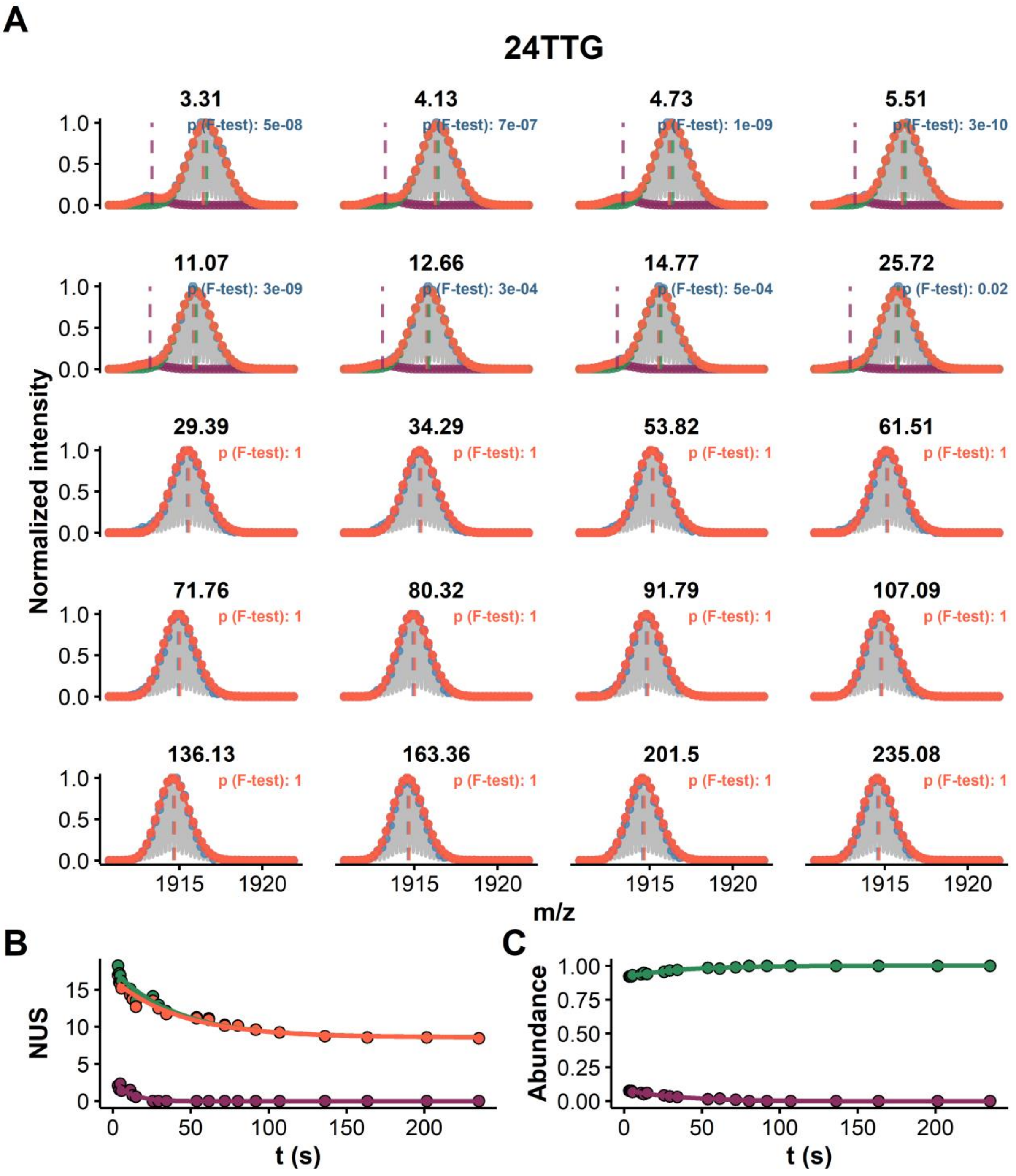


***Figure S36:*** *Bimodal distribution deconvolution on the peak-picked, continuous-flow exchange data of 24TTG (z = 4-, K = 2, initial DC = 90%, final DC = 9%). The experimental data is shown in grey, the peak picking in blue, the overall fit in orange, and individual population in green and purple. The vertical dashed lines show the position of their respective centroids, the blue one being the apparent experimental value (when not visible: superimposed with the orange line)*

#### Mixed EX1/EX2 species

##### 23TAG (2 $K^+$)

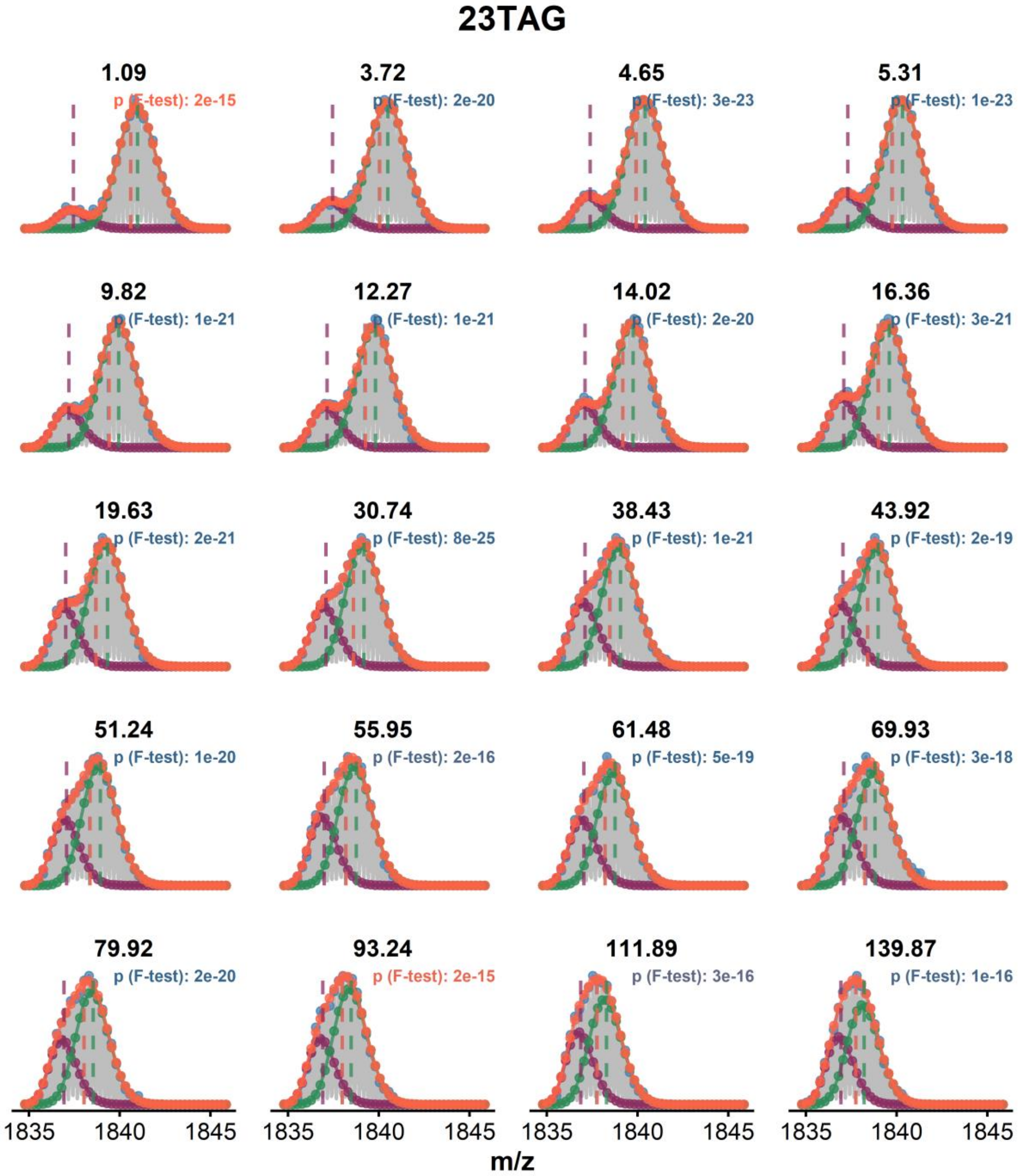


***Figure S37:*** *Bimodal distribution deconvolution on the peak-picked, continuous-flow exchange data of 23TAG (z = 4-, K = 2, initial DC = 90%, final DC = 9%). The experimental data is shown in grey, the peak picking in blue, the overall fit in orange, and individual population in green and purple. The vertical dashed lines show the position of their respective centroids, the blue one being the apparent experimental value (when not visible: superimposed with the orange line)*

#### 23TAG (1 $K^+$)

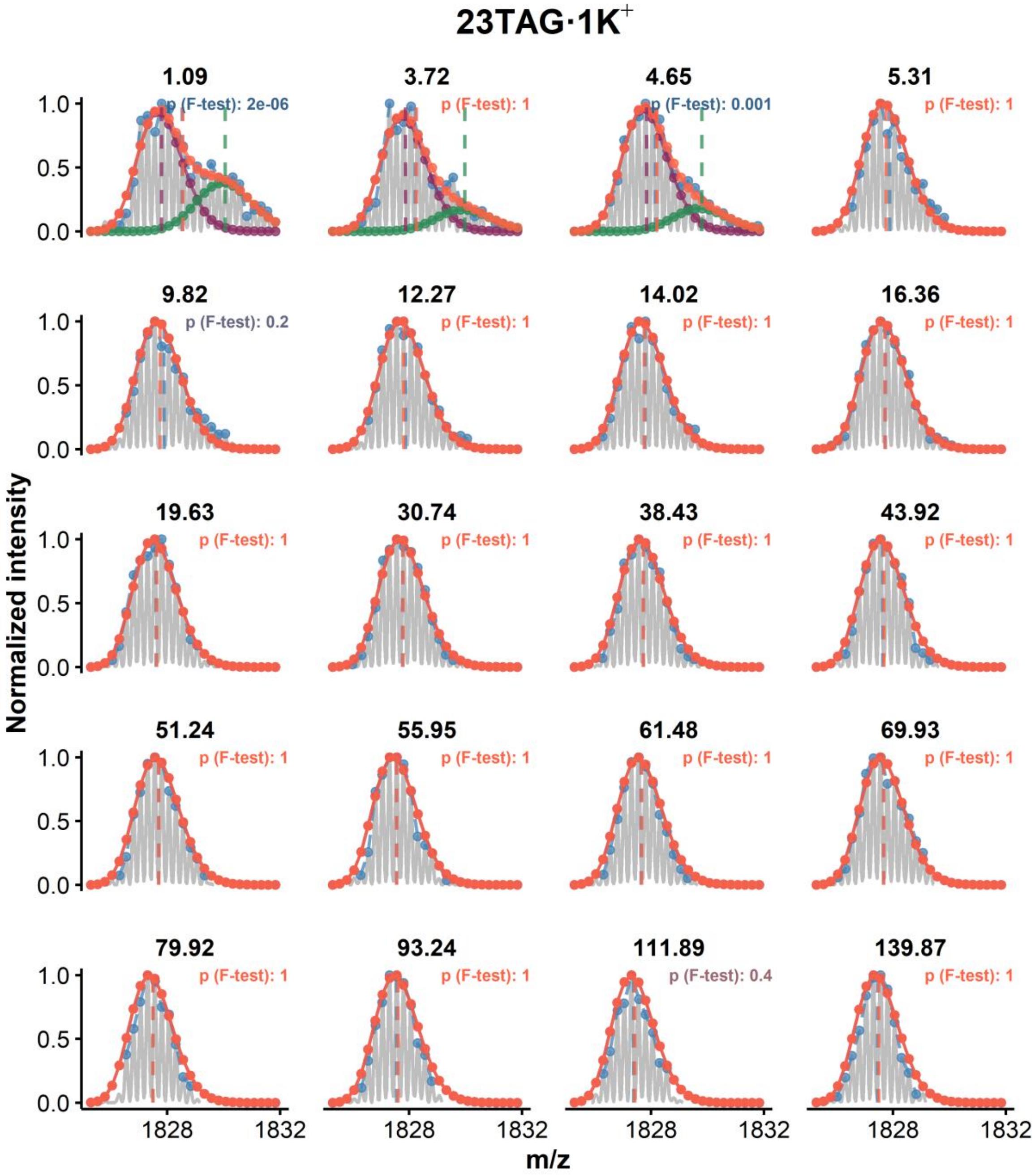


***Figure S38:*** *HDX data modeling and bimodal distribution deconvolution of the continuous-flow exchange data of 23TAG (z = 4-, K = 1, initial DC = 90%, final DC = 9%). The experimental data is shown in grey, the peak picking in blue, the overall fit in orange, and individual population in green and purple. The vertical dashed lines show the position of their respective centroids, the blue one being the apparent experimental value (when not visible: superimposed with the orange line).*

## 22AG (2 $K^+$)

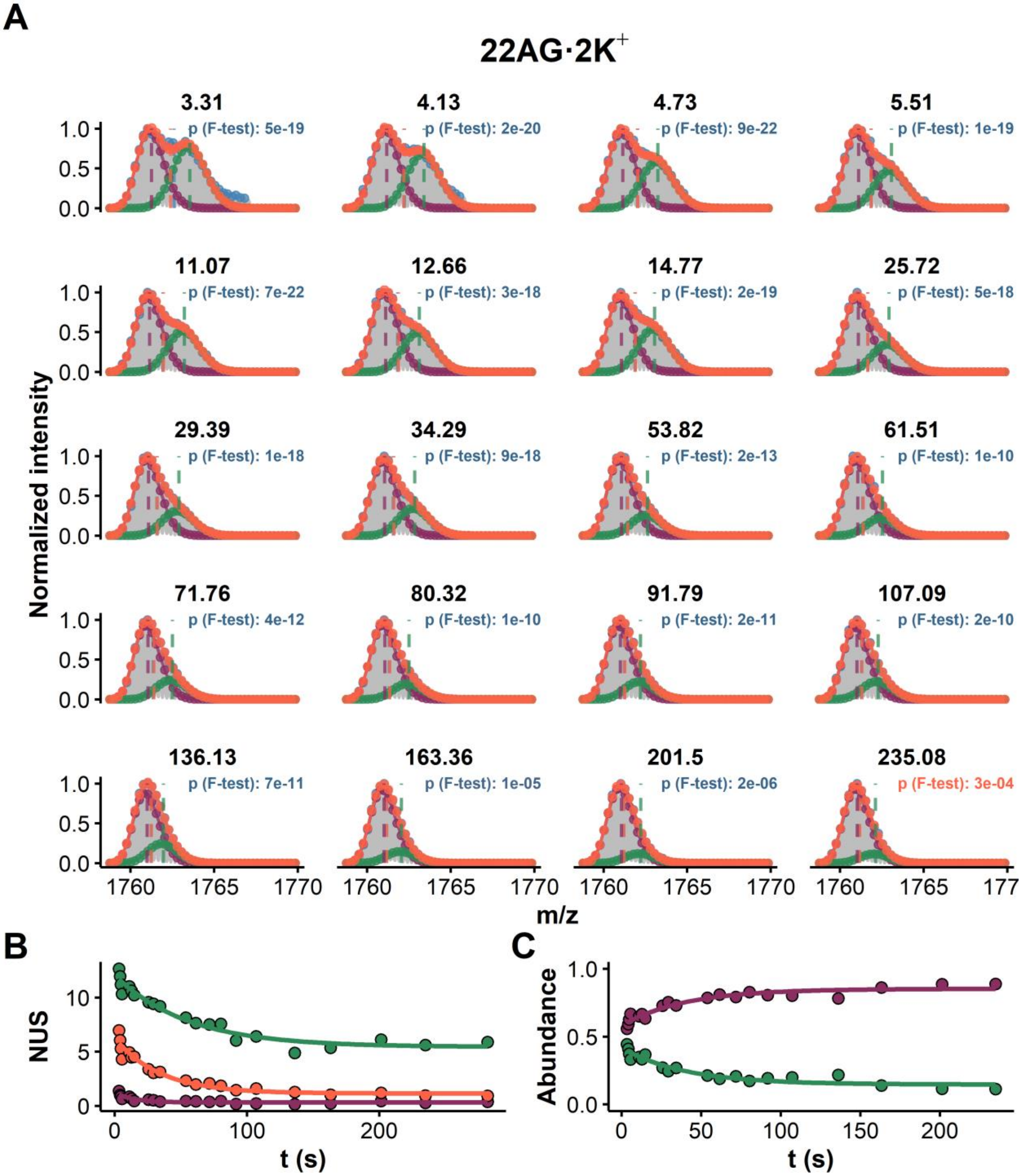


*__Figure S39:__ HDX data modeling and bimodal distribution deconvolution of the continuous-flow exchange data of 22AG (z = 4-, K = 2, initial DC = 90%, final DC = 9%). A. The experimental data is shown in grey, the peak picking in blue, the overall fit in orange, and individual population in green and purple. The vertical dashed lines show the position of their respective centroids, the blue one being the apparent experimental value (when not visible: superimposed with the orange line). B. Exchange kinetics of the overall (orange) and deconvoluted distributions (green and purple). C. Relative abundance of the deconvoluted distributions as a function of the exchange time.*

## 22AG (1 K+)

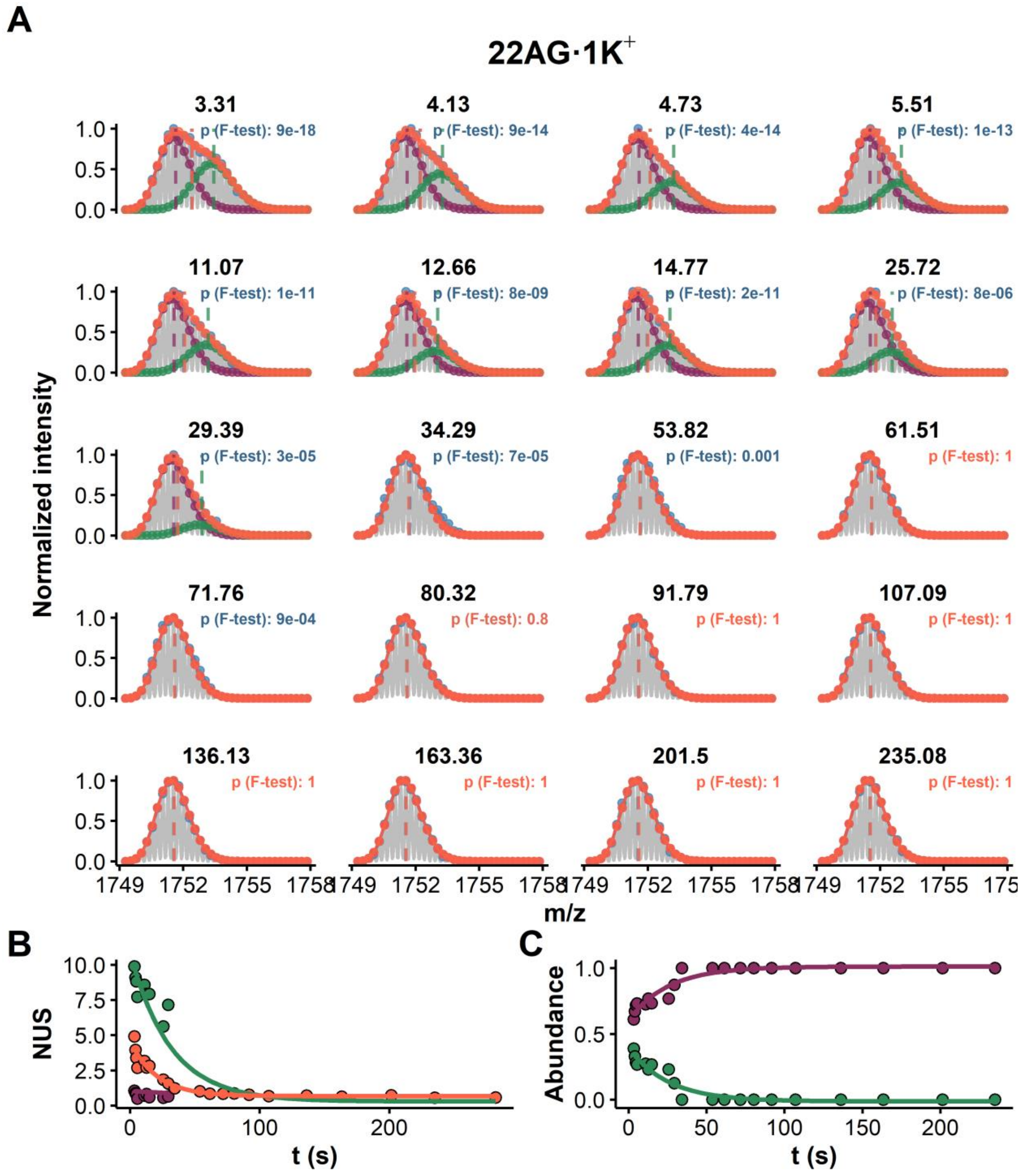


*Figure S40: HDX data modeling and bimodal distribution deconvolution of the continuous-flow exchange data of 22AG (z = 4-, K = 1, initial DC = 90%, final DC = 9%). A. The experimental data is shown in grey, the peak picking in blue, the overall fit in orange, and individual population in green and purple. The vertical dashed lines show the position of their respective centroids, the blue one being the apparent experimental value (when not visible: superimposed with the orange line). B. Exchange kinetics of the overall (orange) and deconvoluted distributions (green and purple). C. Relative abundance of the deconvoluted distributions as a function of the exchange time.*

#### 26TTA (2 $K^+$)

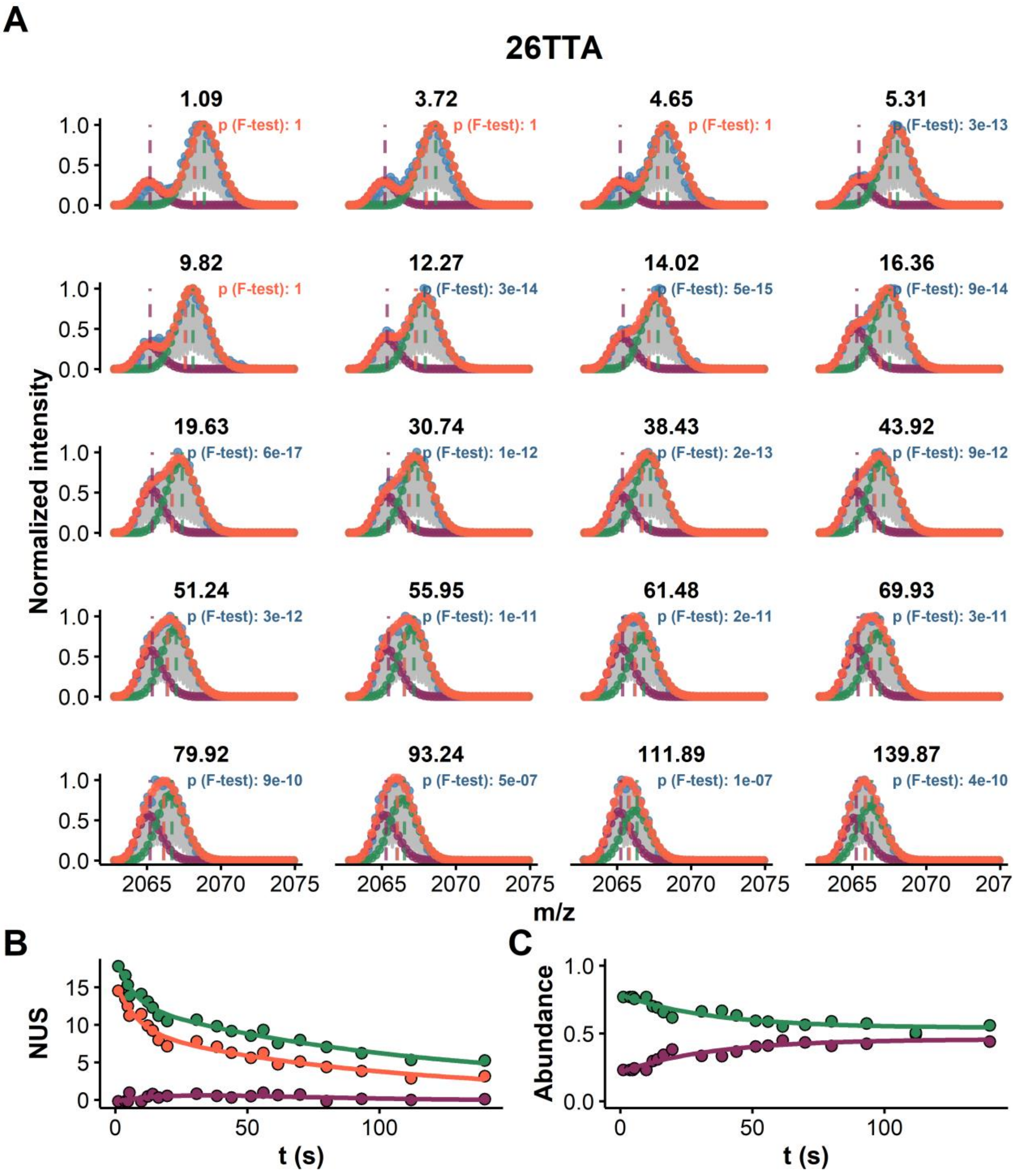


*__Figure S41:__ HDX data modeling and bimodal distribution deconvolution of the continuous-flow exchange data of 26TTA (z = 4-, K = 2, initial DC = 90%, final DC = 9%). A. The experimental data is shown in grey, the peak picking in blue, the overall fit in orange, and individual population in green and purple. The vertical dashed lines show the position of their respective centroids, the blue one being the apparent experimental value (when not visible: superimposed with the orange line). B. Exchange kinetics of the overall (orange) and deconvoluted distributions (green and purple). C. Relative abundance of the deconvoluted distributions as a function of the exchange time.*

#### 26TTA (1 $K^+$)

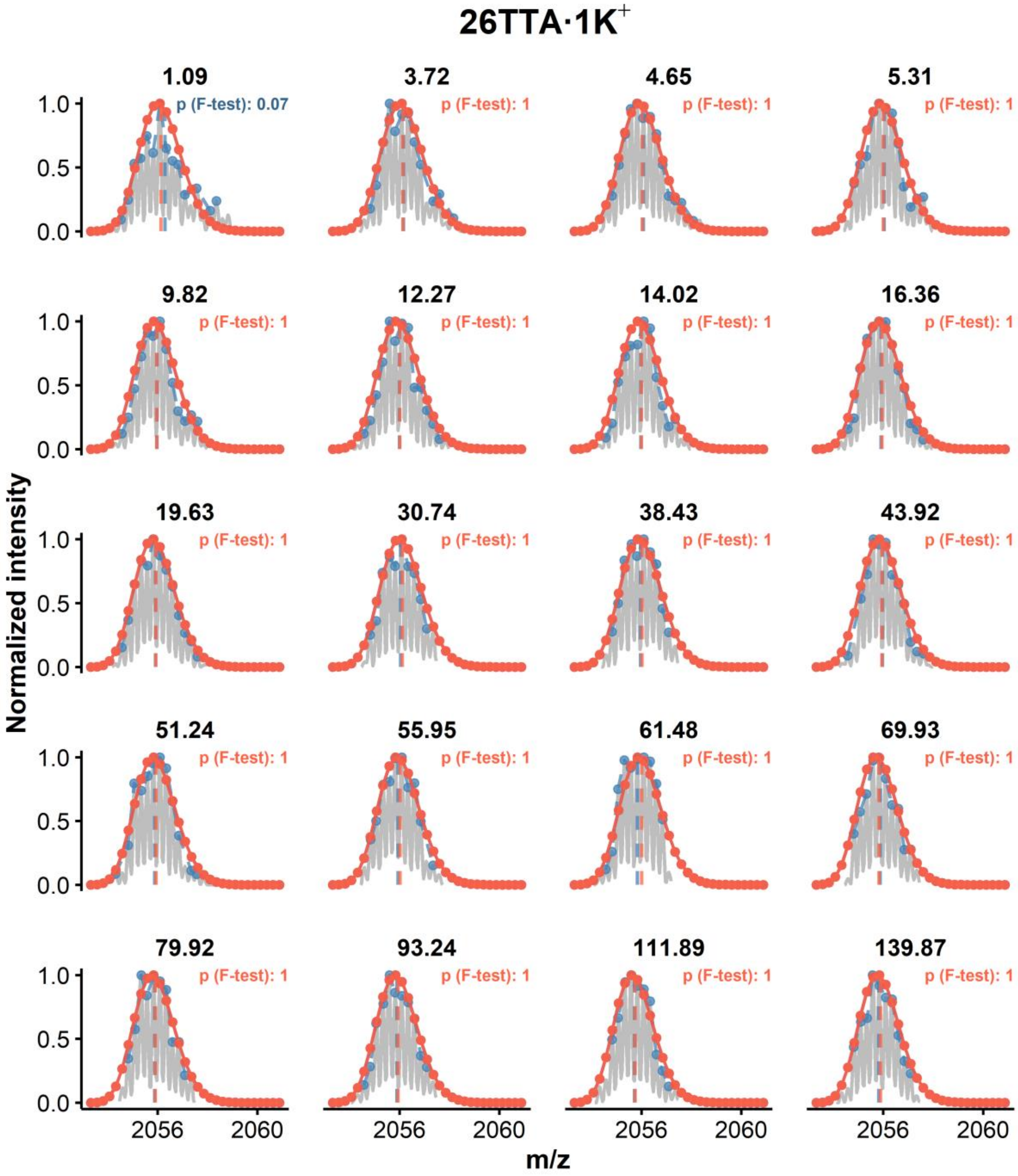


***Figure S42:*** *HDX data modeling and bimodal distribution deconvolution of the continuous-flow exchange data of 26TTA (z = 4-, K = 1, initial DC = 90%, final DC = 9%). The experimental data is shown in grey, the peak picking in blue, the overall fit in orange, and individual population in green and purple. The vertical dashed lines show the position of their respective centroids, the blue one being the apparent experimental value (when not visible: superimposed with the orange line). It is likely that this species exchange with a mixed EX1/EX2 mechanism, however only a couple deuterons remain unexchanged at the shortest exchange time, and the data quality is poor (low species abundance), which ultimately did not yield a satisfactory deconvolution.*

## 222T (2 $K^+$)

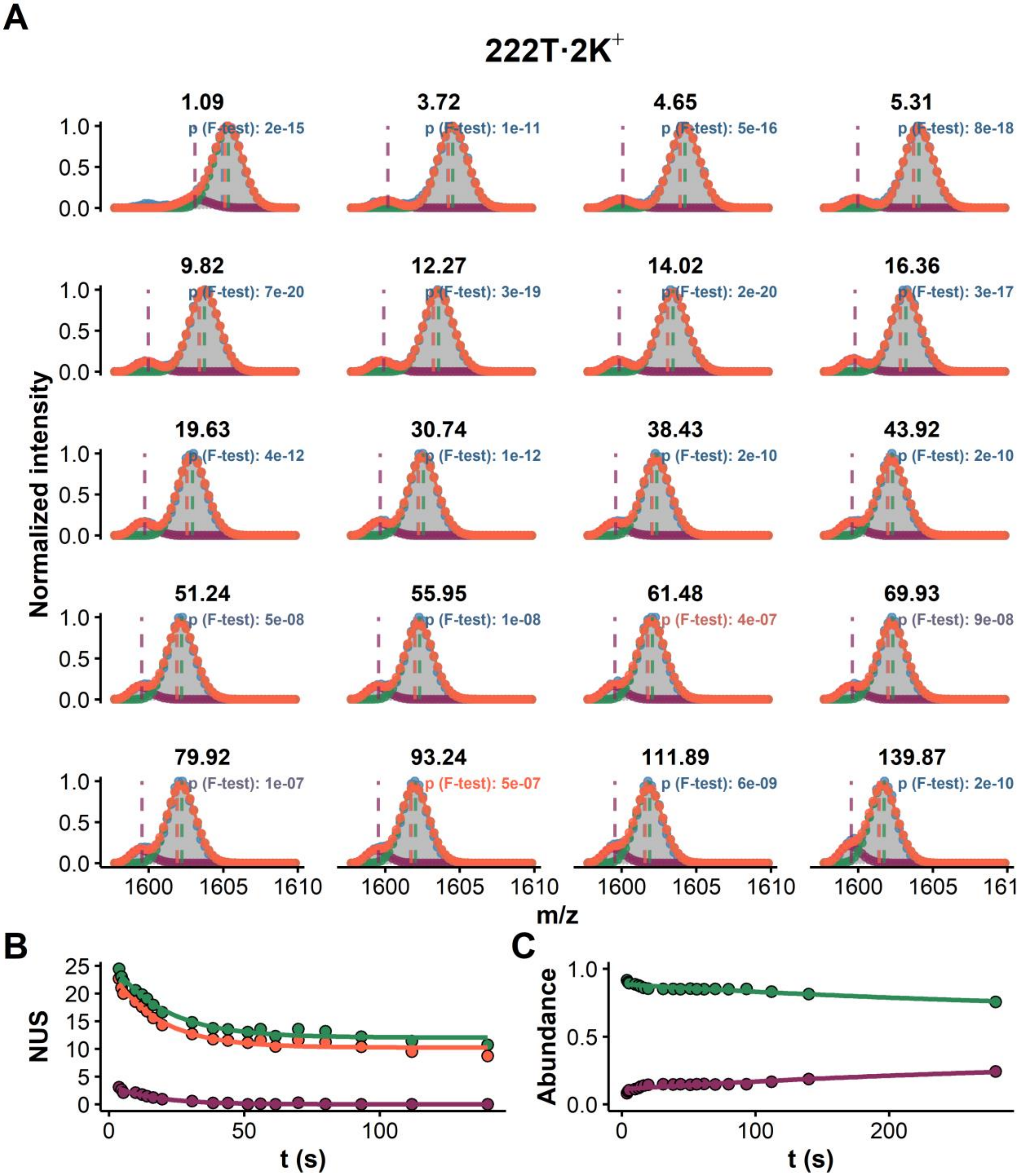


*Figure S43: HDX data modeling and bimodal distribution deconvolution of the continuous-flow exchange data of 222T (z = 4-, K = 2, initial DC = 90%, final DC = 9%). A. The experimental data is shown in grey, the peak picking in blue, the overall fit in orange, and individual population in green and purple. The vertical dashed lines show the position of their respective centroids, the blue one being the apparent experimental value (when not visible: superimposed with the orange line). B. Exchange kinetics of the overall (orange) and deconvoluted distributions (green and purple) (the first point was excluded). C. Relative abundance of the deconvoluted distributions as a function of the exchange time (the first point was excluded).*

#### 222T (2K$^+$) in 0.1 mM KCl

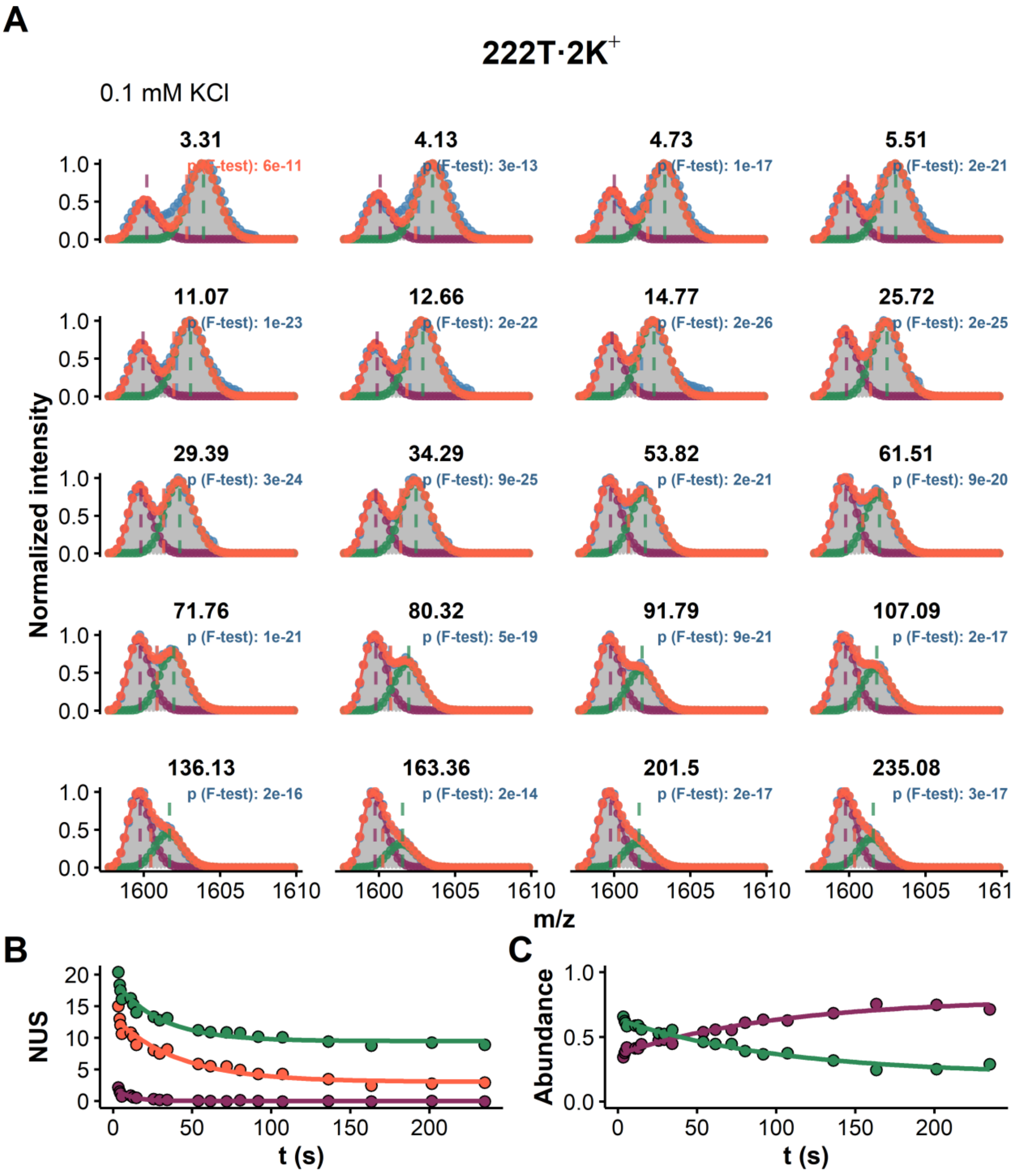


*Figure S44:* *HDX data modeling and bimodal distribution deconvolution of the continuous-flow exchange data of 222T (z = 4-, K = 1, initial DC = 90%, final DC = 9%). A. The experimental data is shown in grey, the peak picking in blue, the overall fit in orange, and individual population in green and purple. The vertical dashed lines show the position of their respective centroids, the blue one being the apparent experimental value (when not visible: superimposed with the orange line). B. Exchange kinetics of the overall (orange) and deconvoluted distributions (green and purple). C. Relative abundance of the deconvoluted distributions as a function of the exchange time.*

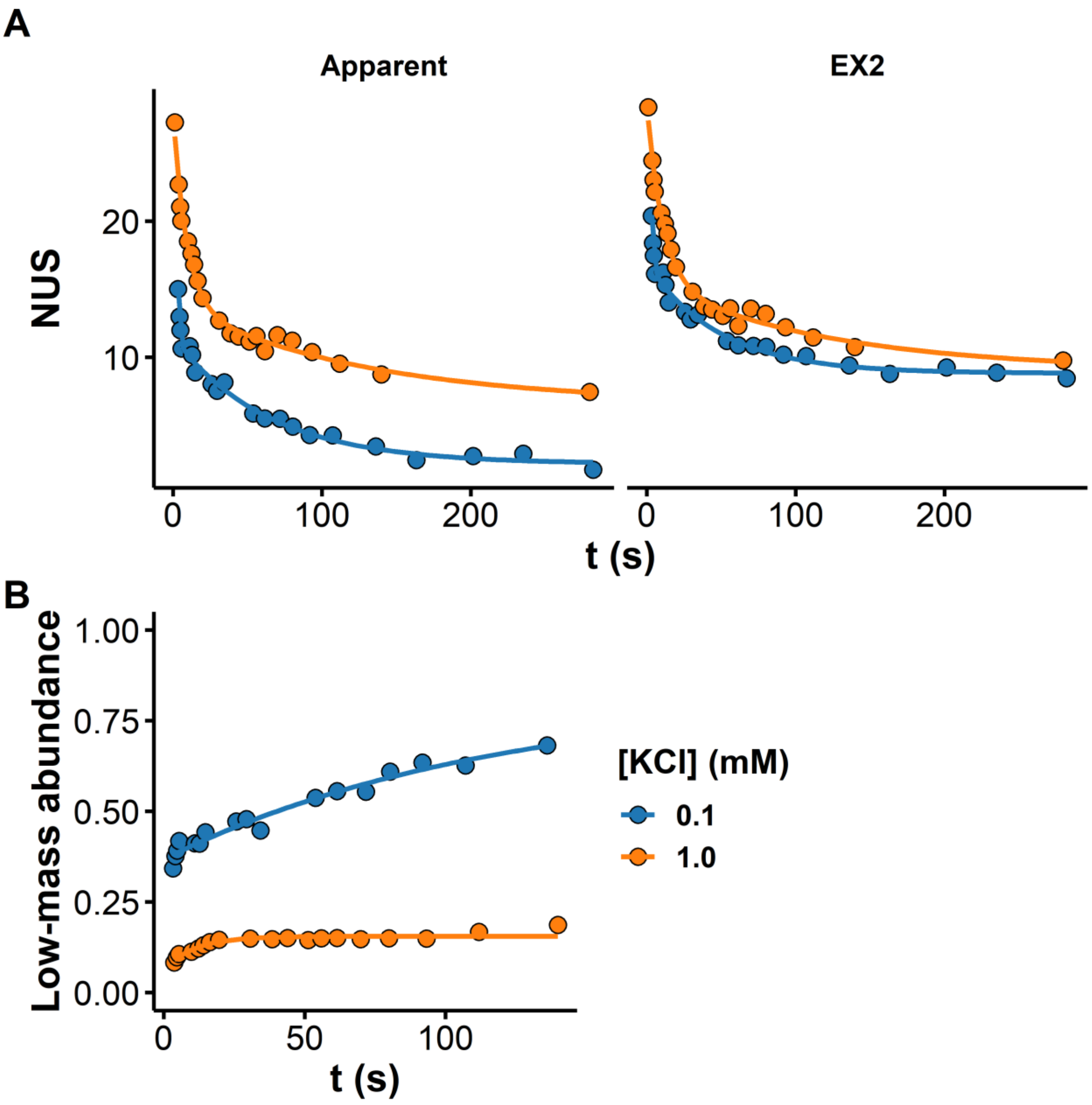


***Figure S45:*** *A. Apparent exchange (left) and EX2-only contribution (right) of 222T in 1.0 or 0.1 mM KCl solutions, B. Relative abundance of the low-mass (high-exchange) population*

## 222T (1 K⁺)

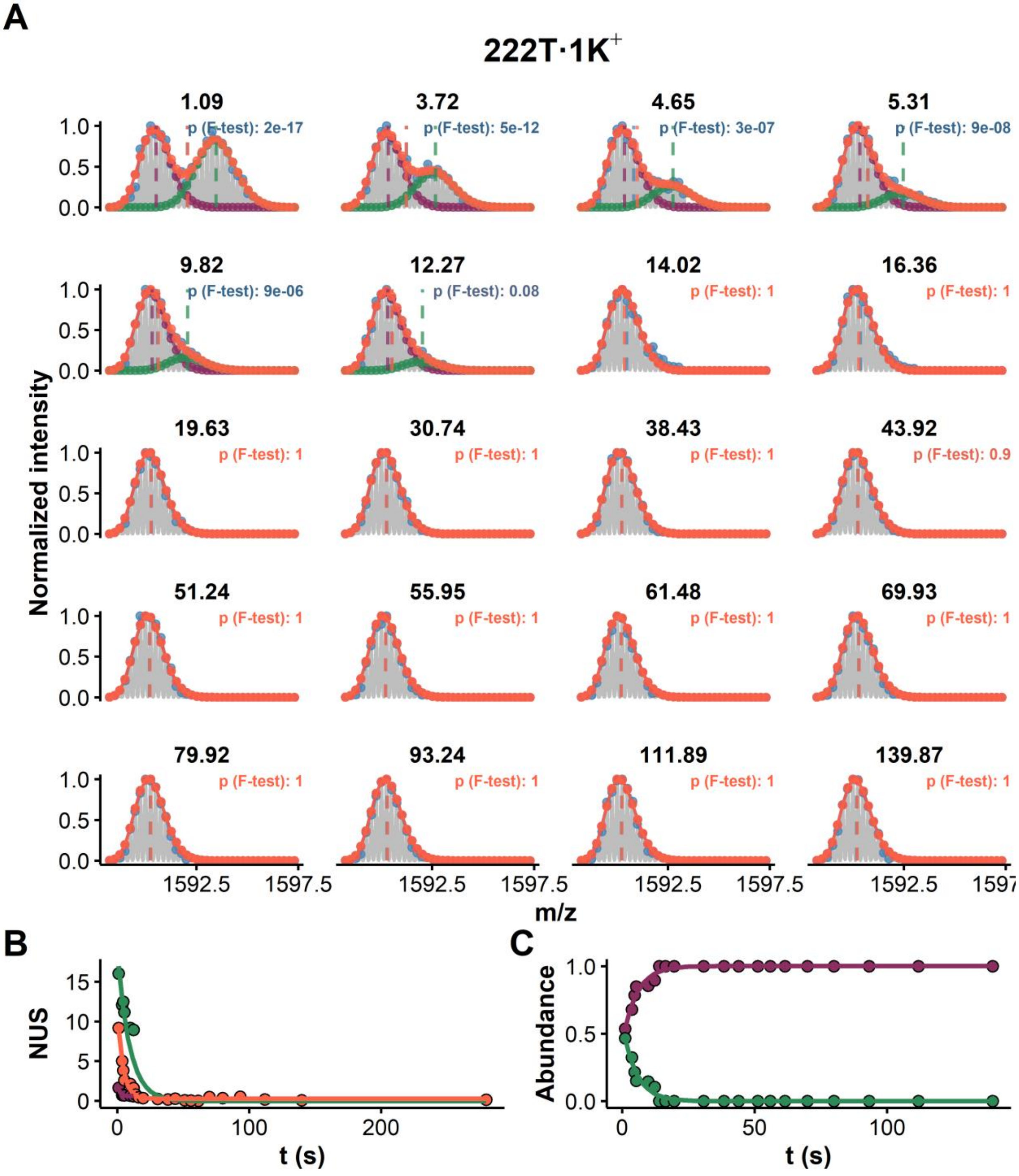


*Figure S46: HDX data modeling and bimodal distribution deconvolution of the continuous-flow exchange data of 222T (z = 4-, K = 1, initial DC = 90%, final DC = 9%). A. The experimental data is shown in grey, the peak picking in blue, the overall fit in orange, and individual population in green and purple. The vertical dashed lines show the position of their respective centroids, the blue one being the apparent experimental value (when not visible: superimposed with the orange line). B. Exchange kinetics of the overall (orange) and deconvoluted distributions (green and purple). C. Relative abundance of the deconvoluted distributions as a function of the exchange time (the first point was excluded).*

#### 222T (1K+) in 0.1 mM KCl

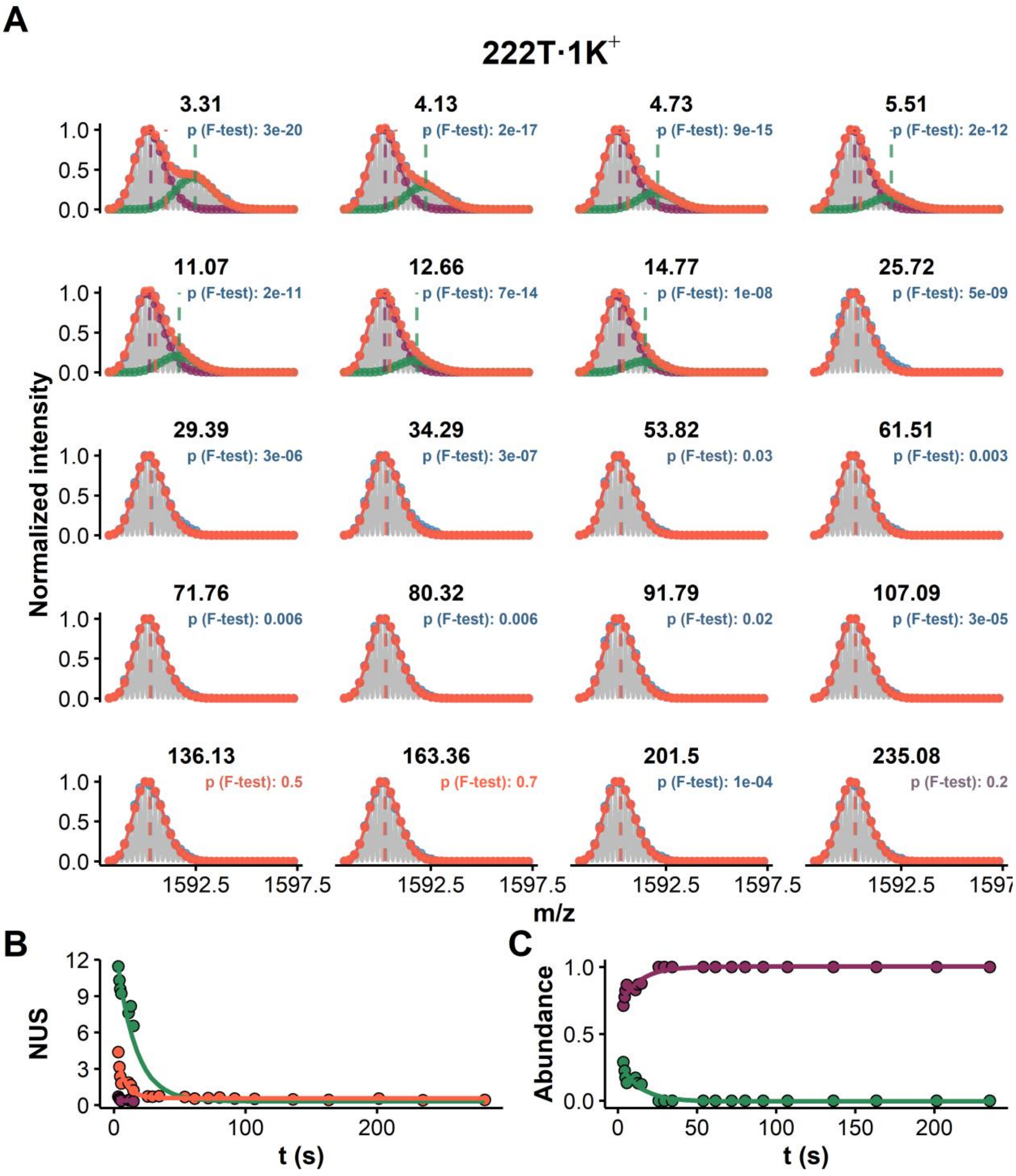


*Figure S47:* *HDX data modeling and bimodal distribution deconvolution of the continuous-flow exchange data of 222T (z = 4-, K = 1, initial DC = 90%, final DC = 9%). A. The experimental data is shown in grey, the peak picking in blue, the overall fit in orange, and individual population in green and purple. The vertical dashed lines show the position of their respective centroids, the blue one being the apparent experimental value (when not visible: superimposed with the orange line). B. Exchange kinetics of the overall (orange) and deconvoluted distributions (green and purple). C. Relative abundance of the deconvoluted distributions as a function of the exchange time.*

#### TBA

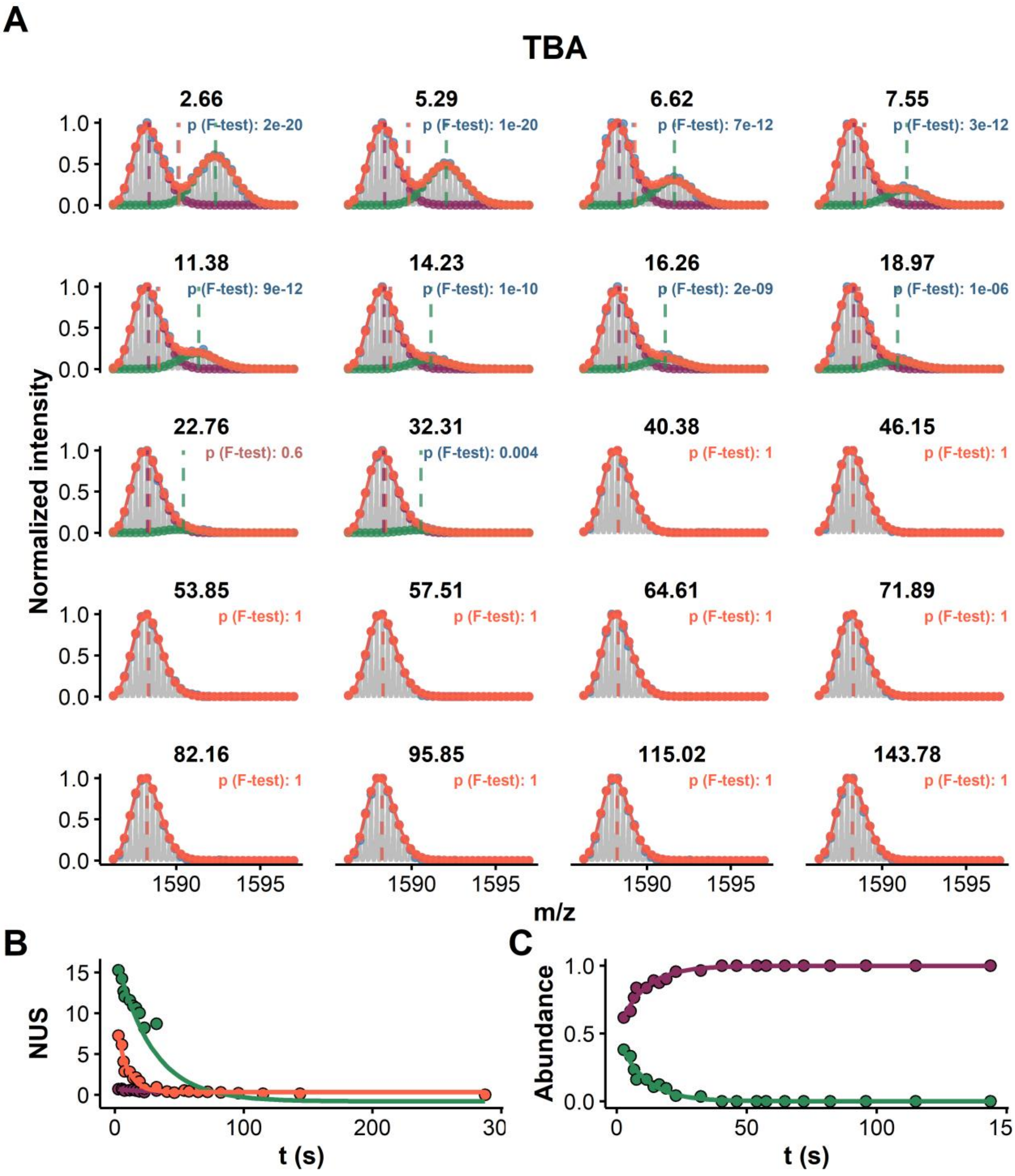


*__Figure S48:__ HDX data modeling and bimodal distribution deconvolution of the continuous-flow exchange data of TBA (z = 3-, K = 1, initial DC = 90%, final DC = 9%). A. The experimental data is shown in grey, the peak picking in blue, the overall fit in orange, and individual population in green and purple. The vertical dashed lines show the position of their respective centroids, the blue one being the apparent experimental value (when not visible: superimposed with the orange line). B. Exchange kinetics of the overall (orange) and deconvoluted distributions (green and purple). C. Relative abundance of the deconvoluted distributions as a function of the exchange time.*

#### 22CTA

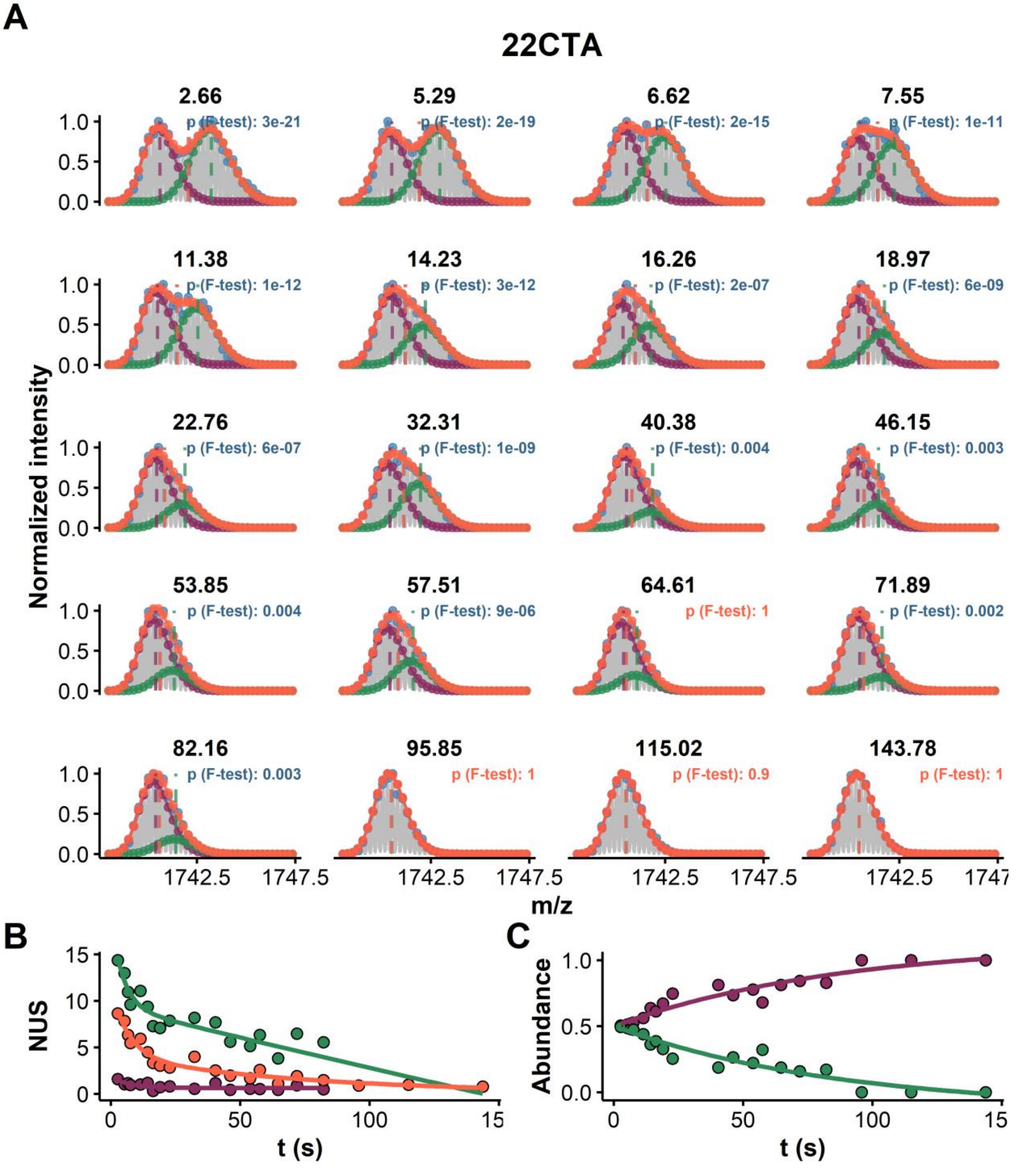


*Figure S49:* *HDX data modeling and bimodal distribution deconvolution of the continuous-flow exchange data of 22CTA (z = 4-, K = 1, initial DC = 90%, final DC = 9%). A. The experimental data is shown in grey, the peak picking in blue, the overall fit in orange, and individual population in green and purple. The vertical dashed lines show the position of their respective centroids, the blue one being the apparent experimental value (when not visible: superimposed with the orange line). B. Exchange kinetics of the overall (orange) and deconvoluted distributions (green and purple). C. Relative abundance of the deconvoluted distributions as a function of the exchange time.*

#### Non-linear fitting of the deconvoluted EX1 contribution

***Table S7:*** *Estimates from the non-linear fitting of the EX1 contributions in mixed EX1/EX2 exchange (i.e. population abundance as a function of time). The high and low mass populations yield the same rates and amplitudes (the sum of the relative abundance is always 1).*

| | Estimates | | | | Standard Errors | | | Significance (Pr(>\|t\|)) | | |
|---|---|---|---|---|---|---|---|---|---|---|
| **Species** | $ab_\infty$ | $ab_\Delta$ | $k_{op}(s^-1)$ | $t_{1/2}(s)$ | $ab_\infty$ | $ab_\Delta$ | $k_{op}(s^-1)$ | $ab_\infty$ | $ab_\Delta$ | $k_{op}(s^-1)$ |
| 23TAG•2K | 0.62 | 0.24 | 0.054 | 12.8 | 0.01 | 0.01 | 0.006 | *** | *** | *** |
| 22AG•2K | 0.15 | 0.27 | 0.025 | 27.7 | 0.02 | 0.02 | 0.006 | *** | *** | *** |
| 22AG•1K | -0.01 | 0.41 | 0.040 | 17.5 | 0.02 | 0.03 | 0.008 | | *** | *** |
| 26TTA•2K | 0.54 | 0.24 | 0.029 | 23.9 | 0.02 | 0.02 | 0.008 | *** | *** | ** |
| 222T•2K | 0.65 | 0.24 | 0.003 | 259.5 | 0.15 | 0.15 | 0.002 | *** | | |
| 222T (0.1 mM KCl) | 0.21 | 0.43 | 0.010 | 70.5 | 0.03 | 0.03 | 0.002 | *** | *** | *** |
| 222T•1K | -0.00 | 0.57 | 0.188 | 3.7 | 0.01 | 0.04 | 0.021 | | *** | *** |
| 222T•1K (0.1 mM KCl | 0.14 | 4.07 | 0.987 | 0.7 | 0.01 | 4.87 | 0.355 | *** | | * |
| TBA | 0.00 | 0.49 | 0.101 | 6.9 | 0.01 | 0.03 | 0.011 | | *** | *** |
| 22CTA | -0.13 | 0.63 | 0.012 | 60.3 | 0.16 | 0.14 | 0.005 | | *** | * |

#### Non-linear fitting of the deconvoluted EX2 contribution

***Table S8:*** *Estimates from the non-linear fitting of the deconvoluted EX2 contributions (using the high-mass data).*

| | Estimates | | | | | | | Standard Errors | | | | | Significance (Pr(>\|t\|)) | | | | |
|---|---|---|---|---|---|---|---|---|---|---|---|---|---|---|---|---|---|
| oligo | $ab_\infty$ | $ab_{\Delta1}$ | $k_1$(s$^{-1}$) | $t_{1/21}$(s) | $ab_{\Delta2}$ | $k_2$(s$^{-1}$) | $t_{1/22}$(s) | $ab_\infty$ | $ab_{\Delta1}$ | $k_1$(s$^{-1}$) | $ab_{\Delta2}$ | $k_2$(s$^{-1}$) | $ab_\infty$ | $ab_{\Delta1}$ | $k_1$(s$^{-1}$) | $ab_{\Delta2}$ | $k_2$(s$^{-1}$) |
| 23TAG | 4.47 | 7.18 | 0.125 | 5.5 | 9.47 | 0.011 | 64.0 | 0.47 | 0.58 | 0.020 | 0.51 | 0.002 | *** | *** | *** | *** | *** |
| 22AG·2K | 5.26 | 6.78 | 0.017 | 40.1 | | | | 0.40 | 0.40 | 0.003 | NA | NA | *** | *** | *** | | |
| 22AG·1K | 0.24 | 10.57 | 0.032 | 21.8 | | | | 0.53 | 0.78 | 0.007 | NA | NA | | *** | *** | | |
| 26TTA·2K | 1.92 | 5.98 | 0.158 | 4.4 | 11.02 | 0.009 | 73.7 | 3.92 | 1.27 | 0.072 | 2.86 | 0.007 | | *** | * | ** | |
| 222T·2K | 12.06 | 14.17 | 0.052 | 13.4 | | | | 0.29 | 0.59 | 0.005 | NA | NA | *** | *** | *** | | |
| 222T·2K (0.1 mM KCl) | 8.99 | 84.25 | 0.920 | 0.8 | 8.01 | 0.022 | 31.9 | 0.27 | 72.77 | 0.264 | 0.43 | 0.003 | *** | | ** | *** | *** |
| 222T·1K | -0.07 | 19.18 | 0.108 | 6.4 | | | | 0.54 | 1.74 | 0.018 | NA | NA | | *** | *** | | |
| 222T·1K (0.1 mM KCl) | 0.31 | 13.85 | 0.069 | 10.0 | | | | 0.28 | 0.91 | 0.010 | NA | NA | | *** | *** | | |
| TBA | -0.80 | 18.29 | 0.036 | 19.3 | | | | 0.79 | 1.14 | 0.006 | NA | NA | | *** | *** | | |
| 22CTA | -0.97 | 13.24 | 0.013 | 51.6 | | | | 3.22 | 2.86 | 0.007 | NA | NA | | *** | - | | |
| c-kit87up | 12.69 | 14.74 | 0.360 | 1.9 | 11.64 | 0.012 | 58.1 | 0.72 | 7.20 | 0.142 | 0.63 | 0.002 | *** | - | * | *** | *** |
| Bcl2 | 8.29 | 40.66 | 0.798 | 0.9 | 9.64 | 0.012 | 56.1 | 0.43 | 34.23 | 0.249 | 0.38 | 0.002 | *** | | ** | *** | *** |
| 24TTG | 8.18 | 12.17 | 0.469 | 1.5 | 7.89 | 0.018 | 39.5 | 0.28 | 7.77 | 0.186 | 0.37 | 0.002 | *** | | * | *** | *** |

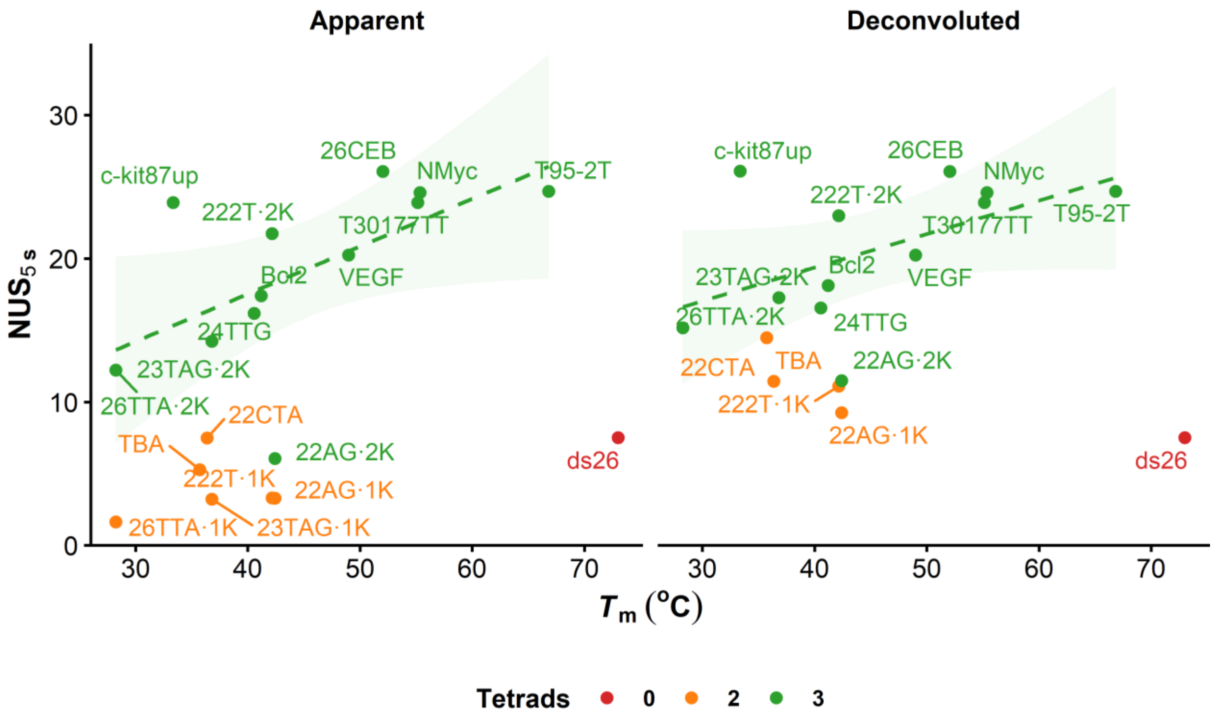


*Figure S50: Relationship between melting temperatures and deconvoluted NUS values: left. Non-deconvoluted data, for reference, right: data extracted from the pure EX2 plots.*

#### Promotion of EX1 by destabilisation of G4s

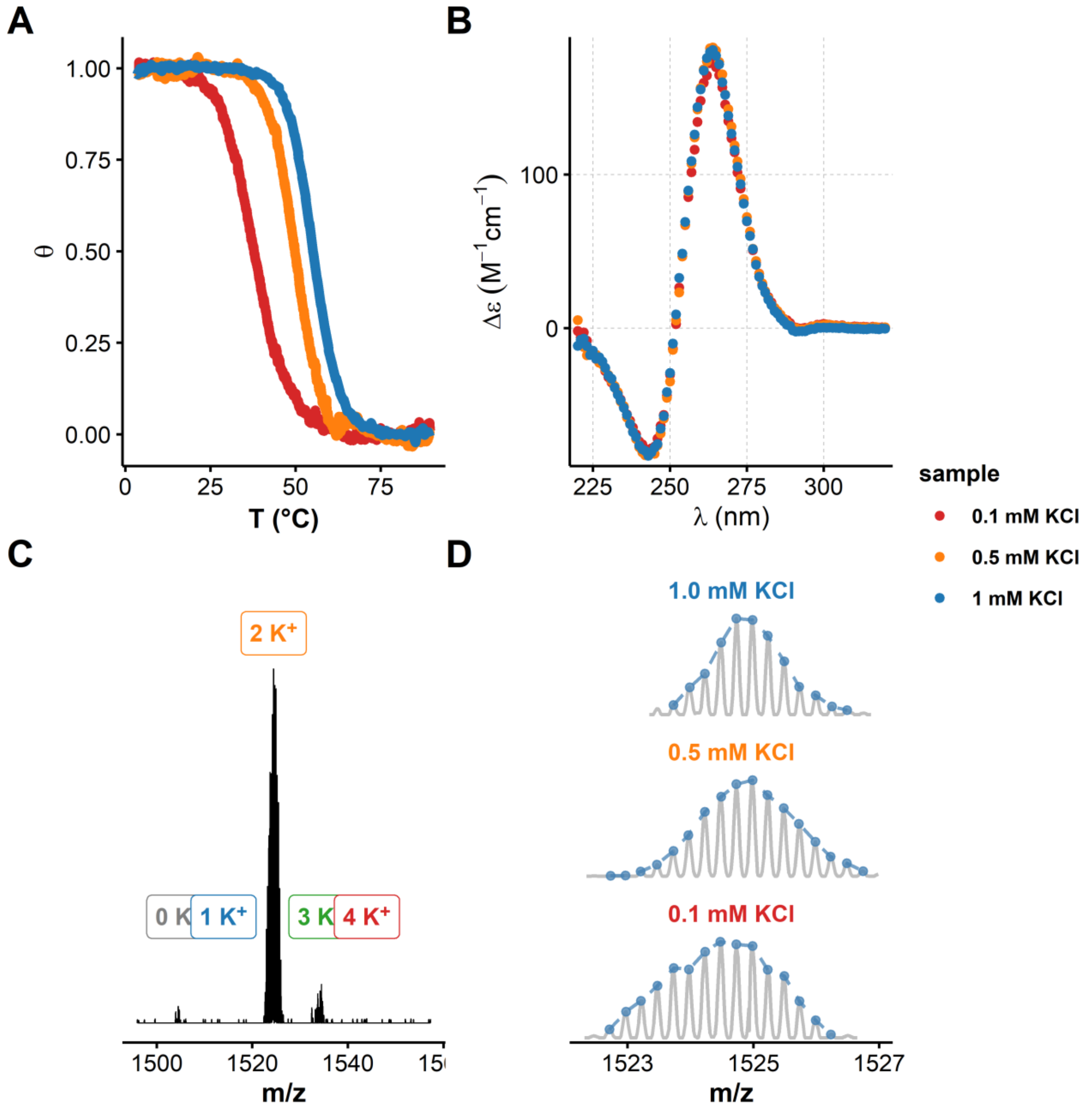


***Figure S51:*** *Promotion of EX1 exchange kinetics of a parallel G4. A. Processed UV-melting data (folded fraction as a function of the solution temperature; cooling ramp) showing the destabilisation of T30177TT upon [KCl] decrease, B. Circular dichroism spectra, showing the parallel folding is quantitatively maintained when decreasing [KCl] to 0.1 mM. C. Native HDX/MS spectrum of T30177-TT (4- charge state) from a 0.1 mM KCl solution confirming the absence of detectable unfolded species (no 0 and 1 $K^+$ complex) at equilibrium. D. Isotopic distributions of T30177-TT·$2K^+$ (4- charge state) after 12 min of exchange, in solutions containing 1.0, 0.5, or 0.1 mM KCl. Isotopic distribution widening is attributed here to the EX1 contribution.*

#### Exchange scenarios

I. Single populated conformer:

1. High $\Delta G_{unf}^{0}$ species: case of an energy landscape with a deep minimum and local exchange *via* microstates. A single isotopic population is detected, e.g., T30177TT.

2. Low $\Delta G_{unf}^{0}$ species: multiple sites are available for exchange *via* partially or fully unfolded states (e.g., triplex, hairpins). Multiple isotopic populations with concerted changes in abundances are detected.

II. Multiple populated conformers:

1. Only high $\Delta G_{unf}^{0}$ species: Multiple isotopic populations experiencing EX2 as in I.1., with unchanged abundances due to very slow interconversion rates. This was not observed in our panel.

2. Presence of at least one low $\Delta G_{unf}^{0}$ species: case of an energy landscape with several shallow minima. Each populated conformer exchanges depending on their propensity to unfold/refold into other conformations:

   a. No/little interconversion between populated conformers within HDX time frame: little/no change in isotopic population abundances (as in II.1.):

      i. Isotopic populations are m/z resolved if the conformers have different m/z (e.g., 222T•2$K^+$ and 222T•1$K^+$) or the same m/z but significantly different HDX rates (e.g., VEGF•2$K^+$ and VEGF•1$K^+$ binding an additional $K^+$ non-specifically). In the latter case, the population may only be resolved in a certain exchange time frame.

      ii. Isotopic populations are not m/z resolved if the conformers have the same m/z and similar exchange rates, e.g., the hybrid-1 and hybrid-2 folds of 24TTG•2$K^+$. Coupling to IMS may allow resolution of these populations.

   b. Significant interconversion between populated conformers within HDX time frame: concerted changes in isotopic population abundances.

      i. Via an unpopulated, partially unfolded conformer: partially exchanged second isotopic population, e.g., 23TAG•2$K^+$

      ii. Via a fully unfolded state: completely exchanged second isotopic population.

Note that (*i*) all populated conformers are potentially subject to EX2, regardless of their propensity to unfold, (*ii*) a single sequence can fall into multiple cases, and that (*iii*) solutions conditions affects the exchange behavior of G4s by altering the apparent $k_{HDX}^{EX1}$.
